# Single-cell virus sequencing of influenza infections that trigger innate immunity

**DOI:** 10.1101/437277

**Authors:** Alistair B. Russell, Jacob R. Kowalsky, Jesse D. Bloom

**Affiliations:** Basic Sciences Division and Computational Biology Program, Fred Hutchinson Cancer Research Center, Seattle, WA 98109, USA; Department of Genome Sciences, University of Washington, Seattle, WA 98109, USA; Howard Hughes Medical Institute, Seattle, WA 98109, USA

**Keywords:** PacBio, single-cell RNA-seq, influenza, interferon, NS1, stochastic

## Abstract

The outcome of viral infection is extremely heterogeneous, with infected cells only sometimes activating innate immunity. Here we develop a new approach to assess how the genetic variation inherent in viral populations contributes to this heterogeneity. We do this by determining both the transcriptome and full-length sequences of all viral genes in single influenza-infected cells. Most cells are infected by virions with defects such as amino-acid mutations, internal deletions, or failure to express a gene. We identify instances of each type of defect that increase the likelihood that a cell activates an innate-immune response. However, immune activation remains stochastic in cells infected by virions with these defects, and sometimes occurs even when a cell is infected by a virion that expresses unmutated copies of all genes. Our work shows that viral genetic variation substantially contributes to but does not fully explain the heterogeneity in single influenza-infected cells.

## INTRODUCTION

Infection with an acute virus such as influenza initiates a race between the virus and immune system. As the virus spreads, some cells detect infection and produce interferon (IFN). This IFN directs expression of anti-viral interferon-stimulated genes (ISGs) in the infected cell and its neighbors via autocrine and paracrine signaling, as well as helping launch a broader immune response (Stetson and Medzhitov, 2006; Honda et al., 2006). If innate immunity is activated sufficiently rapidly, it can reduce viral replication and disease (Solov’ev, 1969; Treanor et al., 1987; Beilharz et al., 2007; Kugel et al., 2009; Steel et al., 2010)—although excessive immune responses later in infection can actually be associated with immunopathology and severe disease (La Gruta et al., 2007; Iwasaki and Pillai, 2014).

Unfortunately for the host, influenza only rarely triggers IFN production by infected cells (Kallfass et al., 2013; Killip et al., 2017). This rareness of IFN induction is just one form of the extreme cell-to-cell heterogeneity that characterizes infection: cells also vary widely in their production of viral mRNA, proteins, and progeny virions (Russell et al., 2018; Steuerman et al., 2018; Sjaastad et al., 2018; Heldt et al., 2015). Because viral growth and the IFN response are both feed-forward processes, early cell-to-cell heterogeneity could have significant downstream consequences for the race between virus and immune system— especially since natural human infections are typically initiated by just a few virions entering a few cells (McCrone et al., 2018; Xue and Bloom, 2018; Varble et al., 2014).

It is unclear why only some infected cells trigger innate-immune responses. Two possible contributors are pure stochasticity and pre-existing variation in cellular state. For instance, only some cells induce IFN even upon treatment with synthetic innate-immune ligands (Shalek et al., 2013, 2014; Wimmers et al., 2018), and the frequency of IFN induction may depend on a cell’s pre-existing chromatin state (Bhushal et al., 2017). But for influenza, a third possible contributor looms large: viral genetic diversity. Because influenza has a high mutation rate, individual virions often have defects (Parvin et al., 1986; Suarez et al., 1992; Bloom, 2014; Pauly et al., 2017). Indeed, many studies have identified mutations that increase IFN induction when engineered into a viral population (te Velthuis et al., 2018; Du et al., 2018; Killip et al., 2017; Pérez-Cidoncha et al., 2014), and viral stocks that are rich in internal deletions in the polymerase genes induce more IFN (Baum et al., 2010; Tapia et al., 2013; Boergeling et al., 2015; Dimmock and Easton, 2015).

However, existing techniques are inadequate to determine how viral genetic diversity contributes to cell-to-cell heterogeneity during infection. Flow cytometry and fluorescent reporters only measure protein levels (Brooke et al., 2013; Guo et al., 2017), and current single-cell transcriptomic techniques primarily measure abundance of transcripts and provide only fragmentary information on their sequences (Russell et al., 2018; Zanini et al., 2018a,b; Steuerman et al., 2018; Saikia et al., 2019; O’Neal et al., 2018). None of these techniques reliably reveal if there are mutations in the virion infecting any given single cell.

Here we develop a new approach to determine both the transcriptome and full sequences of all viral genes in single influenza-infected cells. To do this, we perform both standard Illumina-based transcriptomics and full-length PacBio sequencing of viral genes from single cells. We obtain transcriptomes and sequences of all expressed viral genes in 150 infected cells, 40 of which express IFN. Two-thirds of cells are infected by virions with a mutation or defect in gene expression. This viral diversity is a major contributor to cell-to-cell heterogeneity, with cells infected by unmutated virions having a tighter distribution of viral transcriptional burden. We identify several types of viral defects that increase IFN induction. However, viral genetic variation does not fully explain the heterogeneity, and even unmutated virions sometimes induce IFN. Therefore, viral diversity is an important but not exclusive cause of cell-to-cell heterogeneity during influenza infection.

## RESULTS

### A system to identify and enrich rare IFN+ cells

A challenge in studying IFN induction by influenza virus is its rareness at the level of single cells (Killip et al., 2017; Kallfass et al., 2013; Russell et al., 2018). To identify and enrich rare IFN+ cells, we created A549 cells that carried IFN reporters consisting of a type I (*IFNB1*) or type III (*IFNL1*) promoter driving expression of a cell-surface protein (LNGFRΔC; Bonini et al., 1997; Ruggieri et al., 1997) followed by a fluorescent protein (Figure 1A). Cells that activate the reporter can be enriched by magnetic-activated cell sorting (MACS) or identified by flow cytometry. The reporters were efficiently activated by infection with saturating amounts of a strain of Sendai virus (Strahle et al., 2006) that potently induces IFN (Figure S1A), and activation of the type I and type III IFN reporters was highly correlated (Figure S1B; further validated by the single-cell transcriptomics below). For the rest of this paper, we use “IFN expression” to refer to combined expression of type I and III IFNs.

**Figure 1.**
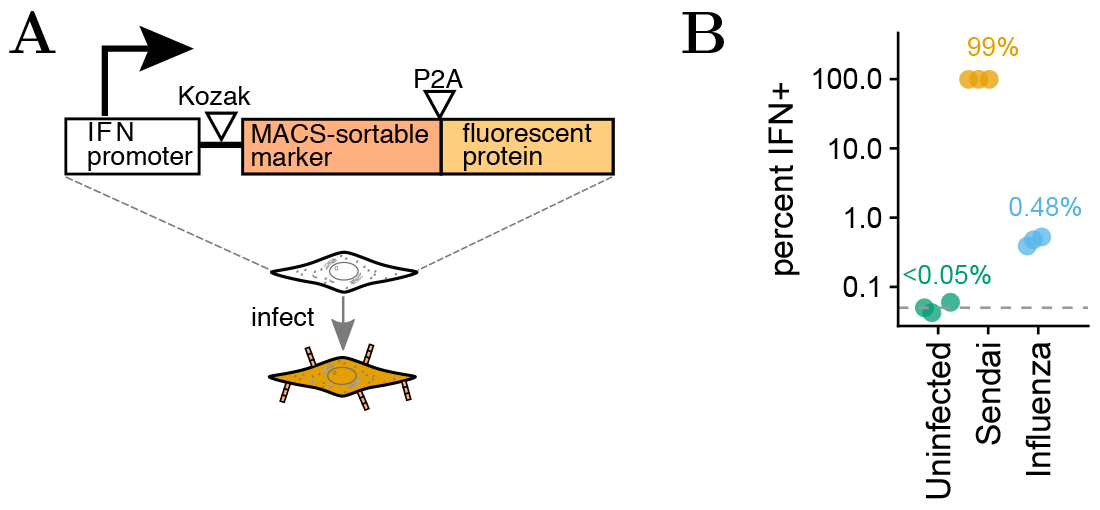
Reporter cells to identify and enrich infections that activate IFN expression. (**A**) The reporter consists of an IFN promoter that drives expression of a cell-surface protein amenable to MACS and a fluorescent protein. We created reporters with type I and type III IFN promoters (File S1). In A549 cells, the reporters were efficiently activated by an IFN-inducing strain of Sendai virus (Figure S1A). (**B**) Frequency of IFN induction upon infection with the influenza virus stock used in the single-cell studies in this paper, as quantified using the type III IFN reporter (see Figure S1C for full data). The plot also shows uninfected cells, and cells infected with saturating amounts of Sendai virus. The limit of detection of 0.05% is indicated with a dashed line, and numbers show the median of three measurements.

We generated a stock of A/WSN/1933 (H1N1) influenza (hereafter referred to as “WSN”), and found that it activated the IFN reporter in ~0.5% of infected cells (Figure 1B), a frequency roughly comparable to that reported by prior studies that have examined IFN induction by influenza in single cells (Killip et al., 2017; Russell et al., 2018; Kallfass et al., 2013).

### Combined transcriptomics and virus-sequencing of single infected cells

To determine if mutations or other defects in the infecting virions contribute to the heterogeneous outcome of infection, we developed the approach in Figure 2 to obtain the entire transcriptome *and* the full sequences of all viral genes in single cells. First, we generated a stock of virus that consisted of a mix of wild-type WSN and a “synonymously barcoded” variant that contained two engineered synonymous mutations near each termini of each gene (File S1). These viral barcodes allow us to identify co-infections, and provide a control for PCR artifacts during full-length sequencing of viral transcripts (see below). We used this viral stock to infect A549 IFN reporter cells (Figure 2A) at a dose that led to detectable viral transcription in about a quarter of cells. From 12 to 13 hours post-infection, we used MACS to enrich cells that activated the IFN reporter (Figure S2). To ensure the presence of IFN-cells, we added back non-enriched cells to ~10% of the total. We also added uninfected canine cells to ~5% of the total as a control for multiplets and to estimate the background amount of viral mRNA detected in truly uninfected cells.

We processed the cells on a commercially available platform (Zheng et al., 2017) that isolates cells in droplets and reverse transcribes polyadenylated mRNAs to append a unique cell barcode to all cDNAs in each droplet, and a unique molecular identifier (UMI) to each cDNA molecule (Figure 2B). Because influenza virus mRNAs are polyadenylated (Robertson et al., 1981), this process appends cell barcodes to both cellular and viral mRNAs. Furthermore, because virtually the entire influenza genome is transcribed, the cell-barcoded cDNA spans almost all 13,581 nucleotides in the segmented viral genome: the only portions not covered are one universally conserved nucleotide upstream of the transcription start site (Koppstein et al., 2015) and 17 to 22 highly conserved nucleotides downstream of the polyadenylation site (Robertson et al., 1981) in each of the eight viral gene segments.

**Figure 2.**
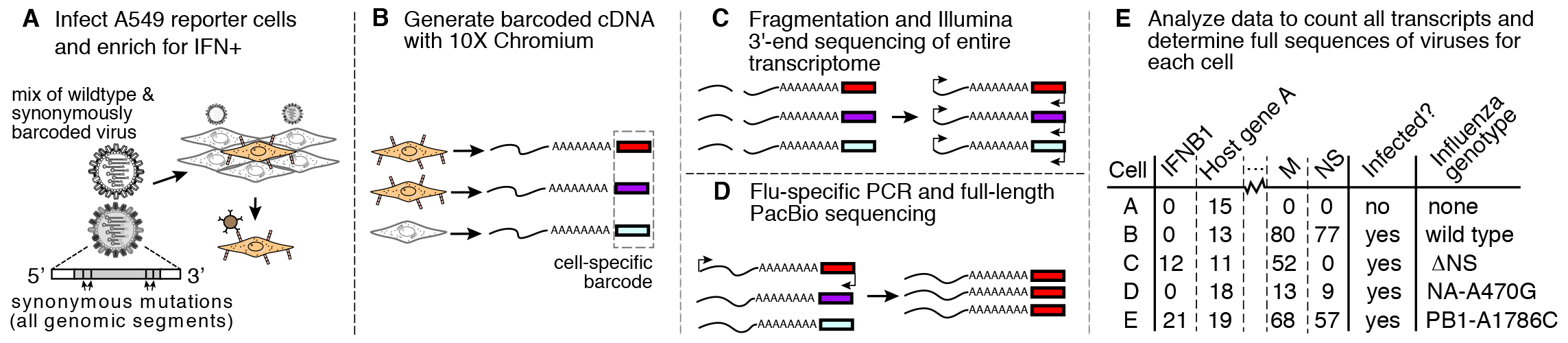
Approach for combined transcriptomics and virus sequencing of single influenza-infected cells that express IFN. (**A**) IFN reporter A549 cells are infected with a mix of wild-type and synonymously barcoded viruses. IFN+ cells are enriched by MACS (Figure S2), and pooled with non-enriched cells and uninfected canine cells that serve as an internal control for multiplets and mRNA leakage. (**B**) The mRNAs from individual cells are converted to cDNAs tagged with cell-specific barcodes. (**C**) Cellular transcriptomes are quantified using standard single-cell 3’-end Illumina sequencing, and (**D**) viral genes are enriched by influenza-specific PCR and fully sequenced by PacBio (in this schematic, only the cell labeled by the red barcode is infected and has viral transcripts that are enriched and sequenced by PacBio). (**E**) The result is a matrix giving the expression of each gene in each cell, as well as the full sequences of the viral genes in infected cells.

We used a portion of the cell-barcoded cDNA for standard single-cell transcriptomics by Illumina 3’-end sequencing (Figure 2C). But we also took a portion and enriched for full-length viral molecules by PCR (Figure 2D). We performed PacBio sequencing on these viral cDNAs to generate high-accuracy circular consensus sequences (CCSs; Travers et al., 2010). These CCSs retain the cell barcodes, and with sufficient sequencing depth we obtain CCSs from multiple unique UMI-tagged cDNAs for each viral gene in each cell. Because most cells are infected by just one or two virions, we can build a consensus of CCSs for each viral gene in each cell to determine the sequence(s) of these virions. Combining this information with the 3’-end sequencing determines the entire transcriptome and full sequences of the infecting virions in single cells (Figure 2E).

### Transcriptomic analyses of single IFN+ and IFN- influenza-infected cells

We obtained transcriptomes for 1,614 human (A549) cells, and 50 of the uninfected canine cells that were spiked into the experiment as a control (Figure 3A). We also obtained 12 transcriptomes with a mix of human and canine transcripts; from the number of such mixed cell-type transcriptomes, we estimate (Bloom, 2018) that ~11% of the transcriptomes are derived from multiple cells. To remove some of these multiplets along with low-quality droplets, we filtered transcriptomes with unusually high or low numbers of cellular transcripts as is commonly done in analysis of single-cell RNA-seq data (Haque et al., 2017). This filtering left 1,490 human cells for further analysis (Figure 3B).

**Figure 3.**
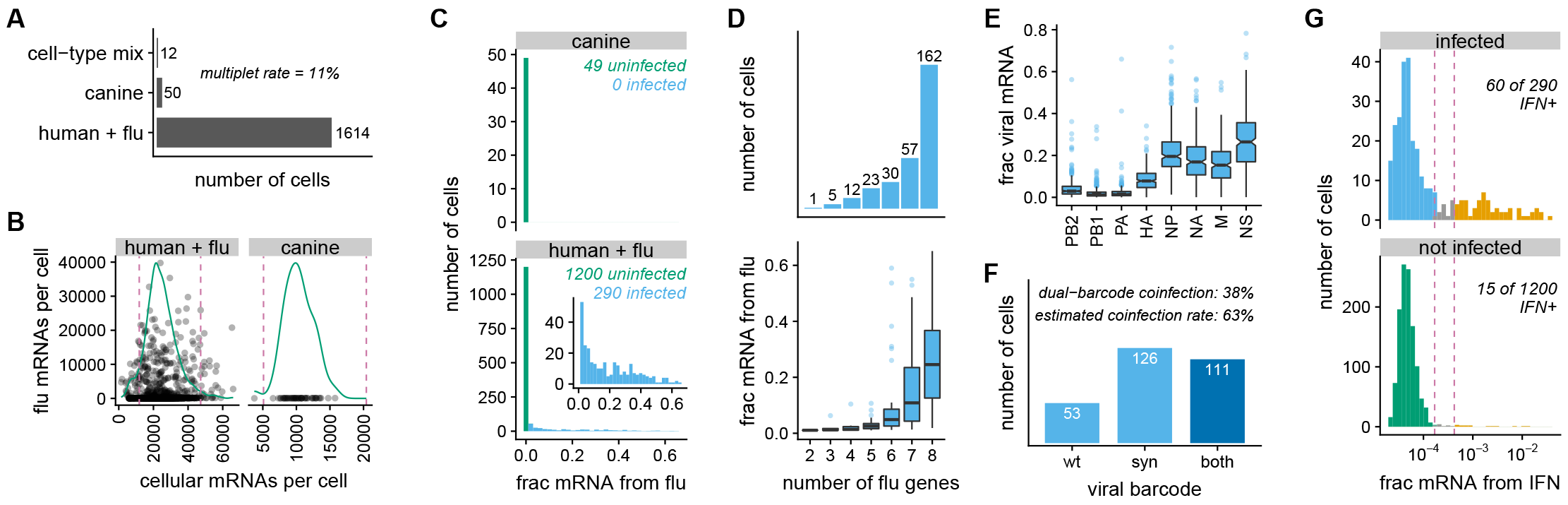
Single-cell transcriptomics of IFN-enriched influenza-infected cells. (**A**) Number of cells for which transcriptomes were obtained. (**B**) The number of cellular and viral mRNAs detected for each cell is plotted as a point. Green lines show the distribution of cellular mRNAs per cell. Cells outside the dashed magenta lines have unusually low or high amounts of cellular mRNA (likely low-quality emulsions or multiplets), and are excluded from subsequent analyses. (**C**) Distribution across cells of the fraction of all mRNA derived from influenza. Cells called as infected are in blue, while other cells are in green. The inset shows the amount of viral mRNA in the human cells that are called as infected. (**D**) Number of influenza genes detected per infected cell, and the amount of viral mRNA in cells expressing each number of viral genes. The majority of cells express all eight gene segments, but a substantial minority fail to express at least one gene. Figure S3A shows the frequency that each viral gene is detected. (**E**) Relative expression of viral genes, quantified as the fraction of all viral mRNA in each infected cell that is derived from each gene. (**F**) Number of cells infected with wild-type virus, synonymously barcoded virus, or both. From the cells infected with both viral barcodes, we estimate (Bloom, 2018) that 63% of infected cells are co-infected. (**G**) Fraction of cellular mRNA from IFN across cells, faceted by whether the cells are infected. Cells to the left of the first dashed magenta line are classified as IFN-, and cells to the right of the second line are classified as IFN+. A pseudocount is added to the number of IFN transcripts detected in each cell, which is why none of the fractions are zero. Many cells that do not express IFN still express ISGs (Figure S3C,D).

To identify infected cells, we examined the fraction of each transcriptome derived from virus (Figure 3C). As expected, only a small fraction (~0.7%) of transcripts in the uninfected canine cells were viral; this low-level background is from lysed cells that release ambient viral mRNA that is captured in droplets prior to reverse-transcription. We tested if each cell contained more viral transcripts than expected under a Poisson model given this background fraction, and classified 290 human cells as definitively infected (Figure 3C). We classified the other cells as uninfected, although it is possible that some were infected with virions that produced very little mRNA. The distribution of the amount of viral mRNA across infected cells is in the inset in Figure 3C. As in our prior work (Russell et al., 2018), the distribution is extremely heterogeneous: many infected cells have only a few percent of mRNA derived from virus, but viral mRNA comprises over half the transcriptome of a few cells.

We called the presence or absence of each viral gene in each infected cell, again using a Poisson model parameterized by background fractions estimated from uninfected canine cells. Figure 3D (top panel) shows that a slight majority (162 of 290) of infected cells express all eight genes (see Figure S3A for frequencies for individual genes). This measured frequency of infected cells expressing all eight genes is slightly higher than in our prior work (Russell et al., 2018) and studies by others (Brooke et al., 2013; Heldt et al., 2015; Dou et al., 2017), which estimated that only 13% to 50% of infected cells express all genes.

The amount of viral mRNA was lower in cells that failed to express viral genes (Figure 3D, bottom). However, viral burden remained variable even after conditioning on the number of viral genes: some cells that failed to express one or even two genes still derived >50% of their mRNA from virus, while other cells that expressed all genes had only a few percent of their mRNA from virus. Consistent with prior work (Russell et al., 2018), despite the wide variation in absolute expression of viral genes, their *relative* expression was fairly consistent (Figure 3E) and similar to values from older bulk studies (Hatada et al., 1989).

By examining the synonymous viral barcodes near the 3’ termini of transcripts, we determined that 38% of cells were co-infected with wild-type and synonymously barcoded virions (Figure 3F; cells called as co-infected if a binomial test rejected null hypothesis that ≥95% of viral mRNA is from one viral barcode variant). From Figure 3F, we estimate (Bloom, 2018) that 63% of infected cells are co-infected, implying that 25% are co-infected with two virions with the same viral barcode (such co-infections cannot be identified from transcriptomic data). This co-infection rate is higher than expected from the relative numbers of infected and uninfected cells (Figure 3C) if infection is Poisson. This discrepancy could arise if the MACS for IFN+ cells also enriches co-infected cells, if infection is not truly Poisson, or if co-infection increases the likelihood that we identify a cell as infected given the thresholds in Figure 3C. This moderately high rate of co-infection may also explain why more cells in our experiment express all eight viral genes compared to some prior studies, as a co-infecting virion can complement a missing viral gene.

We next examined expression of IFN and ISGs (Figure 3G and Figure S3B-D). Over 20% of infected cells were IFN+ given the heuristic thresholds in Figure 3G, indicating that the MACS enriched IFN+ cells far beyond their initial frequency of ~0.5% (Figure 1B). Few (~1.3%) uninfected cells were IFN+; the few that were present might be because the MACS enriched for rare cells that spontaneously activated IFN, or because some cells that we classified as uninfected were actually infected at low levels. Many more cells expressed ISGs than IFN itself: the ratio of ISG+ to IFN+ cells was 1.8 among infected cells, and 7 among uninfected cells (Figure S3C). The IFN+ cells were a subset of the ISG+ cells: IFN+ cells always expressed ISGs, but many ISG+ cells did not express IFN (Figure S3D). These results are consistent with the established knowledge that IFN is expressed only in cells that directly detect infection, but that ISGs are also expressed via paracrine signaling in other cells (Stetson and Medzhitov, 2006; Honda et al., 2006).

### Full genotypes of viruses infecting single IFN+ and IFN- cells

We next used PacBio sequencing (Figure 2D) to determine the full sequences of the viral genes expressed in single infected cells. Using PCR enrichment (see Methods), we obtained over 200,000 high-quality PacBio CCSs that mapped to an influenza gene and contained a cell barcode and UMI (Figure S4A). Crucially, the synonymous viral barcodes at both termini of each gene enabled us to confirm that PCR strand exchange was rare (Figure S4B), meaning that the vast majority of CCSs correctly link the sequence of the viral transcript to cell barcodes and UMIs that identify the cell and molecule of origin.

After calling the presence / absence of each viral gene in each cell using the transcriptomic data as described in the previous section, we called mutations if they were found in at least two CCSs originating from different mRNAs (unique UMIs) and at least 30% of all CCSs for that gene in that cell. For cells co-infected with both viral barcode variants, we called mutations separately for each variant. This strategy identifies mutations in virions that initiate infection of cells infected with at most one virion of each viral barcode variant (~75% of infected cells), as well as high-abundance mutations in cells co-infected with multiple virions of the same viral barcode. It will not identify mutations that arise within a cell after the first few rounds of viral genome replication, since such mutations will not reach 30% frequency in that cell. Therefore, analogous to somatic variant calling in tumor sequencing (Xu et al., 2014; Cibulskis et al., 2013), there is a limit to our detection threshold: we cannot identify mutations that occur on just a small fraction of transcripts in a cell.

We called the sequences of all expressed viral genes in the majority of infected cells (Figure S5). We were most effective at calling viral genotypes in cells that expressed high amounts of viral mRNA and were infected by only one viral barcode variant (Figure S5). But we also called genotypes for many cells that had low viral burden or were co-infected by both viral barcode variants.

The 150 cells for which we called the viral genotypes are shown in Figure 4. Inspection of this figure reveals a wealth of anecdotal relationships between viral genotype and infection outcome. For instance, the cell with the highest viral burden (*cell 1* in Figure 4, which has 65% of its mRNA from virus) was infected by a virion that expressed unmutated copies all genes and did not induce detectable IFN. But 12 of the other 13 cells with at least 50% of their mRNA from virus were infected by virions that had a mutation or failed to express a gene, and five of these cells expressed IFN. As expected, all cells infected by virions that failed to express a component of the viral polymerase complex (PB2, PB1, PA, or NP) expressed low amounts of viral mRNA since they are limited to primary transcription off the incoming proteins (e.g., *cell 132* and *cell 143*).

**Figure 4.**
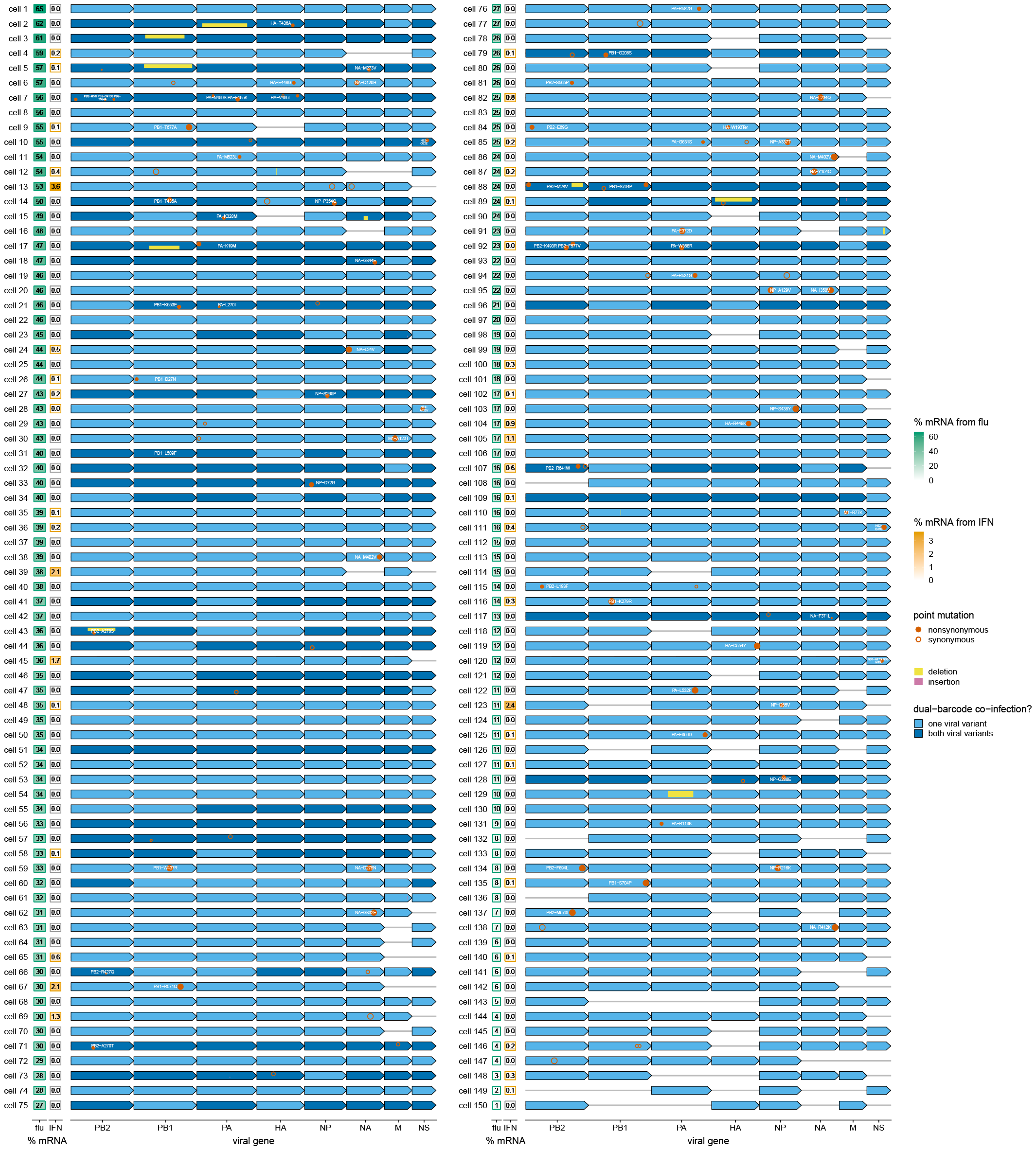
Viral genotypes and infection outcomes in single cells. Green and orange boxes at the left show the percent of all mRNA in that cell derived from virus and the percent of all cellular mRNA derived from IFN, respectively. The second box is framed in orange for cells classified as IFN+ in Figure 3G. Blue arrows indicate the presence of a viral gene from one (light blue) or both (dark blue) viral barcode variants; a dark blue arrow therefore means that a cell was co-infected. Circles and boxes on the arrows indicate mutations or indels as described in the legend at right. The circle areas and box heights are proportional to the fraction of CCSs with that mutation. For dual-barcode infections, mutations / indels for the wild-type and synonymously barcoded viral variants are shown in the top and bottom half of the arrows, respectively. For instance, *cell 5* was co-infected such that it expressed both unmutated and internally deleted PB1.

The two cells that expressed the most IFN (*cell 13* and *cell 123*) both lacked the viral NS gene that encodes the virus’s primary IFN antagonist, NS1 (García-Sastre et al., 1998; Hale et al., 2008). Many other IFN+ cells also lacked NS or had different defects such as large internal deletions (e.g., *cell 5* and *cell 89*) or amino-acid mutations (e.g., *cell 9, cell 28*, and many others).

However, Figure 4 also reveals stochasticity that is independent of viral genotype. This stochasticity sometimes acts to the detriment of the virus, and sometimes to the detriment of the cell. As an example of the former case, expressing unmutated copies of all eight genes does not guarantee a favorable outcome for the virus: for instance, the unmutated virion that infected *cell 139* only expressed viral mRNA to 6% of the total transcriptome, and the unmutated virion that infected *cell 105* still induced IFN. But in other cases, the stochasticity allows a clearly defective virus to still escape immune recognition. For instance, there are a number of cells (e.g., *cell 62* and *cell 78*) that do not activate IFN despite being infected by virions that fail to express NS.

### Viral defects associated with infection outcome in single cells

To systematically assess viral features associated with infection outcome, we divided the 150 cells with viral genotypes into those that expressed unmutated copies of all eight genes (disregarding synonymous mutations) and those that did not. Figure 5A shows that the 49 cells infected by unmutated virions had a tighter distribution of the viral mRNA per cell than the other 101 cells as quantified by the Gini index (Gini, 1921). Therefore, viral defects are a major contributor to the heterogeneity in viral transcriptional burden among cells.

**Figure 5.**
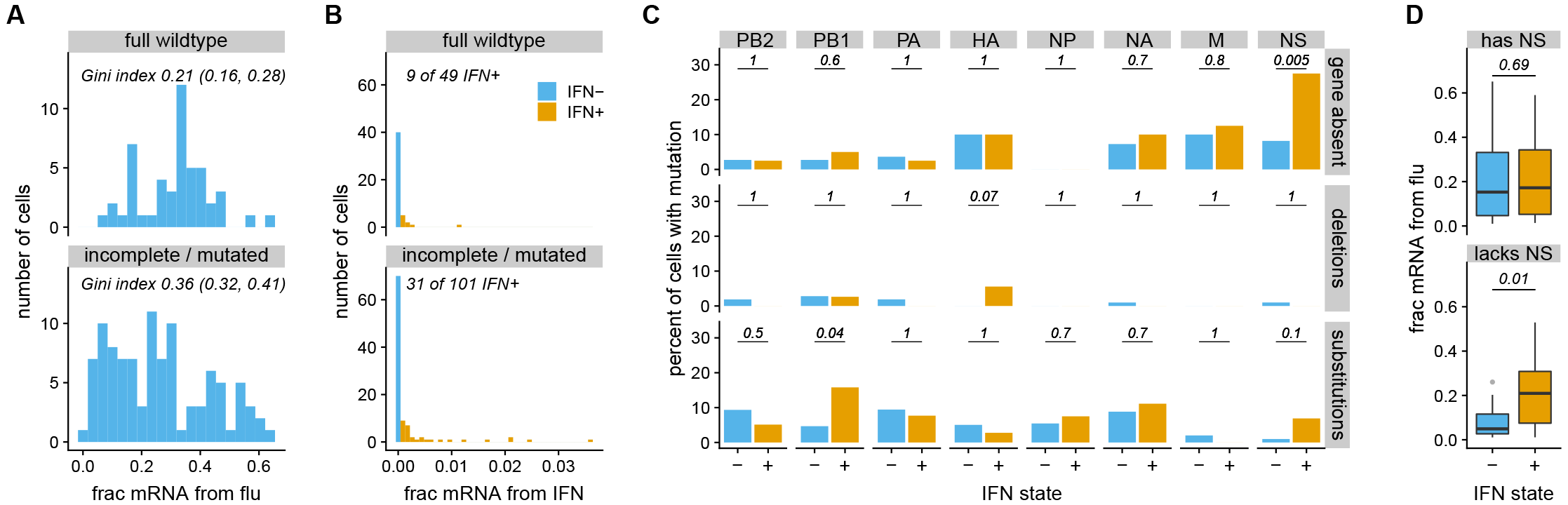
Viral features associated with heterogeneity among cells for which we determined viral genotypes. (**A**) Percent of all mRNA derived from virus, faceted by whether cells express unmutated copies of all eight genes. Cells infected by unmutated virions exhibit less heterogeneity in viral burden as quantified by the Gini index (95% confidence intervals indicated). (**B**) IFN expression among cells expressing unmutated copies of all genes, and cells with mutations or missing genes. (**C**) Specific viral defects associated with IFN induction. The top panel show the percent of IFN- and IFN+ cells that fail to express each viral gene. The middle and bottom panels show the percent of IFN- and IFN+ cells with a deletion or amino-acid substitution in each gene, conditioned on the gene being expressed. Numbers give P-values (Fisher’s exact test) for rejecting null hypothesis that percents are equal among IFN- and IFN+ cells. (**D**) There is no association between IFN induction and the amount of viral mRNA in cells that express NS, but viral burden is associated with IFN among cells that lack NS. This figure only shows non-synonymous substitutions, and disregards insertions as they are very rare.

Cells infected by incomplete or mutated virions also expressed IFN more frequently than cells infected by full wild-type virions (Figure 5B), although this difference was not statistically significant (*P* = 0.12, Fisher’s exact test). However, some specific viral defects were significantly associated with IFN induction: absence of NS and amino-acid mutations in PB1 were significantly enriched in IFN+ cells, and amino-acid mutations in NS and deletions in HA were weakly enriched (Figure 5C). Due to the low number of cells and large number of hypotheses in Figure 5C, the only trend that remained significant at a false discovery rate of 10% was absence of NS. However, the validation experiments in the next section show that the lack of statistical significance for many of the trends is due to the modest number of infected cells that were sequenced rather than a lack of true association between viral defects and IFN induction.

One other interesting trend emerges from the single-cell data. There is no difference in the amount of viral mRNA between IFN+ and IFN- cells that express NS (Figure 5D). But among cells that lack NS, the IFN+ ones have significantly more viral mRNA (Figure 5D). This finding is elaborated on in the validation experiments below.

### Validation that viral defects in single IFN+ cells often increase IFN induction

To test if the viral defects identified in single IFN+ cells directly increase IFN expression, we used reverse genetics to generate bulk stocks of viruses with some of these defects.

The viral defect most strongly associated with IFN induction was failure to express the NS gene (Figure 4, Figure 5C). Although it is sometimes possible to use complementing cells to generate influenza viruses lacking a specific gene (Fujii et al., 2003; Marsh et al., 2007), we were unable to generate viruses that lacked NS. We therefore mimicked the effect of absence of NS by creating a mutant virus (NS1stop) that had multiple stop codons early in the NS1 coding sequence.

The single-cell data also showed that amino-acid substitutions in PB1 and NS were enriched in IFN+ cells (Figure 4, Figure 5C), so we created mutant viruses with some of the substitutions found in IFN+ cells: PB1-D27N, PB1-G206S, PB1-K279R, PB1-T677A, NS1-A122V, and NS2-E47G.

Finally, prior work has suggested that virions with internal deletions in the polymerase genes can induce IFN (Baum et al., 2010; Tapia et al., 2013; Boergeling et al., 2015; Dimmock and Easton, 2015). Although such deletions are not significantly enriched among IFN+ cells in our single-cell data (Figure 5C), there is a co-infected IFN+ cell where one viral variant has a deletion in PB1 spanning nucleotides 385 to 2163 (*cell 5* in Figure 4). We created a virus carrying this deletion, and propagated it in cells constitutively expressing PB1 protein.

We tested the frequency of IFN induction by each viral stock using the reporter cells. Figure 6 shows that five of eight mutant viral stocks induced IFN more often than a wild-type viral stock. The strongest IFN induction was by the NS1stop virus, but the PB1 internal deletion and three point-mutant viruses (PB1-D27N, PB1-T677A, and NS1-A122V) also induced IFN more than wild type. The other three point mutants (PB1-G206S, PB1-K279R, and NS2-E47G) did not increase IFN induction—an unsurprising finding, since we expect some mutations in IFN+ cells by chance.

**Figure 6.**
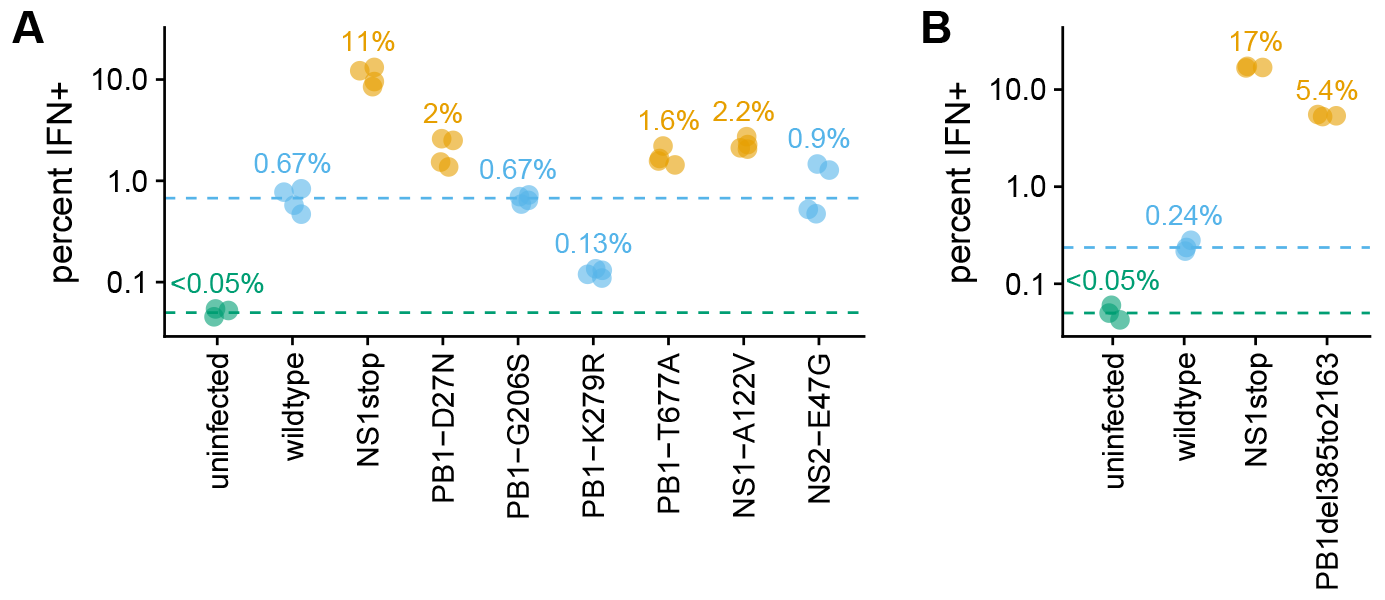
Validation that IFN induction is increased by mutations identified in the single-cell virus sequencing of IFN+ cells. (**A**) Percent of infected cells that become IFN+ after infection with a bulk stock of the indicated mutant, as determined using a reporter cell line. Numbers give the median of four measurements for each mutant. The limit of detection of 0.05% is indicated with a dashed green line, and the median value for wild type is indicated with a dashed blue line. Points are orange if the mutant virus stock induces IFN more frequently than the wild-type viral stock (one-sided t-test, *P* < 0.01), and blue otherwise. (**B**) Similar to the first panel, but validates the increased IFN induction for a large internal deletion in the PB1 gene. See Figure S6 for more detailed experimental data. The experiments in the two panels were performed on different days, and so numerical values can be compared within but not between panels.

However, IFN induction remains stochastic even for the most potently IFN-inducing viral mutants. Figure 6 shows flow cytometry data, which is itself a single-cell measurement, albeit one that does not report the viral genotype. These data reveal that no mutant virus stock induces IFN in more than a fraction of cells. Of course, the mutant virus stocks are themselves genetically heterogeneous, as many virions will have additional defects similar to those revealed by our single-cell sequencing of the “wild-type” viral stock. But our single-cell data show that IFN induction is stochastic even for infections with the same defect, such as absence of NS (e.g., compare *cell 62* and *cell 69* in Figure 4). Therefore, the experiments in Figure 6 not only validate some viral defects that increase IFN induction, but also that induction is stochastic even with these defects.

The single-cell data also suggest heterogeneity in the process by which different viral variants induce IFN. Specifically, these data show that NS-deficient virions are much more likely to induce IFN when they transcribe more RNA, but that there is no such association for other virions (Figure 5D). To validate this conclusion, we used flow cytometry to compare IFN induction among cells that express low or high levels of the viral HA protein under the assumption that protein levels correlate with viral transcription. Consistent with the single-cell data, the NS1stop or NS1-A122V viruses are much more likely to induce IFN if they express high levels of HA (Figure 7A and Figure S7A). In contrast, wild-type or PB1-mutant viruses are only slightly more likely to induce IFN if they express more HA. To confirm that higher viral transcription or genome replication drive IFN induction by NS-deficient virions, we treated infected cells with ribavirin or cycloheximide, which inhibit secondary viral transcription (Vanderlinden et al., 2016; Reuther et al., 2015; Scholtissek, 1976; Killip et al., 2014). Both inhibitors strongly reduced IFN induction by cells infected with NS1stop virus, but not cells infected by virus with a deletion in PB1 (Figure 7B and Figure S7B). These data indicate that different viral variants trigger production of IFN by different processes.

**Figure 7.**
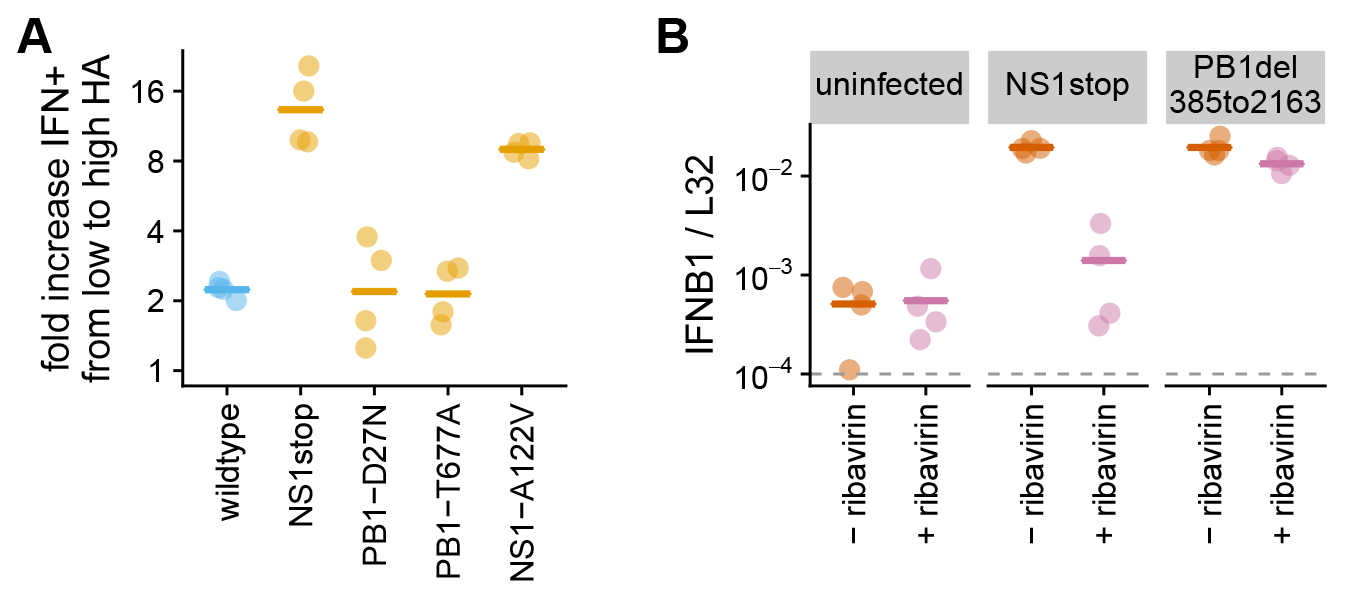
Higher viral gene expression is strongly associated with more IFN induction only in NS1-deficient infections. (**A**) For viruses with defects in NS1, there is more IFN induction among infected cells that express higher levels of HA protein. The y-axis shows the ratio of the percent of IFN+ cells in the highest HA-expression quartile relative to the lowest HA-expression quartile as determined by flow cytometry. Points indicate replicates, and lines indicate the mean. See Figure S7A for details. (**B**) Blocking viral secondary transcription strongly decreases IFN in cells infected with NS1stop virus, but not cells infected with virus carrying a deletion in PB1. A549 cells were infected at doses that induced equivalent IFN, and ribavirin was added at a concentration (100 μM) sufficient to block secondary transcription (Vanderlinden et al., 2016; Reuther et al., 2015; Scholtissek, 1976). After eight hours, IFNB1 transcript was quantified relative to the housekeeping gene L32 by qPCR. The dashed gray line indicates the limit of detection. Figure S7B shows that results are similar if secondary transcription is instead blocked with cycloheximide.

## DISCUSSION

We have determined the full sequences of all viral genes expressed in single influenza-infected cells, and examined how viral mutations and gene-expression defects associate with infection outcome in each cell. Methodologically, our major advance is to measure the *genotypes* of viruses in addition to the abundance of viral components (i.e., transcripts, proteins, or progeny virions) as has been done by prior single-cell studies (Russell et al., 2018; Zanini et al., 2018a,b; Steuerman et al., 2018; Saikia et al., 2019; O’Neal et al., 2018; Zhu et al., 2009; Schulte and Andino, 2014; Akpinar et al., 2016; Heldt et al., 2015; Brooke et al., 2013). Our method builds on the observation by that fragmentary viral genetic information can be obtained by Illumina-based single-cell transcriptomic techniques (Saikia et al., 2019; Zanini et al., 2018b). To make this information complete, we have coupled single-cell transcriptomics with long-read PacBio sequencing of viral genes, a strategy analogous to that used by Gupta et al. (2018) to obtain full-length isoforms of some cellular genes in single cells.

This viral genetic information is crucial for understanding infection outcome and innate-immune induction. Despite the fact that we used a low-passage viral stock generated from plasmids encoding a “wild-type” influenza genome, most infected cells do not express wild-type copies of all viral genes. Although our study is certainly not the first to note that influenza has a high mutation rate (Parvin et al., 1986; Suarez et al., 1992; Bloom, 2014; Pauly et al., 2017) and sometimes fails to express genes (Brooke et al., 2013; Heldt et al., 2015; Dou et al., 2017; Russell et al., 2018), it is the first to directly observe the full spectrum of defects across single cells. Inspection of Figure 4 shows how any experiment that does not sequence viral genes in single cells is averaging across a vast number of hidden viral defects.

These viral defects substantially contribute to the heterogeneity observed in prior singlecell studies of influenza (Russell et al., 2018; Steuerman et al., 2018; Heldt et al., 2015; Sjaastad et al., 2018), including the probability that a cell triggers an IFN response. We identified four types of defects in single IFN+ cells that we validated increase IFN induction. Two types of defects—absence of NS and mutations to the NS1 protein—presumably impair NS1’s well-known ability to antagonize innate-immune pathways (García-Sastre et al., 1998; Hale et al., 2008). The third type of defect, amino-acid mutations in the PB1 protein, is consistent with recent work by showing that mutations to the viral polymerase affect generation of mini-viral RNAs that activate the innate-immune sensor RIG-I (te Velthuis et al., 2018). Finally, we found an internal deletion in PB1 that enhances IFN induction, consistent with prior work showing that such deletions are immunostimulatory (Baum et al., 2010; Tapia et al., 2013; Boergeling et al., 2015; Dimmock and Easton, 2015). Given the extensive prior work describing such deletions, it is surprising we did not identify more of them in our single IFN+ cells. There may be several reasons: we used relatively pure viral stocks (Xue et al., 2016) at modest MOI; our experiments preferentially captured cells with higher viral transcriptional load; and most prior studies have used techniques that can detect large deletions but not subtle point mutations. The relative importance of different defects also likely varies across conditions, viral strains, and cell types—so it remains an open question as to which defects are most relevant for early immune detection of actual viral infections in humans.

But although variation in viral populations is clearly important, it only partially explains heterogeneity among influenza-infected cells. We find substantial breadth in viral transcriptional burden and occasional IFN induction even among cells infected with unmutated virions. Additionally, no viral defect induces IFN deterministically—and the most immunostimulatory defect (absence of NS) also occurs in multiple IFN- cells in our single-cell dataset. Of course, our single-cell virus sequencing only calls mutations present in a cell at high frequency. However, we suspect that stochasticity or pre-existing cellular states explain most remaining heterogeneity, since IFN induction is heterogeneous even among cells treated with synthetic innate-immune ligands (Shalek et al., 2013, 2014; Wimmers et al., 2018; Bhushal et al., 2017) or other viruses (O’Neal et al., 2018).

Perhaps the most intriguing question is how the heterogeneity that we have described ultimately affects the macroscopic outcome of infection. Natural human influenza infections are initiated by just a handful of virions (McCrone et al., 2018; Xue and Bloom, 2018; Varble et al., 2014) that then undergo exponential growth, and early IFN responses are amplified by paracrine signaling (Stetson and Medzhitov, 2006; Honda et al., 2006). It is therefore plausible that early heterogeneity could have a large effect on downstream events. Extending our approaches to more complex systems could shed further light on how viral variation interacts with cell-to-cell heterogeneity to shape the race between virus and immune system.

## ACKNOWLEDGMENTS

We thank Cole Trapnell, Jason Underwood, Robert Bradley, Daniel Stetson, AJ Velthuis, Katherine Xue, and Hannah Itell for helpful suggestions. We thank Andy Marty and the Fred Hutch Genomics Core for performing the deep sequencing. This work was supported by the NIAID of the NIH under grant R01 AI127893 and a Burroughs Wellcome Fund Young Investigator in the Pathogenesis of Infectious Diseases grant to JDB. ABR was supported by a postdoctoral fellowship from the Damon Runyon Cancer Research Foundation (DRG-2227-15). JRK was supported by a Washington Research Foundation Undergraduate Research Fellowship and a Mary Gates undergraduate research scholarship from the University of Washington. JDB is an Investigator of the Howard Hughes Medical Institute.

## AUTHOR CONTRIBUTIONS

ABR and JDB designed the study. ABR performed the experiments with assistance from JRK. ABR and JDB computationally analyzed the data and wrote the paper.

## DECLARATION OF INTERESTS

The authors declare no competing interests.

## METHODS

### IFN reporter cell lines

We created IFN reporter variants of the A549 human lung epithelial cell line (Figure 1A). The parental A549 cell line used to create these reporters was obtained from ATCC (CCL-185), and was tested as negative for mycoplasma contamination by the Fred Hutch Genomics Core and authenticated using the ATCC STR profiling service. The cells were maintained in D10 media (DMEM supplemented with 10% heat-inactivated fetal bovine serum, 2 mM L-glutamine, 100 U of penicillin / ml, and 100 *μ*g of streptomycin / ml) at 37°C and 5% carbon dioxide.

To create the type I interferon reporters, a 1kb promoter region upstream of the human IFNB1 gene were cloned into the pHAGE2 lentiviral vector (O’Connell et al., 2010), with a NotI site immediately downstream of the promoter serving as an artificial Kozak sequence. Downstream of this NotI site, each of the following reporter constructs was cloned: mCherry, mNeonGreen, and low-affinity nerve growth factor lacking the C-terminal signaling domain (LNGFRΔC) (Bonini et al., 1997; Ruggieri et al., 1997) linked to mNeonGreen by a P2A linker (Kim et al., 2011). The sequence of the last of these constructs is provided in File S1.

To create the type III interferon reporters, a 1.2kb region upstream of the human IL29 (IFNL1) gene was cloned into the pHAGE2 vector, with the native Kozak sequence retained at the 3’ end. Downstream of this promoter we cloned LNGFRΔC linked to ZsGreen via a P2A linker. The sequence of this construct is provided in File S1.

We used these constructs to generate lentiviral vectors and transduce of A549 cells in the presence of 5 *μ*g polybrene. We then sorted single transduced cells and expanded them. A portion of the expanded cells were tested for reporter activity by transfecting poly(I:C) (a potent agonist of the RIG-I pathway), and we retained clones with strong activation. Importantly, the cells that we retained for further use were not the same portion that were tested by poly(I:C) treatment, but rather a separate split of the same population—this avoids any selection on the cells from transient activation of IFN. For the dual type I / type III reporter used in Figure S1B, a single-cell clone of the type III reporter cell line was transduced with the type I reporter bearing the mCherry fluorescent marker, and then isolated and propagated as a single cell clone for the other cell lines. All reporter lines tested negative for mycoplasma contamination by the Fred Hutch Genomics Core.

Figure S1A shows validation of the reporter cell lines using infection with saturating amounts of the Cantell strain of Sendai virus (obtained from Charles River Laboratories). For detection of the cell-surface bound LNGFRΔC, cells were stained with PE-conjugated anti-LNGFR (CD271) antibody from Miltenyi Biotec.

### Viruses for single-cell experiments

We performed the single-cell experiments using the A/WSN/1933 (H1N1) strain of influenza virus. We used both the wild-type virus and a variant of the virus where synonymous mutations were added within a few 100 nucleotides of each termini of each gene segment. We have used a similar synonymous viral barcoding strategy in our prior single-cell work (Russell et al., 2018) as it allows us to detect about half of co-infected cells based on the expression of both viral barcode variants. In the current work, we extended this approach by placing synonymous barcodes near *both* termini of the gene segments in order to quantify strand exchange during PacBio sequencing (Figure S4B). The sequences of all gene segments from the wild-type and synonymously barcoded viral strains are in File S1. These genes were cloned into the pHW2000 (Hoffmann et al., 2000) reverse-genetics plasmid.

Both viral strains were generated by reverse genetics using the pHW18* series of bi-directional plasmids (Hoffmann et al., 2000). We controlled the durations and MOI during viral passaging since these factors can greatly affect the accumulation of defective viral particles (Xue et al., 2016). The viruses were generated by reverse genetics in co-cultures of 293T and MDCK-SIAT1 cells in influenza growth media (Opti-MEM supplemented with 0.01% heat-inactivated FBS, 0.3% BSA, 100 U of penicillin/ml, 100 *μ*g of streptomycin/ml, and 100 *μ*g of calcium chloride/ml) and then propagated in MDCK-SIAT1 cells in influenza growth media using the same basic procedures detailed in (Russell et al., 2018). Specifically, after generation by reverse genetics, the wild-type variant was expanded at an MOI of 0.001 for 72 hours twice in MDCK-SIAT1 cells, and the synonymously barcoded variant was expanded once at an MOI of 0.01 for 60 hours. The MOIs for this passaging are based on titers determined using TCID50 assays via the formula of Reed and Muench (Reed and Muench, 1938) as implemented at https://github.com/jbloomlab/reedmuenchcalculator.

### Flow cytometry analyses for HA expression

For the single-cell experiments (which only examine the transcriptional results of a single cycle of infection), we were most interested in the titer of viral particles that are transcriptionally active for a single round of infection of A549 cells. We estimated titers of transcriptionally active virions by staining for HA expression in virus-infected A549 cells. Specifically, we infected A549 cells (or one of the A549 reporter cell line variants as indicated) in influenza growth medium, and at 13 to 14 hours post-infection, we trypsinized cells, re-suspended in phosphate-buffered saline (PBS) supplemented with 2% heat-inactivated fetal bovine serum (FBS), and stained with 10 *μ*g/ml of H17-L19, a mouse monoclonal antibody previously shown to bind to the HA from the A/WSN/1933 strain of virus (Doud et al., 2017). After washing in PBS supplemented with 2% FBS, the cells were stained with a goat anti-mouse IgG antibody conjugated to APC, washed, fixed in 1% formaldehyde in PBS, washed again, and then analyzed by flow cytometry to determine the fraction expressing detectable HA protein.

### Single-cell transcriptomics of IFN-enriched infected cells using 10X Chromium

The single-cell transcriptomics and virus sequencing was performed using the A549 cells with the *IFNB1* LNGFRΔC-P2A-mNeonGreen reporter. A schematic of the experiment is shown in Figure 2.

The wild-type and synonymously barcoded viruses were mixed with the goal of adding equal numbers of transcriptionally active HA-expressing virions of each virus strain. The cells were then infected with this mixture at a dose designed to infect about half the cells (Figure 3C suggests that the actual rate of detectable infection was slightly lower). Infections were allowed to proceed for 12 hours. The cells were then trypsinized, the trypsin was quenched with D10 media, and cells were resuspended in de-gassed PBS supplemented with 0.5% bovine serum albumin and 5 mM EDTA. To enrich IFN+ cells, the cells were then incubated with anti-LNGFR MACSelect Microbeads (Miltenyi Biotec) and twice passed over an MS magnetic column (Miltenyi Biotec), retaining the bound (and presumably IFN-enriched) population each time. This MACS sorting is expected to give approximately the enrichment for IFN+ cells shown in Figure S2. The original, unsorted, population was then added back in to ~10% of the final cell fraction in order to ensure the presence of interferon negative cells. At this point, uninfected canine (MDCK-SIAT1) cells were also added to ~5% of the final cell fraction to enable quantification of the cell multiplet rate (Figure 3A) and background viral mRNA in uninfected cells (Figure 3C). We began this entire process of cell collection and enrichment at 12 hours post-infection, but the process (which was performed at room temperature) took about an hour, and thus we consider the cells to have been analyzed at 13 hours post-infection. The final cell suspension was counted using a disposable hemocytometer and loaded on the 10x Genomics Chromium instrument (Zheng et al., 2017), targeting capture of ~1,500 cells.

This sample was then processed to create libraries for Illumina 3’-end sequencing according to the 10X Genomics protocol using the Chromium Single *Cell 3*’ Library and Gel Bead Kit v2 with one important modification: rather than process all full-length cDNA through enzymatic fragmentation, several nanograms were retained for targeted full-length viral cDNA sequencing as described below. The single-cell transcriptomics library was sequenced on an Illumina HiSeq 2500, and the data analyzed as described below.

### Enrichment and preparation of viral cDNA for PacBio sequencing

We amplified virus-derived molecules from the small amount of cDNA retained from the 10X Genomics protocol for PacBio sequencing of the full-length cDNA. All molecules in this cDNA have at their 3’ end the cell barcode and UMI plus the constant adaptor sequence that is added during the 10X protocol (see Figure 2 for simple schematic, or the detailed analysis notebook at https://github.com/jbloomlab/IFNsorted_flu_single_cell/blob/master/pacbio_analysis.ipynb or File S5 for at detailed schematic). However, we only wanted to PacBio sequence cDNA molecules derived from virus, since it would be prohibitively expensive to sequence all molecules. We therefore needed to enrich for the viral molecules, a process made challenging by the need to also retain the 10X adaptor / UMI / cell barcode at the 3’ end (the primer at the 5’ end can be specific, but the one at the 3’ end must bind the common 10X adaptor shared among all molecules).

We first performed a multiplex PCR reaction on 1 ng of the full-length 10X cDNA using a 3’ primer complementary to the common 10X adaptor, and a multiplex mix of eight 5’ primers, one specific for the mRNAs from each of the eight viral gene segments (although some viral segments produce multiple splice forms, all mRNAs from a given segment share the same 5’ end). The sequences of these primers are in File S2. A major concern during these PCRs is strand exchange (see Figure S4B) which would scramble the cell barcodes and mutations on viral cDNAs. To reduce strand-exchange and hopefully obtain more even PCR amplification across segments, we performed emulsion PCRs using the Micellula DNA Emulsion Kit (Roboklon). Emulsion PCR involves encapsidating DNA template molecules in a reverse-phase emulsion, with each template positive droplet serving as a microreactor. This process physically separates disparate template molecules, preventing strand exchange and allowing each molecule to be amplified to exhaustion of its droplet’s reagents without competing with the broader pool of PCR-amenable molecules (Boers et al., 2015). We performed the PCRs using Kapa HiFi Hotstart ReadyMix, supplementing the reactions with additional BSA to a final concentration of 0.1 mg/ml and using a volume 100*μ*l. Both the common 3’ primer and the multiplex mix of eight 5’ primers were added to a final concentration of 0.5*μ*M. We performed 30 cycles of PCR, using an extension time of 2 minutes 15 seconds at 67°C, and a melting temperature of 95°C. This melting temperature is lower than the standard 98° C melting step suggested by the manufacturer for Kapa HiFi because we wanted to avoid collapse of emulsion integrity at high temperature. We performed 30 cycles in order to saturate the reagents in each emulsion—unlike for open PCR reactions, the physical occlusion of amplified material in emulsion PCR reduces artifacts at high cycle numbers.

Because were were concerned that the multiplex PCR would still result in highly uneven amplification of different influenza cDNAs due to differences in their expression levels and lengths, the product of this multiplex PCR was subjected to eight additional individual emulsion PCR reactions, each using only a single segment-specific 5’ primer as well as the common 3’ primer, using 1 ng of material in each reaction. The material from these eight segment-specific PCRs was then pooled with the goal of obtaining a equimolar ratio of segments, and sequenced on one SMRT Cell in a PacBio RS II and one SMRT Cell of a PacBio Sequel. Detailed results from the analysis of these first two sequencing runs is shown in the PacBio analysis notebook at https://github.com/jbloomlab/IFNsorted_flu_single_cell/blob/master/pacbio_analysis.ipynb and File S5. These results showed that although the PCRs substantially enriched for influenza molecules, the relative coverage of the different viral genes was still very uneven, with the longer genes (especially the polymerase genes) severely under-sampled.

To try to improve coverage of the polymerase genes, we produced two new sequencing pools: one consisting of the five shortest viral segments (HA, NP, NA, M, and NS) from the aforementioned segment-specific emulsion PCRs, and the other consisting of the three longer polymerase segments (PB2, PB1, and PA). The former was sequenced on one cell of a single SMRT Cell of a PacBio Sequel, and the latter on two additional SMRT Cells of a PacBio Sequel. As is shown in the PacBio analysis notebook at https://github.com/jbloomlab/IFNsorted_flu_single_cell/blob/master/pacbio_analysis.ipynb (see also File S5), even with these new sequencing runs the coverage remained relatively low for the polymerase genes—and most of the reads we did obtain were dominated by shorter internally deleted variants of the polymerase genes, which arise commonly during influenza replication (Xue et al., 2016) and are presumably preferentially amplified during PCR.

To obtain more reads for longer full-length polymerase variants, we therefore subjected 10 ng of our amplified material for each polymerase segment to a bead selection using SPRIselect beads at a volume ratio of 0.4 to select for larger molecules. This selection removes most low-molecular weight DNA species including internally-deleted defective segments. Material from this selection was then amplified using 16 (PB1) or 14 (PB2 and PA) cycles of a non-emulsion PCR using the standard conditions recommended by the Kapa HiFi Hotstart ReadyMix (extension at 67°C for 2 minutes 15 seconds, and melting at 98°C). The use of relatively few PCR cycles was designed to prevent the occurrence of the artifacts (including strand exchange) that occur in non-emulsion PCRs as the reactions approach saturation. We pooled the products of these reactions from this size-selection and sequenced on a SMRT Cell of a PacBio Sequel. As is shown in the PacBio analysis notebook at https://github.com/jbloomlab/IFNsorted_flu_single_cell/blob/master/pacbio_analysis.ipynb (see also File S5), this sequencing yielded modestly more full-length polymerase variants, but they were still heavily undersampled compared to other viral genes.

To further to improve recovery of full-length PB1, PB2, and PA, we therefore took an alternative approach that allowed us to perform a specific PCR for full-length polymerase variants. Specifically, we circularized the template molecules (including the full length of genes plus the cell barcode, UMI, and 10X adaptor), and then used two segment-specific primers that annealed in apposition near the center of each polymerase gene to linearize these circular molecules. Only molecules that contain the middle of the polymerase genes (which are typically full-length) are linearized by this process. In the downstream computational analysis, we can then determine the full sequence of the gene as well as the cell barcode of the initial molecule from which the linearized molecule is derived. Specifically, we first used 2.5 ng of our already-amplified segment-specific material in a 10-cycle PCR to append circularization adapters (see File S2 for sequences), and cleaned the resultant mixture using SPRIselect beads at a volume ratio of 0.4. We then used 10 ng of this amplified material in a 20*μ*l NEBuilder reaction using an extended reaction time of 50 minutes in order to circularize the molecules. We next incubated these reactions for 1 hour at 37^°^ C with exonuclease V and additional ATP to a final increase in concentration of 1 mM to digest all non-circularized molecules. The circularized and digested material was then cleaned using SPRIselect beads at a volume ratio of 0. 4. This material was then used as template for three non-emulsion PCRs specific to PB2, PB1, or PA, using two segment-specific primers that align to the central portion of each gene but in apposition to each another (see File S2 for sequences). These linearization reactions used 20 (PB2) or 26 (PB1 and PA) PCR cycles, and the resulting products were cleaned using SPRIselect beads at a volume ratio of 1.0. This material was pooled to produce an equimolar mixture of full-length PB1, PA, and PB2 and sequenced in an additional SMRT Cell of PacBio Sequel. As is shown in the PacBio analysis notebook at https://github.com/jbloomlab/IFNsorted_flu_single_cell/blob/master/pacbio_analysis.ipynb (see also File S5), this process efficiently yielded many full-length polymerase variants.

The computational analyses of the full-length viral gene sequences described below used the combination of the data from all of these reactions. The number of sequences obtained for each gene after pooling the data from all reactions is shown in Figure S4A, which also indicates that the net rate of strand exchange is very low (see Figure S4B for an illustration of how this is determined). A more detailed breakdown of the coverage of each gene and data showing a consistently low rate of strand exchange for all PacBio runs is at https://github.com/jbloomlab/IFNsorted_flu_single_cell/blob/master/pacbio_analysis.ipynb (see also File S5). Importantly, the various PCR biases and enrichment schemes mean that the coverage of molecules by the PacBio sequencing is not proportional to their original abundance in the starting mRNA. However, as described in the computational analysis section below, the final analyses use the cell barcodes and UMIs in conjunction with the standard 10X Illumina sequencing to ensure that none of the conclusions are affected by the disproportionate amplification of some molecules during the PacBio library preparation (for instance, duplicate UMIs are removed from the PacBio data, and all conclusions about gene abundance or absence are based on the Illumina data).

### qPCR for viral genes and IFN

We performed qPCR on reverse-transcribed mRNA for influenza HA (to quantify viral transcription), IFNB1 (to quantify IFN induction), and L32 (a cellular housekeeping gene for normalization). For the qPCR, we used the SYBR Green PCR Master Mix (Thermo Fisher) according to the manufacturer’s protocol using oligo-dT primers. The qPCR primers were: HA primer 1, 5’-GGCCCAACCACACATTCAAC-3’; HA primer 2, 5’-GCTCATCACTGCTAGACGGG-3’; IFNB1 primer 1, 5’-AAACTCATGAGCAGTCTGCA-3’; IFNB1 primer 2, 5’-AGGAGATCTTCAGTTTCGGAGG-3’; L32 primer 1, 5’-AGCTCCCAAAAATAGACGCAC-3’; L32 primer 2, 5’-TTCATAGCAGTAGGCACAAAGG-3’.

For the qPCR in Figure S6C, A549 cells were seeded at a density of 10^4^ cells/well in a 96-well plate in D10 media 24 hours prior to infection, with four independent wells seeded per experimental treatment. Immediately prior to infection D10 media was removed and replaced with influenza growth media and infected with the indicated influenza strains at a MOI of 0.4 based on TCID50 in MDCK-SIAT1 cells. For the cells with cycloheximide added to block protein expression (and hence secondary transcription), cycloheximide was added to a final concentration of 50 *μ*g/ml (a concentration sufficient to block secondary transcription (Killip et al., 2014)) at the time of infection. After 8 hours, mRNA was harvested using the CellAmp Direct RNA Prep Kit for RT-PCR, reverse-transcribed using an oligo-dT primer, and qPCR was performed as described above.

For the qPCR in Figure 7B, A549 cells were infected with the NS1stop virus at a MOI of 0.1 (based on TCID50) in MDCK-SIAT1 and with the PB1del385to2163 virus at a dose that induced equivalent IFN in the absence of ribavirin. The ribavirin was added to cells at the time of infection to a final concentration of 100*μ*M, which is sufficient to block secondary transcription (Vanderlinden et al., 2016; Reuther et al., 2015; Scholtissek, 1976). All other steps were performed as for the cycloheximide experiments described immediately above.

### Viruses and experiments for validation experiments

In Figure 6, we tested the IFN inducing capacity of a variety of viral mutants identified in the single-cell experiments. For point-mutant viruses, we created variants for all amino-acid substitutions found in PB1 and NS among IFN+ cells that did not also lack NS. One of these mutants (amino-acid substitution S704P in PB1) did not reach sufficient titers in a single attempt to generate it by reverse genetics, and so was dropped from the experiment (note that we did not attempt replicates of the reverse genetics for this mutant, and so are *not* confident in drawing strong conclusions about its actual attenuation). This left six point-mutant viruses: four with point mutations in PB1, and two with point mutations in NS. We also created a mutant virus that contained the internal deletion in PB1 found in an IFN+ cell. In addition, we created a virus with an inactivated NS1 to mimic the infections that failed to express NS (we were unable to use complementing cells to generate a viral stock that completely lacked the NS segment). This NS1stop virus contained six nucleotide changes resulting in the addition of five in-frame stop codons in NS1 starting 10 nucleotides downstream of the 5’ splice donor site, thereby disrupting NS1 while leaving NS2 (NEP) intact. All of these mutants were cloned into the pHW2000 bi-directional reverse-genetics plasmid (Hoffmann et al., 2000) in order to enable generation of viruses encoding the mutant genes. File S1 provides the full sequences for all of these plasmids.

We generated the wild-type and point-mutant viruses for the validation experiments in Figure 6A by reverse genetics using the pHW18* series of WSN reverse genetics plasmids (Hoffmann et al., 2000), but substituting the appropriate mutant plasmid listed in File S1 for the wild-type plasmid for that gene. To generate the viruses from these plasmids, we transfected an equimolar mix of all eight plasmids into co-cultures of 293T and MDCK-SIAT1 cells seeded at a ratio of 8:1. At 24 hours posttransfection, we changed media from D10 to influenza growth media. At 50 hours post-transfection (for the replicate 1 viruses in Figure S6A) or 72 hours (for the replicate 2 viruses in Figure S6), we harvested the virus-containing supernatant, clarified this supernatant by centrifugation at 300×g for 4 min, and stored aliquots of the clarified viral supernatant at -80° C. We then thawed aliquots and titered by TCID50 on MDCK-SIAT1 cells. For the infections in Figure S6A, we wanted to use equivalent particle counts, so we normalized all viruses to an equivalent hemagglutination titer on turkey red blood cells (Hirst, 1942). Briefly, a solution of 10% v/v red blood cells (LAMPIRE Biological Laboratories, Fisher Scientific catalogue number 50412942) was washed in PBS and diluted to a final concentration of 0.5% v/v. Two-fold serial dilutions of virus were added to an equal volume of diluted red blood cells, and titer was measured as the highest dilution of viral stock at which complete hemagglutination of red blood cells was observed. We then performed infections of the A549 reporter cell line at equivalent hemagglutination titer and analyzed the data as described in Figure S6A.

To generate the NS1stop mutant virus and the wild-type and PB1del385to2163 mutant viruses in Figure S6B, we used slightly different procedures. The wild-type virus was generated by reverse genetics as described for the point-mutant viruses above, harvested at 48 hours post-transfection, and then passaged on MDCK-SIAT1 cells for 36 hours at an MOI of 0.05—conditions that we previously validated to lead to relatively little accumulation of defective particles (Russell et al., 2018). The NS1stop virus was similarly generated, but was passaged for 48 rather than 36 hours, since it had slower growth kinetics and so needed a longer period of time to reach high titers. The viruses with deletions in the PB1 segment could not be generated in normal 293T and MDCK-SIAT1 cells, since they required the exogenous expression of the PB1 protein. Therefore, these viruses were generated in previously described 293T and MDCK-SIAT1 cells that had been engineered to constitutively express PB1 (Bloom et al., 2010). These viruses were harvested from transfections at 72 hours, and passaged twice in the MDCK-SIAT1 cells constitutively expressing PB1 at a MOI of 0.001 for 72 hours and 0.01 for 48 hours. This passaging was necessary as viral titers from transfections were too low to generate sufficient virus from a single passage. The wild-type and NS1stop viruses were titered by TCID50 on MDCK-SIAT1 cells, and the PB1 deletion viruses were titered on the MDCK-SIAT1 cells constitutively expressing PB1. The infections in Figure S6B were performed at equivalent TCID50s as described in the legend to that figure. That these equivalent TCID50s were also roughly equivalent in terms of particles capable of undergoing primary transcription is shown in Figure S6C.

### Computational analysis of single-cell transcriptomic and viral sequence data

A computational pipeline that performs all steps in the data analysis is available at https://github.com/jbloomlab/IFNsorted_flu_single_cell. This pipeline is orchestrated by Snakemake (Koster and Rahmann, 2012), and begins with the raw sequencing data and ends by generating the figures shown in this paper. The sequencing data and annotated cell-gene matrix are available on the GEO repository under accession GSE120839 (https://www.ncbi.nlm.nih.gov/geo/query/acc.cgi?acc=GSE120839).

Briefly, the raw deep sequencing data from the Illumina 3’-end sequencing were processed using the 10X Genomics software package `cellranger` (version 2.2.0). We built a multi-species alignment reference consisting of a concatenation of the human and influenza virus transcriptomes (the first “species”) and the canine transcriptome (the second “species”). The human transcriptome was generated by filtering genome assembly GRCh38 for protein-coding genes defined in GTF file GRCh38.87. The influenza virus transcriptome consisted of the mRNAs for the wild-type A/WSN/1933 virus strain in File S1 (the `cellranger` alignment is sufficiently permissive that it aligns sequences from both the wild-type and synonymously barcoded viral variants to this transcriptome). The canine transcriptome was generated by filtering genome assembly CanFam3.1 for protein-coding genes defined in GTF file CanFam3.1.87. The `cellranger` software was used to align the Illumina 3’-end sequencing reads to this multi-species transcriptome, call human+influenza and canine cells (Figure 3A), and generate a matrix giving the expression of each gene in each single cell. We used a custom Python script to determine the number of influenza virus reads that could be assigned to the wild-type or synonymously barcoded virus, and added this information to the annotated the cell-gene matrix.

The PacBio sequences of the full-length viral genes were analyzed as follows. First, we used version 3.1.0 of PacBio’s ccs program (https://github.com/PacificBiosciences/unanimity) to build circular consensus sequences (CCSs) from the subreads files, requiring at least 3 passes and a minimum accuracy of 0.999. We further processed these CCSs using custom Python code and the minimap2 (Li, 2018) long-read aligner (version 2.11-r797). The Python code has been implemented in the API of dms_tools2 (https://jbloomlab.github.io/dms_tools2/ (Bloom, 2015)) package (version 2.3.0). A Jupyter notebook that performs these analyses is at https://github.com/jbloomlab/IFNsorted_flu_single_cell/blob/master/pacbio_analysis.ipynb, and is also provided in HTML form as File S5. We refer the reader to this notebook for a detailed description and extensive plots showing the results at each step. Here is a brief summary: we filtered for CCSs that had the expected 5’ termini (from the influenza-specific primers) and 3’ termini (corresponding to the 10X adaptor), and for which we could identify the cell barcode, UMI, and polyA tail. We aligned the cDNAs flanked by these termini to the influenza transcriptome, and performed a variety of quality control steps. At this point, we examined whether cDNAs had the synonymous viral barcodes at both ends or neither end as expected in the absence of strand exchange, and reassuringly found that strand exchange was rare (Figure S4). The small number of CCSs with identifiable strand exchange were filtered from further analysis. We then further filtered for CCSs that contained valid cell barcodes as identified by the `cellranger` pipeline, and kept just one CCS per UMI (preferentially retaining high-quality CCSs that aligned to full-length cDNAs). We then removed from the CCSs the barcoding synonymous mutations that we had engineered into one of the two viral variants. Finally, we used the CCSs to call the sequence of the viral gene in each cell, calling mutations separately for each viral barcode variant. We called mutations (insertions, deletions, and substitutions) in the viral gene sequences as follows:

1. Mutations with accuracies less than 0.999 (which constitute <0.5% of all mutations) were ignored.
2. If all CCSs for a particular viral-barcode variant of a gene in a cell were wild-type, it was called as wild type.
3. If any CCSs for a particular viral-barcode variant of gene in a cell had a mutation, then require at least two CCSs to call the sequence.
4. If at least two and >30% of the CCSs had a specific mutation, then call that mutation as present and note its frequency among the CCSs. The exception was single-nucleotide indels in homopolymers, for which we required three CCSs to call a mutation (the reason is that the main mode of PacBio sequencing errors is short indels in homopolymers).

The plots in https://github.com/jbloomlab/IFNsorted_flu_single_cell/blob/master/pacbio_analysis.ipynb or File S5 indicate that these are reasonable mutation-calling criteria. We could call the sequences of all expressed viral genes in about half of the infected cells (Figure S5). The mutations called using this pipeline are shown in Figure 4, and File S3 gives the number of CCSs supporting each mutation call. The called sequences of the viral genes were added to the annotated cell-gene matrix.

Finally, we process the annotated cell-gene matrix in R to generate the plots shown in this paper. This analysis utilized a variety of R and Bioconductor (Huber et al., 2015) packages, including Monocle (Qiu et al., 2017; Trapnell et al., 2014) and ggplot2. A Jupyter notebook that performs these analyses is at https://github.com/jbloomlab/IFNsorted_flu_single_cell/blob/master/monocle_analysis.ipynb, and is also provided in HTML form as File S6. We refer the reader to this notebook for a detailed description and a variety of additional plots not included in the paper. Briefly, we first filtered cells that were extreme outliers in the amount of mRNA as shown in Figure 3B. We used the uninfected canine cells to estimate the percentage of total mRNA in a cell that would come from influenza purely due to background (e.g., from cell lysis) in the absence of infection, and called as infected the human cells for which significantly more than this amount of mRNA was derived from influenza under a Poisson model (Figure 3C). We next used a Poisson model parameterized by the amount of expected background mRNA for each influenza gene to call the presence or absence of each influenza gene in each infected cell (Figure 3D and Figure S3A). To identify cells that were co-infected with both viral barcodes (Figure 3F), we used a binomial test to identify cells for which we could reject the null hypothesis that at least 95% of viral mRNA was derived from the more common viral barcode. We called IFN+ and ISG+ cells using the heuristic thresholds shown in Figure 3G and Figure S3C,D, respectively. We counted IFN mRNAs as any IFN-*α*, IFN-*β*, or IFN-*λ* transcripts, since type I and type III IFN were highly correlated (Figure S3B). We counted ISG mRNAs as any of CCL5, IFIT1, ISG15, or Mx1. The plot in Figure 4 summarizes all of the genotypic information, and was created in substantial part using `gggenes` (https://github.com/wilkox/gggenes).

## SUPPORTING INFORMATION

**Figure S1.**
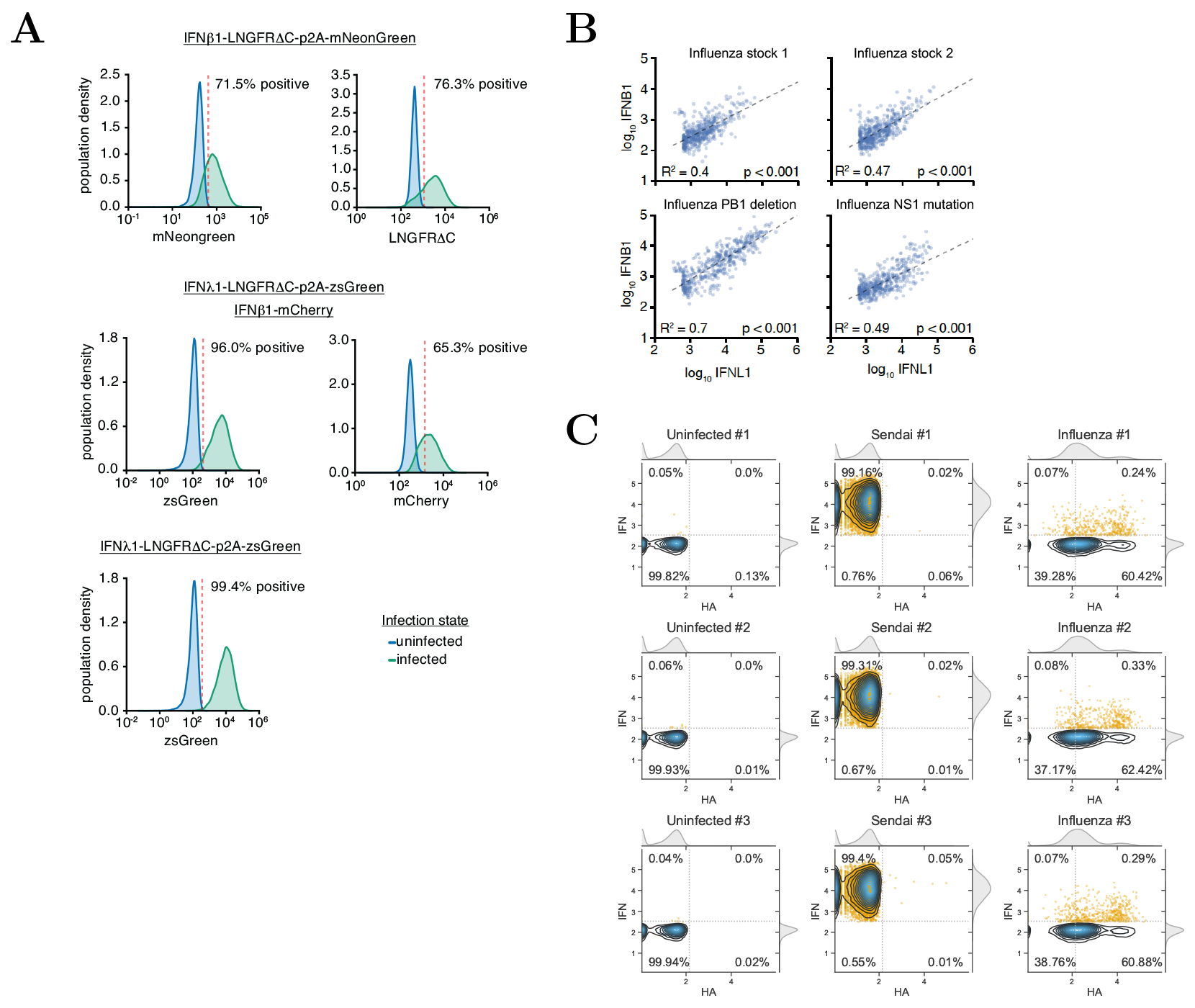
Validation of IFN reporter, related to Figure 1. (**A**) IFN reporter cell lines were infected at high MOI with the Cantell strain of Sendai virus (Strahle et al., 2006). The name of the reporter is indicated above the plots. At 13 hours post-infection, activation of the reporter was monitored by flow cytometry using the marker indicated below each plot (a fluorescent protein or antibody staining for the LNGFRΔC using PE-conjugated anti-LNGFR antibody from Miltenyi Biotec). Sendai infection efficiently activated the IFN reporter in all cases. (**B**) The type I and type III IFN reporters are highly correlated in their activation. An A549 cell line was generated by transduction with both IFN-β and IFN-λ reporters driving mCherry and ZsGreen, respectively. The cells were infected with two different stocks of WSN influenza, or stocks with a deletion in PB1 or stop codons in NS1. After 13 hours, cells were analyzed by flow cytometry. Cells positive for either reporter were further analyzed. As shown in the FACS plots, expression of the *IFNB1* and *IFNL1* reporters is highly correlated. (**C**) Flow cytometry data for Figure 1B. A549 cells with the *IFNL1* reporter driving LNGFRΔC-ZsGreen were not infected, infected with saturating amounts of the Cantell strain of Sendai virus, or infected at a target MOI of 0.3 with the stock of influenza virus used in the single-cell experiment. At 13 hours, cells were analyzed by FACS for HA and ZsGreen expression. Each condition was done in triplicate. Contour plots show density of all cells, and IFN+ cells are also indicated by orange dots. Cells were classified as HA+ or IFN+ based on gates set to put 0.05% of the uninfected cells in these populations. For influenza-infected cells, the percentage IFN+ was calculated only among HA+ cells (since these are the ones that are infected). For uninfected and Sendai-infected cells, the percentage IFN+ was calculated among all cells, since the cells do not express HA.

**Figure S2.**
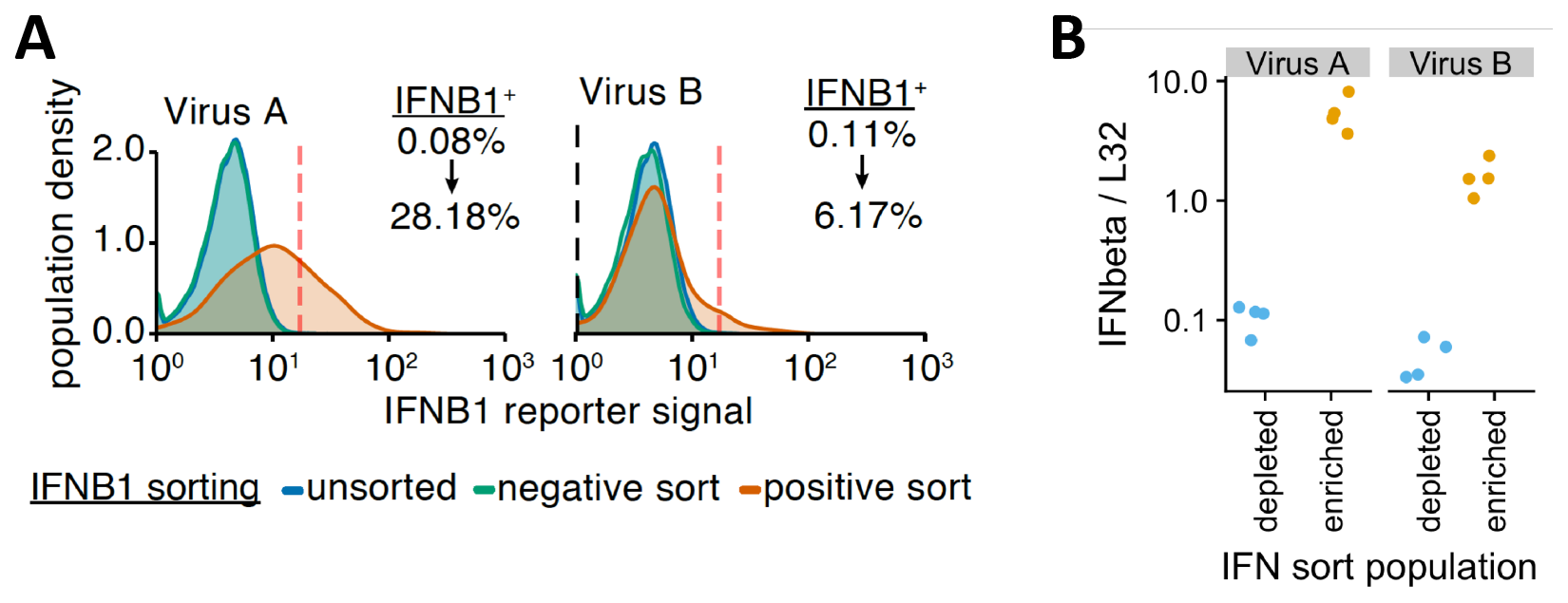
Example MACS enrichments of IFN+ influenza-infected cells, related to Figure 2. A549 cells with the *IFNB1* LNGFRΔC-mNeonGreen reporter were infected with wild-type WSN influenza (two different viral stocks) at a target MOI of 0.1 TCID50 per cell. After infection had proceeded for 12 hours, the cells were twice magnetically sorted for LNGFRΔC expression over magnetic columns as detailed in the methods for the single-cell sequencing experiment. (**A**) After sorting, the populations were analyzed by flow cytometry for IFN expression using the mNeonGreen fluorescent protein. The plots show the distribution of fluorescence in the original population, the flow-through from the first column, and the MACS-sorted positive population after two columns. As indicated by the percentages shown for the original and MACS-sorted population, this process led to substantial enrichment in IFN+ cells. We expect that the IFN sorting for the actual single-cell sequencing led to similar enrichment, although we could not directly quantify this as the sorted cells in that case were immediately used for the sequencing and so could not be analyzed by flow cytometry. (**B**) Analysis of expression of IFNB1 (relative to the housekeeping gene L32) by qPCR in the positive (IFN enriched) and negative (IFN depleted) populations from panel (A). The qPCR validates a roughly 50- to 100-fold enrichment in total IFNB1 expression. The qPCR was performed in quadruplicate (hence the four points for each sample).

**Figure S3.**
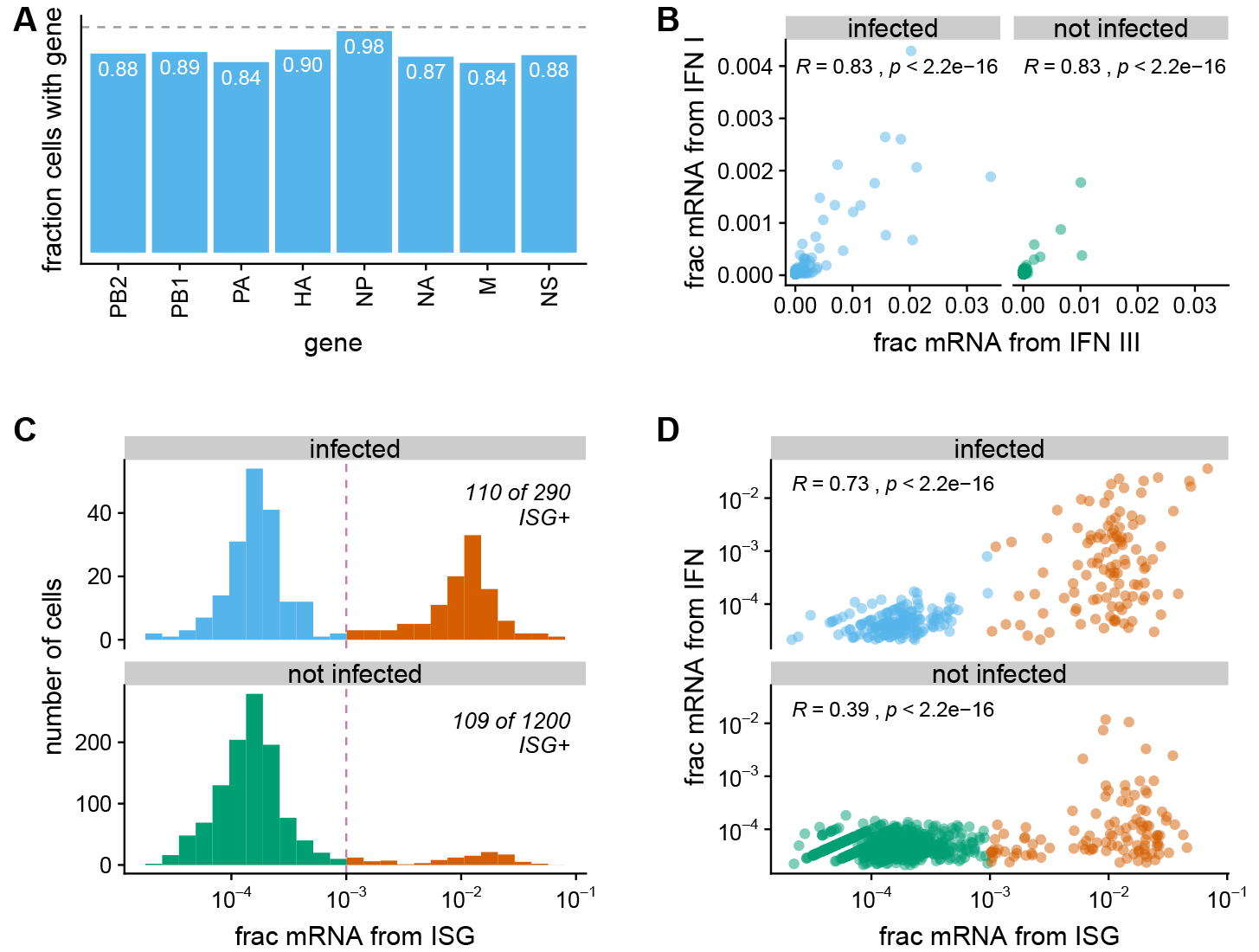
Additional plots from analysis of transcriptomic data, related to Figure 3. (**A**) Fraction of infected cells that are called as expressing each viral gene. The gray dashed line is at one. Each gene is detected in ~80-90% of cells, roughly in line with prior estimates (Brooke et al., 2013; Heldt et al., 2015; Dou et al., 2017; Russell et al., 2018). The exception is NP, which is detected in virtually all infected cells. The higher frequency of detecting NP could reflect a biological phenomenon, but we suspect it is more likely that cells lacking NP tend to have lower viral gene expression and so are not reliably called as infected in our experiments because the number of viral mRNAs is below the detection limit. (**B**) Correlation between the fraction of cellular mRNA derived from type I and type III IFN in the A549 cells in our single-cell transcriptomics. Each point represents one cell. The plots are faceted by whether the cells are called as infected, and the Pearson correlation coefficient is shown. (**C**) Distribution of ISGs expression across single infected and uninfected cells. For each cell, we quantified ISG expression as the total fraction of cellular mRNAs derived from four prototypical ISGs (IFIT1, ISG15, CCL5, and Mx1). We heuristically classify as ISG+ cells with > 10^-3^ of their cellular mRNA from these ISGs, and color these cells red. Comparison to Figure 3G shows that more cells are ISG+ than IFN+, probably because paracrine signaling can induce ISG expression in cells that do not themselves express IFN (Stetson and Medzhitov, 2006; Honda et al., 2006). (**D**) Correlation between the fraction of cellular mRNA derived from IFN and ISGs. Each point is one cell, and the Pearson correlation coefficient is shown. IFN and ISG expression are more correlated for infected than uninfected cells, probably because in the latter the ISG expression is more often due to paracrine signaling that does not induce expression of IFN itself. Among both the infected and uninfected populations, there are many cells that express ISGs but not IFN, but no cells that express IFN but not ISGs.

**Figure S4.**
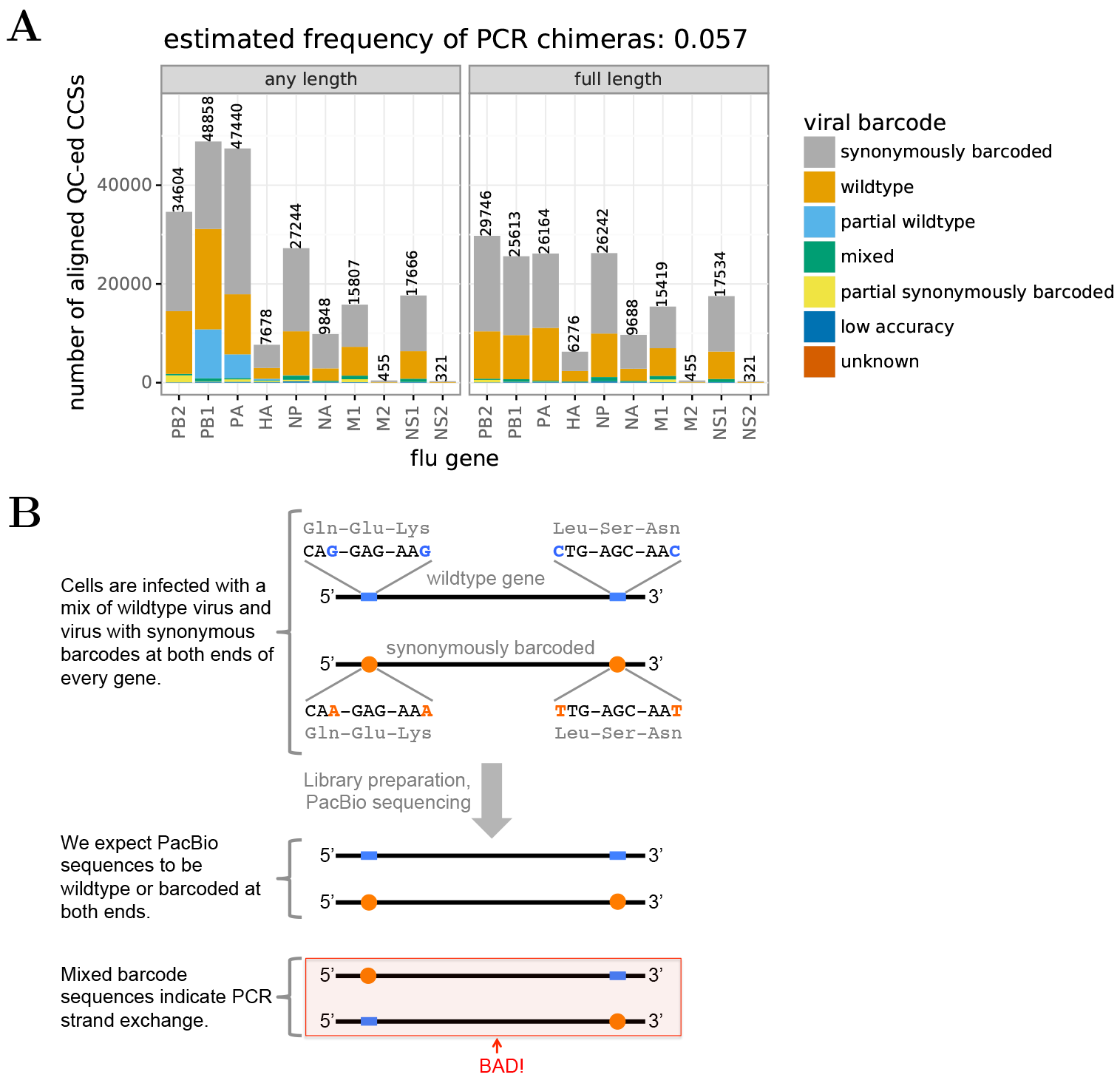
PacBio CCS quality control, related to Figure 4. **(A)** Number of PacBio CCSs that passed quality-control and aligned to each viral gene. CCSs were obtained using several PacBio runs which were loaded with different amounts of each gene to balance coverage (see File S5). Therefore, unlike the transcriptomic data in Figure 3, the numbers of CCSs is *not* an indicator of a transcript’s abundance. Especially for the polymerase genes, many CCSs corresponded to genes with internal deletions, since these shorter forms are preferentially amplified by PCR. Therefore, the plot is faceted by CCSs for any length of the gene, and for full-length genes. The disproportionate sequencing of the shorter internally deleted genes does not affect the genotype calling since UMIs were used to collapse sequences from the same cDNA, and cell barcodes were used to collapse sequences from the same cell. The bars in the plot are colored by whether the sequence is derived from the wild-type viral variant, the synonymously barcoded viral variant, or is a mixed-barcode molecule (see panel B). We estimate (Bloom, 2018) that 5.7% of molecules are chimeric due to PCR strand exchange. About half of these chimeras can identified by the mixed viral barcodes and removed from subsequent analyses, leaving ~3% un-identified chimeras. For some molecules (mostly polymerase genes with internal deletions) one of the barcode sites was deleted from the molecule and so the viral barcode could only be partially called. (**B**) Strategy for detecting strand exchange in PacBio sequencing. Preparation for PacBio sequencing required many PCR cycles, which can cause strand exchange that scrambles mutations and 10X cell barcodes / UMIs from different molecules. If there is no strand exchange, all molecules are either wild-type or have the synonymous barcoding mutations at *both* termini. Strand exchange creates molecules with wild-type nucleotides at one termini and synonymous barcoding mutations at the other. Panel A shows the frequencies of these molecules. Since homologous recombination in influenza virus in negligible (Boni et al., 2008), mixed-barcode molecules are *not* generated naturally during co-infection.

**Figure S5.**
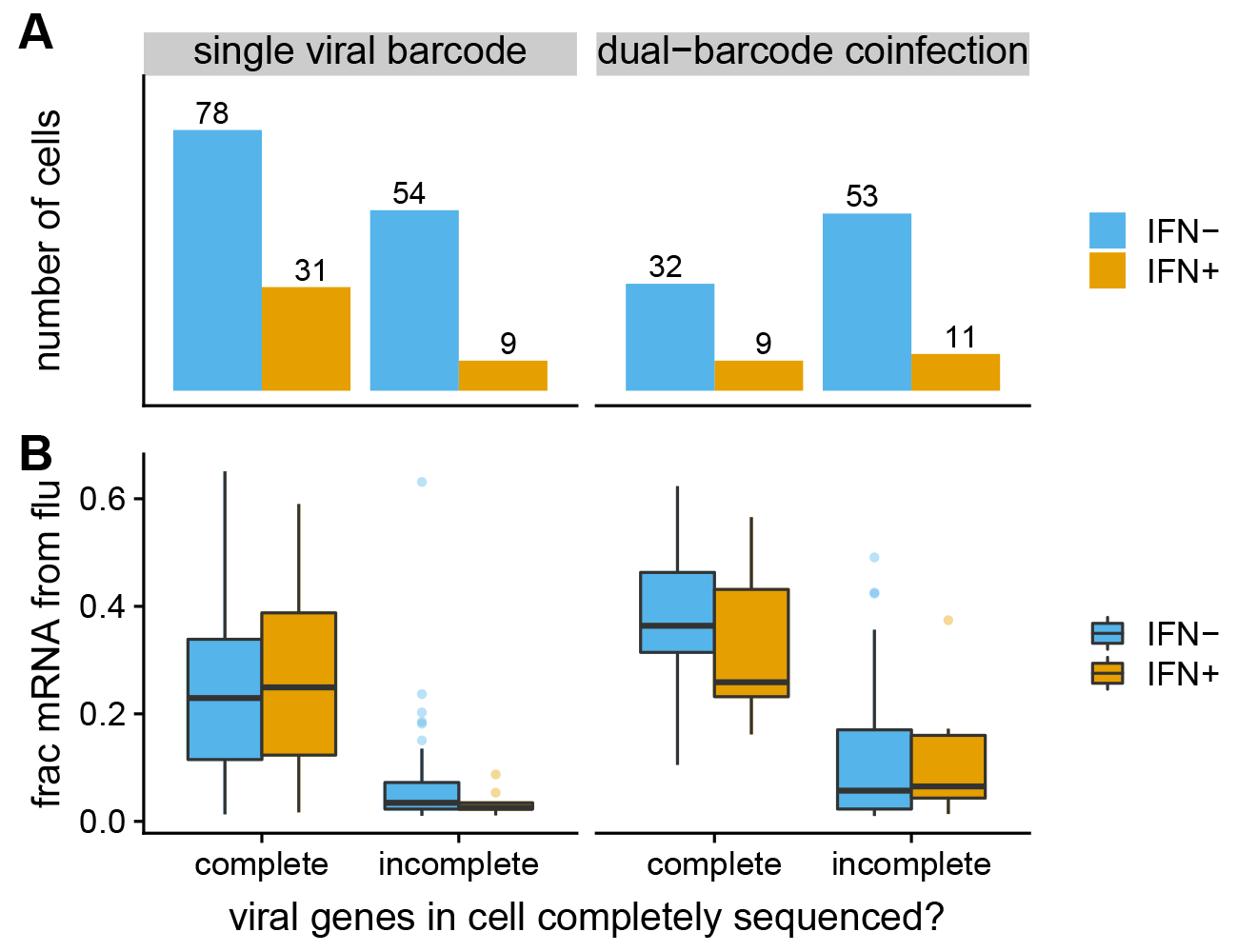
Number of cells for which we could determine the full sequences of all genes expressed by the infecting virion(s). (**A**) We could call the complete genotypes of the infecting virion(s) for the majority of cells infected with just a single viral barcode variant, but only a minority of cells co-infected with both viral barcodes. (**B**) The cells for which we could call complete viral genotypes tended to have higher expression of viral mRNA than cells for which we could not call complete genotypes. Both facts makes sense. Cells with more viral mRNA are more likely to have their viral cDNA captured in the PacBio sequencing, which is only captures a small fraction of the total transcripts identified by the 3’-end sequencing transcriptomic sequencing. The lower calling rate for dual-barcode co-infections is probably because these co-infections have more viral genes that must be sequenced (potentially a copy of each viral gene from each viral variant), increasing the chances that one of these genes is missed by the PacBio sequencing. An important implication of this plot is that the cells for which we call complete viral genotypes are *not* a random subsampling of all infected cells in the experiment, but are rather enriched for cells that have high levels of viral mRNA and do not have dual-barcode viral infections. Note also that this plot is limited to the cells that were called as infected (Figure 3C) and could clearly be classified as IFN- or IFN+ (Figure 3G).

**Figure S6.**
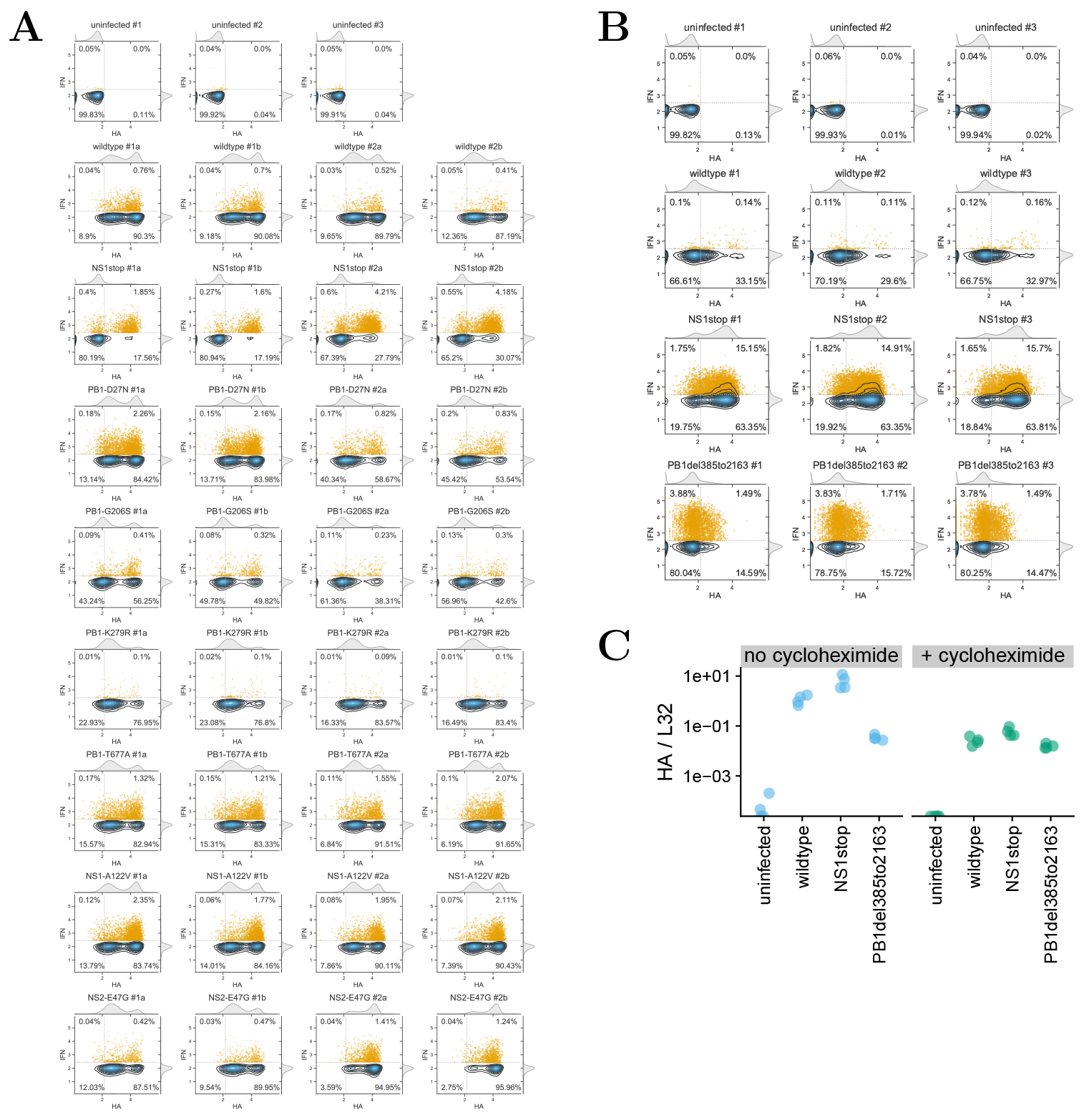
Additional data for validation experiments, related to Figure 6. (**A**) Flow cytometry data for Figure 6A. Data were collected and analyzed as in Figure S1C. For each viral mutant, two independent stocks were assayed in duplicate (#1a and #1b are one viral stock; #2a and #2b are the other). Infections with replicate #1 of wild-type virus were performed at an MOI of 0.1 as determined by TCID50, and other viruses were infected at an equivalent particle number as determined by HI assay. (**B**) Flow cytometry data for Figure 6B. The virus with the deletion in PB1 cannot be normalized by HA expression since it expresses less HA due to the lack of secondary transcription. Therefore, cells were infected at a MOI of 0.3 as determined by TCID50 on MDCK-SIAT1 cells for wild type and NS1stop, and on MDCK-SIAT1 cells expressing PB1 (Bloom et al., 2010) for PB1del385to2163. Panel C shows that at these TCID50s, all variants had similar amounts of transcriptionally active virus in the absence of secondary transcription. The percent IFN+ was calculated for *all* cells (HA+ and HA-) since that is a more fair comparison for PB1del385to2163. (**C**) Validation that infections in panel B were performed at similar doses of virions capable of initiating primary transcription. A549 cells were infected at MOI of 0.4 (based on TCID50 as described panel B), and after 8 hours mRNA was harvested for qPCR on oligo-dT primed reverse-transcription products. The y-axis shows the ratio of viral HA mRNA to the housekeeping gene L32. Infections were performed in the presence of absence of 50 *μ*g/ml cycloheximide, which blocks protein synthesis and hence secondary transcription (Killip et al., 2014). In the absence of cycloheximide, viruses with deletions in PB1 produced less viral mRNA because they could not produce PB1 protein for secondary transcription. But in the presence of cycloheximide, all viruses produced similar amounts of viral mRNA, indicating that the dose of particles active for primary transcription is roughly equivalent across variants. Each measurement was performed in quadruplicate.

**Figure S7.**
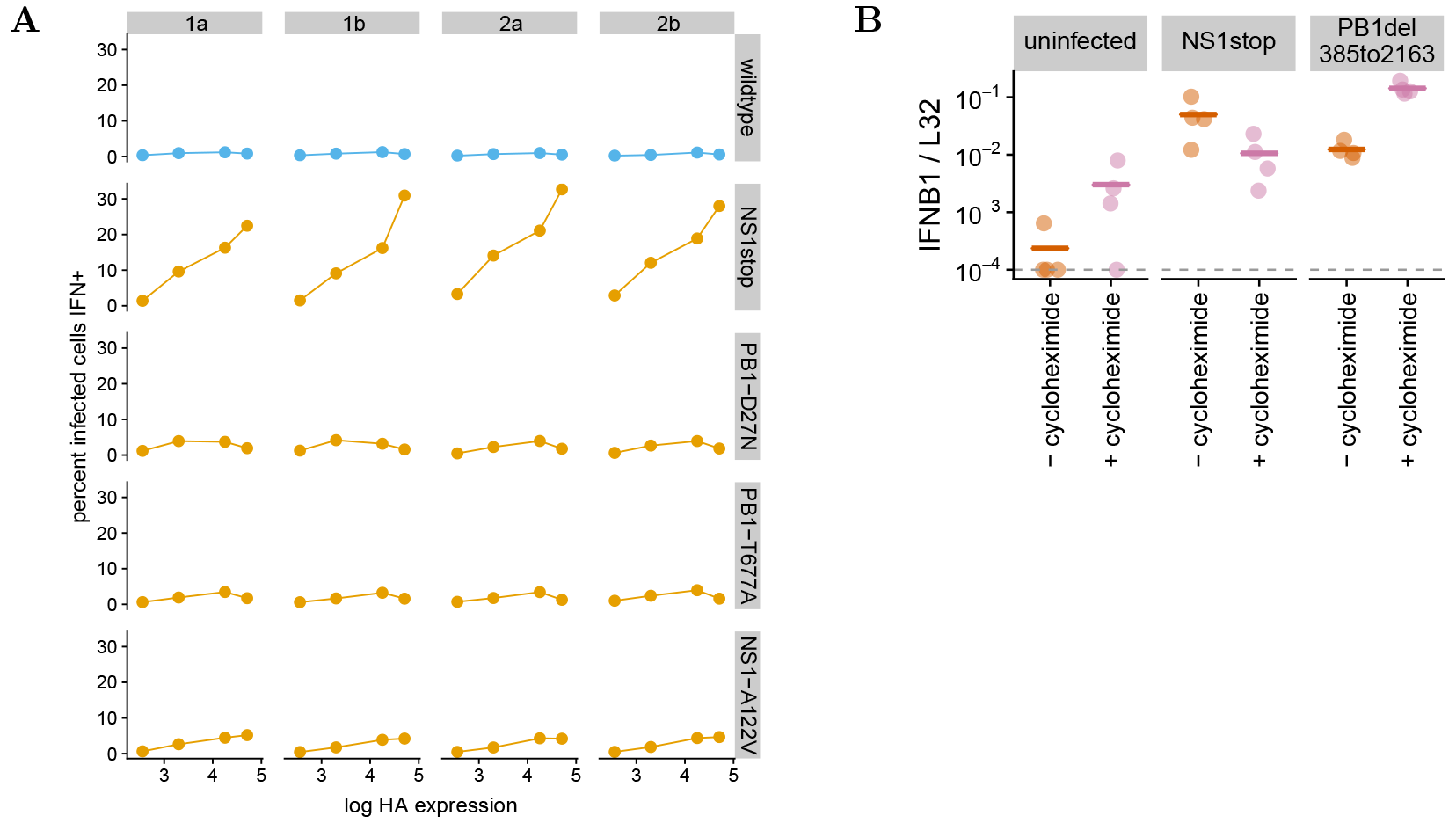
Additional data for comparison of IFN induction as a function of viral load, related to Figure 7. (**A**) Detailed analysis of the relationship among HA expression and IFN induction for different viral mutants that is summarized in Figure 7A. We re-analyzed all the infected cells in Figure S6A (defining “infected” using the same HA-gating scheme in Figure S6A) for the wild-type virus, the NS1stop virus, and the three amino-acid mutants with significantly increased IFN induction. For each virus and replicate, we binned the infected cells into HA expression quartiles. We then calculated the percent of cells that were IFN+ in each quartile (defining “IFN+” using the same gating scheme in Figure S6A). The plots show the mean HA expression in each quartile versus the percent of cells that are IFN+. The results clearly indicate that for the NS1stop virus, increased viral burden (more HA) correlates with IFN induction. A similar trend also holds for the virus with an amino-acid substitution in NS1 (NS1-A122V). But for wild type and the two PB1 mutants, there is no clear trend for higher viral burden to correlate with IFN induction. Figure 7A summarizes the data in this figure by simply showing the ratio of the percent IFN+ of the highest HA-expression quartile to the lowest HA-expression quartile for each sample and replicate. (**B**) Similar to Figure 7B, but uses cycloheximide rather than ribavirin. Preventing viral secondary transcription using cycloheximide also decreases IFN induction by the NS1stop virus relative to the PB1 deletion virus, corroborating the result in Figure 7B with a different inhibitor. However, the results are more complex here, since cycloheximide alone increases IFN production, even in response to non-viral challenges such as poly(I:C) (Raj and Pitha, 1981; Maroteaux et al., 1983; Ringold et al., 1984; Killip et al., 2014). Therefore, we see an increase in IFN levels upon cycloheximide treatment for uninfected cells and cells infected with the PB1 deletion virus, but a decrease for cells infected with the NS1stop virus. This plot shows qPCR results for the same samples as in Figure S6C, but with the qPCR for expression of IFNB1 transcripts relative to the housekeeping gene L32. As described in Figure S6C, A549 cells were infected at an MOI of 0.4, such that HA expression was equivalent between viral variants when secondary transcription was blocked, and some of the samples treated with 50 *μ*g/ml of cycloheximide (a concentration sufficient to block secondary transcription; Killip et al., 2014). After 8 hours, transcript levels were quantified by qPCR. Points indicate four replicates, and lines indicate the mean. The dashed line shows the limit of detection, and points below this limit where set to that value.

**File S1. Sequences of plasmids and viruses used in this study.** The plasmid maps are for the IFN reporters in Figure 1A and the mutant viral genes cloned into the pHW* bi-directional reverse genetics plasmid (Hoffmann et al., 2000). In addition, there are Genbank files giving the sequences of the wild-type and synonymously barcoded WSN viruses. A README file explains the contents of the ZIP.

**File S2. A text file giving the primers used to amplify the influenza cDNAs for PacBio sequencing.**

**File S3. A CSV file giving the genotypes in Figure 4.**

**File S4. A CSV file giving the viral mutations and related information in Figure 5.**

**File S5. HTML rendering of Jupyter notebook that analyzes the PacBio data.** The actual Jupyter notebook is at https://github.com/jbloomlab/IFNsorted_flu_single_cell/blob/master/pacbio_analysis.ipynb.

**File S6. HTML rendering of Jupyter notebook that analyzes the annotated cell-gene matrix.** The actual Jupyter notebook is at https://github.com/jbloomlab/IFNsorted_flu_single_cell/blob/master/monocle_analysis.ipynb.

## Notes

#### Summary of Updates

This is a modestly revised version of the original pre-print we posted, with modest textual changes and one additional figure panel added in the last section of the Results. These changes are based on feedback that we received on the earlier version.

