## Supplementary material for "Single-cell virus sequencing of influenza infections that trigger innate immunity": File S1: README.md

# Genbank sequences of plasmids and viruses

All files are in Genbank file format.

## Plasmid maps for IFN reporters

The reporters are:

 - [1517_pHAGE2_IFNbeta_prom_LNGFR_P2A_mNeongreen.gb](1517_pHAGE2_IFNbeta_prom_LNGFR_P2A_mNeongreen.gb)

 - [1767_pHAGE2_IL29_promnkozak_LNGFR_P2A_zsGreen.gb](1767_pHAGE2_IL29_promnkozak_LNGFR_P2A_zsGreen.gb)

## Plasmid maps for viral mutants
The numbers in the plasmid names indicate their numbers in the Bloom lab plasmid log.

Nucleotide mutations are numbered in the reverse-sense orientation of vRNA (so that mRNA is in positive sense) in sequential 1, 2, ... numbering.
Amino-acid mutations are numbered in sequential in 1, 2, ... numbering as well.

 - [1481_pHH21-WSN-HAflank-startmut-mCherry.gb](1481_pHH21-WSN-HAflank-startmut-mCherry.gb): encodes HA gene segment with mCherry in place of the HA coding sequence. There are flanking packaging regions of HA around the mCherry, with start codons that would be upstream of mCherry mutated away.

 - [1651_pHW182-PB1-d385-2163.gb](1651_pHW182-PB1-d385-2163.gb): encodes PB1 from WSN strain with nucleotides 385 to 2163 (inclusive) deleted.

 - [1717_pHW188_NS1mut.gb](1717_pHW188_NS1mut.gb): encodes NS gene from WSN strain of influenza with point mutations that create 5 stop codons shortly downstream of first splice site. This will disrupt NS1, but should leave NS2 intact.

 - [2278_pHW182-PB1_A860G.gb](2278_pHW182-PB1_A860G.gb): encodes PB1 from WSN strain with nucleotide mutation A860G, which introduces amino-acid change Lys279Arg in PB1.

 - [2281_pHW188-NS_A638G.gb](2281_pHW188-NS_A638G.gb): encodes NS from WSN strain with nucleotide mutation A638G, which introduces Glu47Gly in NS2.

 - [2288_pHW182-PB1_G103A.gb](2288_pHW182-PB1_G103A.gb): encodes PB1 from WSN strain with nucleotide mutation G103A, which introduces Asp27Asn in PB1.

 - [2289_pHW182-PB1_G640A.gb](2289_pHW182-PB1_G640A.gb): encodes PB1 from WSN strain with nucleotide mutation G640A, which introduces amino-acid mutation Gly206Ser in PB1.

 - [2290_pHW182-PB1_A2053G.gb](2290_pHW182-PB1_A2053G.gb): encodes PB1 from WSN strain with nucleotide mutation A2053G, which introduces amino-acid mutation Thr677Ala in PB1.

 - [2292_pHW188-NS_C391T.gb](2292_pHW188-NS_C391T.gb): encodes NS from WSN strain with nucleotide mutation C391T, which introduces amino-cid mutation Ala122Val in NS1.

## Sequences of wildtype and synonymously barcoded WSN influenza
Genbank files giving the sequences of the viral RNAs (in the orientation of the mRNA, so reverse complement of actual negative-sense viral RNA) for the wildtype and synonymous barcoded WSN influenza variants:

 - [flu-wsn.gb](flu-wsn.gb): viral RNAs for wild-type WSN influenza.

 - [flu-wsn-double-syn.gb](flu-wsn-double-syn.gb): viral RNAs for synonymously barcoded WSN influenza.
