## Supplementary material for "Single-cell virus sequencing of influenza infections that trigger innate immunity": File S5

pacbio\_analysis


### Table of Contents

- Analysis of full-length PacBio sequencing of influenza mRNAs
  - Set up for analysis
    - Import Python modules
    - Define / create directories
  - Get the CCSs
    - Specify PacBio runs of interest
    - Read CCSs
    - Filter for CCS accuracy and passes
    - CCS accuracy / length
  - Align CCSs to flu genes
    - Barcoded CCS features
    - Flu alignment targets
    - Amount to trim termini
    - Un-circularize CCSs on circularized template
    - Match and align CCSs
    - Stats on matching / alignment
    - Get CCSs with gene aligned
  - Quality-control alignments
    - Barcode accuracy
    - Additional alignments
    - Excessive alignment trimming
      - Trimming of query start
      - Trimming of target start
      - Trimming query ends
      - Trimming target ends
    - Get QC-ed alignments
  - Summary of QC-ed alignments
    - Number of alignments per gene
    - Distribution of alignment lengths
    - Near full-length alignments
    - Passes/accuracy for alignments
  - Analyze viral barcodes
    - Inspect viral barcodes by gene
    - Estimate rate of PCR strand exchange
    - Figure with number CCSs and strand exchange
    - Filter chimeras, assign viral barcodes
  - 10X cell barcodes / UMIs
    - Get valid cell barcodes
    - Filter alignments from valid cells
    - Per-gene barcodes / UMI sampling
    - Get just one sequence per UMI
    - Genes sequenced per cell
  - Call mutations
    - Number of mutations per gene
    - Lengths of indels
    - Accuracy of mutations
  - Analyze flu sequences at cell level
    - Aggregate data at the cell level
    - Genes sequenced per cell
  - Call mutations for viruses in cells
    - Call consensus mutations
    - Frequency of mutations
    - Mutations vs number of sequences
  - Add to cell-gene matrix annotations

### Analysis of full-length PacBio sequencing of influenza mRNAs¶

The material from the 10X libraries for the *IFN\_enriched* sample was enriched for viral mRNAs by semi-specific PCR, and then sequenced by PacBio.
Here we analyze those data.

#### Set up for analysis¶

First, we do some things to set up this Jupyter notebook.

##### Import Python modules¶

We make extensive use of dms\_tools2 to handle the PacBio data.

We import plotnine for plotting using a ggplot2-like syntax.

In [1]:

```
import os
import re
import math
import glob
import subprocess
import shutil
import multiprocessing
import warnings
import collections

import numpy
import pandas
from IPython.display import display, HTML, Image
import Bio.SeqIO

# import plotnine for ggplot2 style plotting
from plotnine import *
_ = theme_set(theme_bw(base_size=12))

# ignore warnings that clutter output
warnings.simplefilter('ignore')

import dms_tools2
print("Using dms_tools2 version {0}".format(dms_tools2.__version__))
from dms_tools2.ipython_utils import showPDF
from dms_tools2.plot import COLOR_BLIND_PALETTE_GRAY
```

```
Using dms_tools2 version 2.3.0
```

##### Define / create directories¶

We define the names of key directories for input and output, and create these directories if needed:

In [2]:

```
# top results directory
resultsdir = './results/'

# directory with PacBio results
pacbioresultsdir = os.path.join(resultsdir, 'pacbio')

# directory with paper figures from this notebook
figsdir = 'paper/figures/pacbio_single_cell_figures/'
os.makedirs(figsdir, exist_ok=True)
```

#### Get the CCSs¶

We will work with the circular consensus sequences (CCSs) generated by the PacBio ccs program.

The actual generation of these CCSs is time consuming, and is performed by the commands in Snakefile.
This notebook assumes that those CCSs have already been generated.

##### Specify PacBio runs of interest¶

We read the list of PacBio runs for which we have generated CCSs from ./data/PacBio\_runs.tsv:

In [3]:

```
seqrunsfile = 'data/PacBio_runs.csv'

with open(seqrunsfile) as f:
    seqruns = [line.split(',')[0] for line in f]
    
print("Using the following PacBio sequencing runs:\n\t{0}".format('\n\t'.join(seqruns)))
```

```
Using the following PacBio sequencing runs:
	2017-06-08
	2017-12-07
	2018-06-22_nonPol
	2018-06-22_Pol-1
	2018-06-22_Pol-2
	2018-06-22_Pol_open
	2018-08-08_Pol_circ
```

Note that we have multiple sequencing runs:

- A run from June-8-2017 done at the UW PacBio core. This run used a PacBio RSII machine.
- A run from December-7-2017 done at the Fred Hutch Genomics core. This run used a PacBio Sequel machine.
- A series of runs done on June-22-2018 at the Fred Hutch Genomics Core using a PacBio Sequel machine. These runs were:
  - One that was run on a mix of all non-polymerase genes
  - Two that were run on a mix of just the polymerase genes
  - One that was run on a mix of just the polymerase genes that were amplified using an open PCR rather than an emulsion PCR, and for which a size-selection was performed to attempt to obtain more full-length genes.
- A run from August-8-2018 designed to target full-length polymerase genes in which the products were circularized and then linearized by PCR from the center of the genes.

The material sequenced for these various runs was **not** identical.
The relative amount of each viral gene differed across runs, as well as the size selection and the way that the libraries were loaded.
Therefore, we do not expect the same balance of different reads in the two runs.

##### Read CCSs¶

We now read each set of results into a dms\_tools2.pacbio.CCS object, taking only CCSs with at least the minimum accuracy indicated below:

In [4]:

```
ccsdir = os.path.join(pacbioresultsdir, 'ccs')

ccslist = []
for seqrun in seqruns:       
    reportfile = os.path.join(ccsdir, seqrun + '_report.csv')
    bamfile = os.path.join(ccsdir, seqrun + '_ccs.bam')
    ccslist.append(dms_tools2.pacbio.CCS(seqrun, bamfile, reportfile))
```

Here are the number of CCSs from each run:

In [5]:

```
zmw_plot = os.path.join(ccsdir, 'ZMW_plot.pdf')

ccs_report = dms_tools2.pacbio.summarizeCCSreports(
                ccslist, 'zmw', zmw_plot)
showPDF(zmw_plot)
```

As expected, we get fewer CCSs from the first run on the UW RSII machine, and then similar numbers of CCSs for all runs on the Sequel at the Hutch.
It consistently looks like about a third of ZMWs give successful CCSs.
When they're not successful, it's usually because there weren't enough passes.

##### Filter for CCS accuracy and passes¶

Only retain CCSs with accuracy of at least 99.9% and at least 3 passes on the CCS sequencing.
Depending on the settings used to run `ccs` in Snakefile, this may be all the CCSs.

In [6]:

```
ccsminaccuracy = 0.999
minpasses = 3

for ccs in ccslist:
    n_orig = len(ccs.df)
    ccs.df = ccs.df.query('(CCS_accuracy >= @ccsminaccuracy) & (passes >= @minpasses)')
    print("For {0} CCSs, {1} of {2} met the accuracy threshold of {3:.3f}"
            .format(ccs.samplename, len(ccs.df), n_orig, ccsminaccuracy))
```

```
For 2017-06-08 CCSs, 18063 of 18063 met the accuracy threshold of 0.999
For 2017-12-07 CCSs, 209930 of 209930 met the accuracy threshold of 0.999
For 2018-06-22_nonPol CCSs, 177109 of 177109 met the accuracy threshold of 0.999
For 2018-06-22_Pol-1 CCSs, 145507 of 145507 met the accuracy threshold of 0.999
For 2018-06-22_Pol-2 CCSs, 168907 of 168907 met the accuracy threshold of 0.999
For 2018-06-22_Pol_open CCSs, 133137 of 133137 met the accuracy threshold of 0.999
For 2018-08-08_Pol_circ CCSs, 126668 of 126668 met the accuracy threshold of 0.999
```

##### CCS accuracy / length¶

Now we plot the distributions of lengths, accuracies, and number of passes for the CCSs.
We do this on all of the CCSs combined rather than on the individual runs:

In [7]:

```
ccs_plot = os.path.join(ccsdir, 'ccs_plot.pdf')

dms_tools2.plot.plotColCorrs(
            pandas.concat([ccs.df for ccs in ccslist]), 
            ccs_plot,
            ['passes', 'CCS_accuracy', 'CCS_length'])

showPDF(ccs_plot, width=700)
```

We see that most CCSs have high accuracy and plenty of passes, well above the thresholds used when running `ccs` in Snakefile.

#### Align CCSs to flu genes¶

The PacBio sequencing was performed on PCR-amplified product.
This PCR product was amplified off the 10X barcoded material using semi-specific PCR with one end specifically annealing to flu transcripts.
We want to identify the CCS's that represent properly barcoded 10X material from influenza genes, and then call the barcodes and align the barcoded mRNA to influenza transcripts.

##### Barcoded CCS features¶

This 10X technical note outlines the sequences appended by the v2 10X single-cell 3' kit used in these experiments.
Specifically:

In [8]:

```
display(Image('./data/images/10Xschematic.png'))
```

Zooming in on the 3' adaptor sequence, it is:
`CTACACGACGCTCTTCCGATCT-NNNNNNNNNNNNNNNN-NNNNNNNNNN-TTTTTTTTTTTTTTTTTTTTTTTTTTTTTTVN` where the dash-delimited sequences are the read-1 priming site, the 16XN cell barcode, the 10XN UMI, the 30XT oligo-dT primer, followed by `V` (anything but `T`) and `N` (nominally, first nucleotide of upstream of mRNA polyA tail).

For the PCR to enrich the 10X product, Alistair used a 3' primer that anneals to this adaptor, namely:

In [9]:

```
primer3 = 'CTACACGACGCTCTTCCGATCT'
print("The length of the 3' primer is {0} nt".format(len(primer3)))
```

```
The length of the 3' primer is 22 nt
```

For the 5' primer, Alistair used a mix of primers that covered each of the 8 flu gene segments:

In [10]:

```
primer5_mix = {'fluPB2':'GCGAAAGCAGGTCAATTATATTCAATATGGAAAG',
               'fluPB1':'GCGAAAGCAGGCAAACCATTTG',
               'fluPA' :'GCGAAAGCAGGTACTGATTCAAAATGG',
               'fluHA' :'GCAAAAGCAGGGGAAAATAAAAACAACC',
               'fluNP' :'GCAAAAGCAGGGTAGATAATCACTCAC',
               'fluNA' :'GCGAAAGCAGGAGTTTAAATGAATCCAAAC',
               'fluM'  :'GCAAAAGCAGGTAGATATTGAAAGATGAGTC',
               'fluNS' :'GCAAAAGCAGGGTGACAAAGACATAATG',
              }
print("The length of the 5' primer ranges from {0} to {1} nt".format(
        min(map(len, primer5_mix.values())), max(map(len, primer5_mix.values()))))
```

```
The length of the 5' primer ranges from 22 to 34 nt
```

##### Flu alignment targets¶

We want to align the barcoded sequences to the flu mRNAs.
We will align to the wildtype flu sequences using a dms\_tools2.minimap2.Mapper, and then call the synonymous barcodes using a dms\_tools2.minimap2.TargetVariants object.
Both of these objects are initialized here.

The wildtype flu mRNA sequences are in ./data/flu\_sequences/flu-wsn-mRNA.fasta, and the synonymous barcoded ones are in ./data/flu\_sequences/flu-wsn-mRNA-double-syn.fasta.
We want to align to the sequence **interior** to the primer binding sites defined above: the region between the custom 5' primer and the polyA tail.
We therefore read in the mRNAs, and then trim each from the 5' termini to the end of the primer binding site:

In [11]:

```
# get the targets and synonymous-barcoded targets mRNAs
targets = collections.defaultdict(list)
for variant in ['', '-double-syn']:
    for full_mRNA in Bio.SeqIO.parse(
            './data/flu_sequences/flu-wsn{0}-mRNA.fasta'.format(variant), 'fasta'):
        full_mRNA_seq = str(full_mRNA.seq)
        primer = [p for p in primer5_mix.values() if full_mRNA_seq.count(p)]
        assert len(primer) == 1, "match multiple primers in {0}".format(full_mRNA.name)
        targetseq = full_mRNA_seq[full_mRNA_seq.index(primer[0]) + len(primer[0]) : ]
        targets[variant].append((full_mRNA.name, targetseq))
        if not variant:
            print("Aligning to {0} nts of {1}".format(len(targetseq), full_mRNA.name))

# write the targets to files
aligndir = os.path.join(pacbioresultsdir, 'alignments')
os.makedirs(aligndir, exist_ok=True)
targetfile = os.path.join(aligndir, 'targets.fasta')
with open(targetfile, 'w') as f:
    f.write('\n'.join('>{0}\n{1}'.format(*tup) for tup in targets['']))
syntargetfile = os.path.join(aligndir, 'targets-double-syn.fasta')
with open(syntargetfile, 'w') as f:
    f.write('\n'.join('>{0}\n{1}'.format(*tup) for tup in targets['-double-syn']))

# initialize mapper
mapper = dms_tools2.minimap2.Mapper(targetfile,
        dms_tools2.minimap2.OPTIONS_VIRUS_W_DEL,
        target_isoforms={'fluM1':['fluM2'], 'fluM2':['fluM1'],
                         'fluNS1':['fluNS2'], 'fluNS2':['fluNS1']})
print("\nAligning with minimap2 version {0}".format(mapper.version))

# initialize target variant caller
targetvariants = dms_tools2.minimap2.TargetVariants({'wildtype':targetfile,
        'synonymously barcoded':syntargetfile}, mapper, variantsites_min_acc=0.99)
```

```
Aligning to 2285 nts of fluPB2
Aligning to 2297 nts of fluPB1
Aligning to 2183 nts of fluPA
Aligning to 1725 nts of fluHA
Aligning to 1516 nts of fluNP
Aligning to 1357 nts of fluNA
Aligning to 973 nts of fluM1
Aligning to 285 nts of fluM2
Aligning to 839 nts of fluNS1
Aligning to 367 nts of fluNS2

Aligning with minimap2 version 2.12-r827
```

##### Amount to trim termini¶

In looking for the linear amplicons described above, we allow a bit of trimming of the 5' termini (defined by *primer5\_mix*) and the 3' termini (defined by *primer3*) since sometimes the CCSs don't seem to begin exactly on the expected nucleotide, perhaps due to additions during end repair or slopping adaptor trimming by CCS.

Therefore, we trim a few nucleotides from our required termini ends:

In [12]:

```
trim_termini = 5
```

##### Un-circularize CCSs on circularized template¶

For the polymerase genes, a major problem was that a disproportionate number of the reads aligned to short segments with internal deletions rather than full-length genes.
The reason is that many cycles of PCR were done in the library prep, and these preferentially enriched the shorter segments.

Alistair therefore devised a strategy to specifically enrich for full-length polymerase reads in one sample.
This strategy involved circularizing the amplicons from the 10X, and then re-linearizing them with primers that annealed near the center of the polymerase genes.
This strategy only create linear product from amplicons with polymerase genes that include a region near the center the gene.

However, for the downstream analysis we need to "un-circularize" these CCSs by identifying the ends of the original amplicon from the 10X material and re-slicing the CCSs to make those the termini.
Here we do this.

First, get the samples that were circularized in the way described above:

In [13]:

```
circ_samples = [ccs.samplename for ccs in ccslist if 'circ' in ccs.samplename]
print("Un-circularizing the following samples:\n\t" + 
      '\n\t'.join([sample for sample in circ_samples]))
```

```
Un-circularizing the following samples:
	2018-08-08_Pol_circ
```

Now define the sequences of the primers used for the linearization of the circularized products, and make sure that these primers anneal uniquely and end-to-end in the expected flu genes:

In [14]:

```
circ_primers = {
    'fluPB2':{
        'for':'CGAAGCAATAATTGTGGCCATG',
        'rev':dms_tools2.utils.reverseComplement('GCAATCGACTGTTCGTCTCTC')
        },
    'fluPB1':{
        'for':'CCTGGAATGATGATGGGCATG',
        'rev':dms_tools2.utils.reverseComplement('GCTCAATGATGCAGTCCCATC')
        },
    'fluPA':{
        'for':'TGAAGCAATATGATAGTGATGAACCAG',
        'rev':dms_tools2.utils.reverseComplement('AATCGCCTACATCTTTACAATCGTC')
        }
    }

for flugene, primers in circ_primers.items():
    geneseq = mapper.targetseqs[flugene]
    assert all(geneseq.count(primer) == 1 for primer in primers.values())
    ifor = geneseq.index(primers['for']) 
    irev = geneseq.index(primers['rev'])
    print(f"For {flugene} (length {len(geneseq)}), rev and for circ primer span nts "
          f"{irev + 1} to {irev + len(primers['rev'])} and "
          f"{ifor + 1} to {ifor + len(primers['for'])}.")
    assert ifor == irev + len(primers['rev']), "primers not end-to-end"
```

```
For fluPB2 (length 2285), rev and for circ primer span nts 1156 to 1176 and 1177 to 1198.
For fluPB1 (length 2297), rev and for circ primer span nts 1193 to 1213 and 1214 to 1234.
For fluPA (length 2183), rev and for circ primer span nts 1140 to 1164 and 1165 to 1191.
```

Now we create a regular expression match string that gives the sequence expected for the CCSs from the circularized amplicons.
These should have termini defined by the linearization primers above, and internally should have the sequences that will be the termini after our un-circularization.
We create a match string that can match any of the genes that were circularized in Alistair's library prep:

In [15]:

```
circ_match_str = []
for flugene, primers in circ_primers.items():
    # 5' and 3' termini of gene when un-circularized
    gene_termini5 = primer5_mix[flugene][trim_termini :] 
    gene_termini3 = dms_tools2.utils.reverseComplement(primer3)[ : -trim_termini] 
    
    # match str for gene
    circ_match_str.append(
            f'(?P<circ_for>{primers["for"]})' + # forward circularization primer
            'N+' + # chunk of un-circularized amplicon
            f'(?P<gene_termini3>{gene_termini3})' +
            'N+' + # a few nucleotides between circularized termini
            f'(?P<gene_termini5>{gene_termini5})' +
            'N+' + # chunk of un-circularized amplicon
            f'(?P<circ_rev>{primers["rev"]})' # reverse circularization primer
            )
    
# call whole match *oriented_CCS* to get CCS and qvals in match orientation
circ_match_str = f"^(?P<oriented_CCS>{'|'.join(circ_match_str)})$"
```

Now we match all sequences with the expected patterns using dms\_tools2.pacbio.matchSeqs, indicating the matched ones as *circularized* in the resulting data frame:

In [16]:

```
circ_df = dms_tools2.pacbio.matchSeqs(
        pandas.concat(ccs.df for ccs in ccslist if ccs.samplename in circ_samples),
        circ_match_str,
        'CCS',
        'circularized'
        )
```

Plot how many of the CCSs had the sequence as expected if they were from the circularized products.
As shown below, a large majority can be identified as the expected circularized products:

In [17]:

```
circ_plot = os.path.join(aligndir, 'circ_match_plot.pdf')

(ggplot(circ_df, aes('samplename', fill='circularized')) +
    geom_bar(position='dodge') +
    scale_fill_manual(COLOR_BLIND_PALETTE_GRAY[1 : ])
    ).save(circ_plot, width=1.5 * (1 + len(circ_samples)), height=3)

showPDF(circ_plot, width=400)
```

Now we un-circularize all of the identified circularized CCSs.
To do this, we slice from start of the gene's 5'-termini to the end of the CCS, and then concatenate from the start of the CCS to the end of the genes' 3'-termini.
Note that we do this on the CCSs that have already been oriented above, and also un-circularize the Q-values.
Finally, we add the un-circularized CCSs back to the original list of CCSs:

In [18]:

```
# un-circularize in `circ_df`
circ_df = (
    circ_df
    # only un-circularize circularized CCSs
    .query('circularized')
    # get start of gene's 5 termini, and end of 3' termini
    .assign(istart=lambda x: x.apply(lambda row: 
                row.oriented_CCS.index(row.gene_termini5), axis=1),
            iend=lambda x: x.apply(lambda row:
                row.oriented_CCS.index(row.gene_termini3) + len(row.gene_termini3),
                axis=1))
    .assign(CCS=lambda x: x.apply(lambda row:
                row.oriented_CCS[row.istart : ] + row.oriented_CCS[ : row.iend],
                axis=1),
            CCS_qvals=lambda x: x.apply(lambda row: numpy.concatenate(
                [row.oriented_CCS_qvals[row.istart : ], 
                row.oriented_CCS_qvals[ : row.iend]]), axis=1))
    # keep only columns present in all entries in all CCS dfs
    [ccslist[0].df.columns] 
    )

for ccs in ccslist:
    if ccs.samplename in circ_samples:
        ccs.df = circ_df.query('samplename == @ccs.samplename')
```

##### Match and align CCSs¶

We use the function dms\_tools2.pacbio.matchAndAlignCCS to go through the CCS's, find the ones that are "barcoded" (have the proper elements), and the align these barcoded genes.
All the results go into a pandas data frame called *df\_ccs*.

Specifically, we look for CCS's that have:

- Any 5' primer from Alistair's mix, trimmed a bit at the start since the first few nucleotides in the CCS might be off. We call this *termini5*.
- The mRNA itself, which we call the *gene*. Since we define the polyA tail as beginning on the first `A` at the polyA signal, the gene must end on a non-`A` nucleotide (which is the IUPAC code `B`).
- The polyA tail, which is effectively a spacer between the *gene* and the *UMI* / *barcode*. This tail is expected to be 30 `A`'s from 10X primer, but we allow it to be anything greater than 25 to account for the sloppiness in sequencing and primer synthesis associated with runs. We also use regex fuzzy matching to allow the polyA to have up to two non-`A` nucleotides internal to it, since for instance the `VN` tooth in the oligo-dT primer might sometimes anneal in the wrong spot. We call this the *spacer* in the call to dms\_tools2.pacbio.matchAndAlignCCS, but then rename to *polyA*.
- The 10-nucleotide UMI from the 10X primer, which we call *UMI*.
- The 16-nucleotide cell barcode from the 10X primer, which we call *barcode*.
- The 3' primer that Alistair used, which anneals in the read 1 primer binding site on the 10X primer. We call this *termini3*.

Note also that when initialized the dms\_tools2.minimap2.Mapper used in the call to dms\_tools2.pacbio.matchAndAlignCCS, we specified that *M1* and *M2* are isoforms, and *NS1* and *NS2* are isoforms.

We use a dms\_tools2.minimap2.MutationCaller to call mutations.
The mutation calling uses modest clipping of the target and soft clipping of the query to avoid calling mutations right at the ends.
We use a dms\_tools2.seqnumbering.TranscriptConverter to convert the numbers of the mutations into 1-based numbering of the full viral RNA (in mRNA orientation) and call amino-acid substitutions, using the mRNA / CDS annotations in ./data/flu\_sequences/flu-wsn.gb.

In [19]:

```
transcriptconverter = dms_tools2.seqnumbering.TranscriptConverter(
        'data/flu_sequences/flu-wsn.gb', ignore_other_features=True)

mutationcaller = dms_tools2.minimap2.MutationCaller(mapper,
        transcriptconverter=transcriptconverter,
        query_softclip=10, target_clip=20)

df_ccs = dms_tools2.pacbio.matchAndAlignCCS(
        ccslist=ccslist,
        mapper=mapper,
        termini5='|'.join([s[trim_termini : ] for s in primer5_mix.values()]),
        gene='N+B',
        spacer='AAA(A{19,}){e<=2}AAA', # regex fuzzy matching allows 2 mismatch
        umi='N{10}',
        barcode='N{16}',
        termini3=dms_tools2.utils.reverseComplement(primer3)[ : -trim_termini],
        targetvariants=targetvariants,
        mutationcaller=mutationcaller
        ).rename(columns={'has_spacer':'has_polyA'})

print("Attempted to match and align all {0} CCSs".format(len(df_ccs)))
```

```
Attempted to match and align all 953652 CCSs
```

##### Stats on matching / alignment¶

Below we analyze the statistics on the matching and aligning of the CCSs:

In [20]:

```
# possible matching / alignment categories
match_align_cats = ['total', 'has_termini3', 'has_polyA', 'has_termini5',
                    'barcoded', 'gene_aligned', 'CCS_aligned']

# tabulate statistics on matching / alignment
match_align_df = (
    df_ccs
    .assign(total=True)
    .melt(id_vars=['samplename'], 
          value_vars=match_align_cats,
          var_name='category',
          value_name='number of CCSs')
    .groupby(['samplename', 'category'], as_index=False)
    .aggregate('sum')
    .assign(category=lambda x: pandas.Categorical(
            x.category.str.replace('_', ' '), 
            [col.replace('_', ' ') for col in match_align_cats]))
    .sort_values(['samplename', 'category'])
    .assign(percent=lambda x: 100 * x['number of CCSs'] / 
               x.groupby('samplename')['number of CCSs'].transform('max'))
    )

# plot the matching alignment statistics
match_align_plot = os.path.join(aligndir, 'match_align_plot.pdf')
(ggplot(match_align_df, aes('samplename', 'number of CCSs')) +
    geom_bar(aes(fill='category'), position='dodge', stat='identity') +
    scale_fill_manual(COLOR_BLIND_PALETTE_GRAY) +
    xlab('sequencing run') 
    ).save(match_align_plot,
           width=1.7 * (1 + len(seqruns)),
           height=3)
showPDF(match_align_plot)
```

We see that across all samples, the vast majority of CCSs have the 3' termini and the polyA, as expected given that those need to be there for the 3' primer to work.

However, only a fraction have the 5' termini.
This could be because only a small fraction of all sequences in the initial PCR template pool will be flu mRNAs with the right termini for the 5' primer, so there may be spurious amplification (by the 3' primer or just linear amplification product) that doesn't have the 5' termini.

Most of the CCSs that have the 5' termini are also *barcoded* in the sense that they fully match the expected patterns that allow us to call a barcode and UMI.
The frequency with which CCSs have the 5' termini is lower for the samples that only contain polymerase genes, as expected if these are amplified less efficiently.

Of these *barcoded* CCSs, most of them have mRNA inserts ("genes") that align to the flu targets, and so are in the *gene aligned* category.

We also see that if we align the full CCS without requiring it to match the termini / barcode / polyA, we get a few more than for the *gene aligned* category, indicating that some of the CCSs for which we can't call barcodes still have flu in them.
These may have mutations in the termini / polyA, or they may be some sort of chimera.
In any case, since the *CCS aligned* category is only modestly larger than the *gene aligned* category, for all subsequent analyses we'll focus just on the *gene aligned* CCSs.

The exception to all of this is the run where we circularized the material and then un-circularized it.
For this run, most CCSs align--part of this is because in the un-circularization process above, we already got rid of sequences that don't have the right termini.

##### Get CCSs with gene aligned¶

For the reasons explained immediately above, all remaining analyses will focus only on the CCSs for which the gene aligned.
So only keep these in our *df\_ccs* data frame.

In [21]:

```
df_ccs = df_ccs.query('gene_aligned').reset_index(drop=True)
```

#### Quality-control alignments¶

For the CCSs for which the gene insert aligned to flu, we now do some quality controlling on several important aspects.

We do this by adding a column to our data frame named `QC_filtered`.
Initially all entries in this columns are the string "passes\_filters".
As we go through each filter in order below, we add a string describing the filtering reason for each CCS that fails that filter.
Once a CCS fails one filter (in the order they are provided below), we don't continue checking it against the other filters.

In [22]:

```
df_ccs['pass_QC'] = True
df_ccs['QC_fail_reason'] = 'passes_QC'
```

##### Barcode accuracy¶

We want the barcodes called in the CCSs to be high accuracy.
This might not be the case if molecules with different barcodes anneal during the PCR, such that a different barcode is sequenced on each strand of the SMRTbell that forms the CCS.
So we tabulate some statistics about the distribution of barcode accuracies among CCSs that pass our filters so far:

In [23]:

```
(df_ccs
    .query('pass_QC')
    [['barcode_accuracy']]
    .describe(percentiles=[0.001, 0.01])
    )
```

Out[23]:

|  | barcode\_accuracy |
| --- | --- |
| count | 239914.000000 |
| mean | 0.999865 |
| std | 0.001624 |
| min | 0.910318 |
| 0.1% | 0.974971 |
| 1% | 0.997512 |
| 50% | 1.000000 |
| max | 1.000000 |

Above we see that the barcode accuracies are consistently very high, with over 99% having accuracies that exceed the accuracy threshold of 0.999 used when building the CCSs with PacBio's ccs program, and virtually all of them having accuracies of >98%.
Just to be safe, we will filter for CCSs that have a barcode accuracy $\ge$0.999, which will eliminate only a very small fraction:

In [24]:

```
fail_index = df_ccs.query('pass_QC & (barcode_accuracy < 0.999)').index
df_ccs.loc[fail_index, 'pass_QC'] = False
df_ccs.loc[fail_index, 'QC_fail_reason'] = 'low barcode accuracy'
```

##### Additional alignments¶

We want to get rid of CCSs that align to multiple different influenza genes.
The reason is that these probably represent some sort of PCR chimera / fusion during the library preparation.
Note that we are only looking at multiple alignments to different targets / isoforms (we specified that M1 / M2 and NS1 / NS2 are isoforms in the `target_isoforms` argument to the dms\_tools2.minimap2.Mapper), so this will not purge alignments that involve the same gene (these could be complex deletions) or different splice forms.
First, we tabulate some statistics about the distribution of the number of additional alignments to different targets among CCSs that pass our filters so far:

In [25]:

```
(df_ccs
    .query('pass_QC')
    [['gene_aligned_n_additional_difftarget']]
    .describe(percentiles=[0.995, 0.999, 0.9999])
    )
```

Out[25]:

|  | gene\_aligned\_n\_additional\_difftarget |
| --- | --- |
| count | 236706.000000 |
| mean | 0.001496 |
| std | 0.048268 |
| min | 0.000000 |
| 50% | 0.000000 |
| 99.5% | 0.000000 |
| 99.9% | 1.000000 |
| 99.99% | 2.000000 |
| max | 2.000000 |

There are very few CCSs with additional alignments to other targets (less than 1%), and we remove them:

In [26]:

```
fail_index = df_ccs.query('pass_QC & gene_aligned_n_additional_difftarget').index
df_ccs.loc[fail_index, 'pass_QC'] = False
df_ccs.loc[fail_index, 'QC_fail_reason'] = 'aligns to multiple targets'
```

##### Excessive alignment trimming¶

For perfect matches the of query CCSs to the target flu mRNAs, alignment of the query to the target is end-to-end, with no trimming of either the query or the target at either end.

However, in practice, in many cases there is trimming of the query or the target.
Here we QC filter based on the presence of excessive trimming.

###### Trimming of query start¶

Based on the way that the products were PCR-amplified, we expect the query starts to align exactly with the target start, so there shouldn't be any need for trimming.
Below are tabulated statistics about the distribution of trimming at the query start:

In [27]:

```
(df_ccs
    .query('pass_QC')
    [['gene_aligned_n_trimmed_query_start']]
    .describe(percentiles=[0.99, 0.999, 0.9999])
    )
```

Out[27]:

|  | gene\_aligned\_n\_trimmed\_query\_start |
| --- | --- |
| count | 236550.000000 |
| mean | 0.170146 |
| std | 13.132135 |
| min | 0.000000 |
| 50% | 0.000000 |
| 99% | 0.000000 |
| 99.9% | 3.000000 |
| 99.99% | 730.177000 |
| max | 2464.000000 |

Most (>99%) queries align exactly with the target at their start, with no trimming.
There are a few nucleotides trimmed from a modest fraction (<1%).
This modest trimming is explainable---the 5' primers are similar for the different flu genes, so there may occassionally be mis-priming so that the primer amplifies a different flu mRNA, which might lead to a bit of trimming at the very start of the alignment.
There are then a very small number (less than <0.01%) that have very large amounts trimmed.
We will discard CCSs with more than 5 nucleotides trimmed from the query start:

In [28]:

```
fail_index = df_ccs.query('pass_QC & (gene_aligned_n_trimmed_query_start > 5)').index
df_ccs.loc[fail_index, 'pass_QC'] = False
df_ccs.loc[fail_index, 'QC_fail_reason'] = 'excessive trimming of query start'
```

###### Trimming of target start¶

We also expect all of the alignments to start exactly at the beginning of the target, because the PCR-amplification should lead to this.
Below are tabulated statistics about the distribution of trimming at the target start:

In [29]:

```
(df_ccs
    .query('pass_QC')
    [['gene_aligned_n_trimmed_target_start']]
    .describe(percentiles=[0.99, 0.999, 0.9999])
    )
```

Out[29]:

|  | gene\_aligned\_n\_trimmed\_target\_start |
| --- | --- |
| count | 236433.000000 |
| mean | 0.107866 |
| std | 10.119755 |
| min | 0.000000 |
| 50% | 0.000000 |
| 99% | 0.000000 |
| 99.9% | 4.000000 |
| 99.99% | 110.976000 |
| max | 1884.000000 |

This distribution looks very similar to that for trimming of the query starts immediately above, and (for the same logic explained there), we will discard CCSs with more than 5 nucleotides trimmed from the target start:

In [30]:

```
fail_index = df_ccs.query('pass_QC & (gene_aligned_n_trimmed_target_start > 5)').index
df_ccs.loc[fail_index, 'pass_QC'] = False
df_ccs.loc[fail_index, 'QC_fail_reason'] = 'excessive trimming of target start'
```

###### Trimming query ends¶

We expect the queries to end at the end of the target, and so there to be no clipping of the query ends.
There are three plausible reasons for clipping of the query ends:

1. A PCR artifact that leads to fusion of a flu gene with a cellular gene. We tried to filter out fusions of two different flu genes above by removing CCSs with additional alignments to a different flu target, but that filter won't remove fusions to things that aren't flu genes--although these are expected to be much more abundant. We would like to filter CCSs for which this occurs.
2. The transcription of the mRNA from the query continuing past the polyA tail, or the polyA tail not being fully trimmed and removed. We might want to retain these. However, we expect the extension that has to be trimmed in this case to be small, probably less than 50 nt.
3. There is an internal deletion and for whatever reason the mapping failed to handle it properly and just trimmed the region after the deletion. We want to keep these. We would expect them to align with "additional alignments" since the trimmed region should be in a different alignment.

Below we tabulate some statistics on the distribution of trimming, for CCSs both with and without additional alignments (these distributions are hard to plot since they are so skewed):

In [31]:

```
(df_ccs
    .query('pass_QC')
    .assign(has_additional_alignment=lambda x: x.gene_aligned_n_additional > 0)
    .groupby(['has_additional_alignment'])
    .gene_aligned_n_trimmed_query_end
    .describe(percentiles=[0.9, 0.95, 0.98, 0.99, 0.999])
    )
```

Out[31]:

|  | count | mean | std | min | 50% | 90% | 95% | 98% | 99% | 99.9% | max |
| --- | --- | --- | --- | --- | --- | --- | --- | --- | --- | --- | --- |
| has\_additional\_alignment |  |  |  |  |  |  |  |  |  |  |  |
| False | 200208.0 | 6.947869 | 32.853912 | 0.0 | 1.0 | 16.0 | 39.0 | 65.0 | 83.0 | 473.793 | 2201.0 |
| True | 36195.0 | 8.989115 | 51.169202 | 0.0 | 1.0 | 21.0 | 44.0 | 74.0 | 101.0 | 943.000 | 2434.0 |

We see that the distributions don't look much different between CCSs with and without additional alignments, so the internal deletion explanation doesn't appear to explain most of the clipping.
We will discard CCSs with more than 50 nt of trimming at the query end as almost certainly spurious, but keep those with less than 50 nt in case there is transcription past the polyA and/or problems calling the polyA.
Looking at the table above, we can see that this means we are discarding about 5% of the CCSs:

In [32]:

```
fail_index = df_ccs.query('pass_QC & (gene_aligned_n_trimmed_query_end > 50)').index
df_ccs.loc[fail_index, 'pass_QC'] = False
df_ccs.loc[fail_index, 'QC_fail_reason'] = 'excessive trimming of query end'
```

###### Trimming target ends¶

If transcription proceeds all the way to the target end, there should also not be trimming of the target ends.
Here are some actual tabulated statistics:

In [33]:

```
(df_ccs
    .query('pass_QC')
    [['gene_aligned_n_trimmed_target_end']]
    .describe(percentiles=[0.75, 0.85, 0.9, 0.95, 0.99, 0.999])
    )
```

Out[33]:

|  | gene\_aligned\_n\_trimmed\_target\_end |
| --- | --- |
| count | 228300.000000 |
| mean | 90.344779 |
| std | 345.567945 |
| min | 0.000000 |
| 50% | 0.000000 |
| 75% | 2.000000 |
| 85% | 4.000000 |
| 90% | 6.000000 |
| 95% | 862.000000 |
| 99% | 1771.000000 |
| 99.9% | 1783.701000 |
| max | 1962.000000 |

It's important to make sure that the target trimming doesn't tend to occur on the same sequences with query trimming, as this might indicate that the end of the query just failed to align because it's too short of a final chunk after a deletion.
The plot below shows this is **not** the case, which alleviates the concern that the query and target end trimming involves deletions where the 3' end is just too short to align:

In [34]:

```
(df_ccs
    .assign(substantial_query_trimming=lambda x: x.gene_aligned_n_trimmed_query_end > 10)
    .groupby('substantial_query_trimming')
    .gene_aligned_n_trimmed_target_end
    .describe(percentiles=[0.75, 0.85, 0.9, 0.95, 0.99, 0.999])
    )
```

Out[34]:

|  | count | mean | std | min | 50% | 75% | 85% | 90% | 95% | 99% | 99.9% | max |
| --- | --- | --- | --- | --- | --- | --- | --- | --- | --- | --- | --- | --- |
| substantial\_query\_trimming |  |  |  |  |  |  |  |  |  |  |  |  |
| False | 208247.0 | 99.094335 | 360.677432 | 0.0 | 1.0 | 2.0 | 4.0 | 8.0 | 1003.0 | 1771.00 | 1787.0 | 1962.0 |
| True | 31667.0 | 15.256260 | 142.198420 | 0.0 | 0.0 | 0.0 | 0.0 | 0.0 | 1.0 | 857.68 | 1762.0 | 2165.0 |

So overall, the table two above shows that there is substantial trimming of the target ends in 5-15% of CCSs.

There are plausible legitimate and artifactual explanations:

- *legitimate*: premature poly-adenylation of the flu transcripts leading to truncation of the query, or some form of internal deletion in the vRNA moving the polyA signal up in the transcript.
- *artifactual*: the polyA primer anneals in an A rich stretch internal to the gene.

We can't really distinguish between these without prior work, but some loose analyses not shown here favor the artifactual explanation.
Also, it is thought that at least at the vRNA level, both termini are needed for viral transcription, so if premature polyadenylation is occuring, then it is *probably* not a genetic feature of the viruses but rather a transcriptional feature of some mRNAs--and our current downstream analysis is focused more on genetic features of the virus.

Therefore, we will filtering any alignments with more than 10 nucleotides of trimming for now (but this probably deserves further study at some point):

In [35]:

```
fail_index = df_ccs.query('pass_QC & (gene_aligned_n_trimmed_target_end > 10)').index
df_ccs.loc[fail_index, 'pass_QC'] = False
df_ccs.loc[fail_index, 'QC_fail_reason'] = 'excessive trimming of target end'
```

##### Get QC-ed alignments¶

Now we tabulate the number of alignments that failed QC for each reason:

In [36]:

```
(df_ccs
    .assign(number=1)
    .groupby(['pass_QC', 'QC_fail_reason'])
    [['number']]
    .agg('count')
    )
```

Out[36]:

|  |  | number |
| --- | --- | --- |
| pass\_QC | QC\_fail\_reason |  |
| False | aligns to multiple targets | 156 |
|  | excessive trimming of query end | 8103 |
|  | excessive trimming of query start | 117 |
|  | excessive trimming of target end | 18379 |
|  | excessive trimming of target start | 30 |
|  | low barcode accuracy | 3208 |
| True | passes\_QC | 209921 |

As the table above shows, less than 25% of the alignments fail the filters, and most of the failures are due to excessive trimming of the query end or target end.

We keep just the QC-ed aligned CCSs for subsequent work:

In [37]:

```
df_ccs = df_ccs.query('pass_QC')
```

#### Summary of QC-ed alignments¶

We are now going to analyze how many QC-ed alignments we have for each gene, and the length distribution of these alignments.

First, make the aligned genes a categorical variable so gene names are displayed in the desired order:

In [38]:

```
targetnames = list(mapper.targetseqs.keys())
df_ccs['gene_aligned_target'] = pandas.Categorical(
        df_ccs.gene_aligned_target, targetnames)
```

##### Number of alignments per gene¶

Now we examine the number of QC-ed aligned CCSs for each gene.
We do this both aggregating over all sequencing runs and looking at the runs individually:

In [39]:

```
# base plot of number of sequences per gene
plot_nseqs = (
    ggplot(df_ccs, aes('gene_aligned_target')) +
        geom_bar() +
        theme(axis_text_x=element_text(angle=90, hjust=0.5)) +
        xlab("flu gene") +
        ylab("QC-ed aligned sequences")
    )

# plot total for all runs and by run
ncol = 3
for plottype, facet, n in [
        ('all_runs', geom_blank(), 1),
        ('by_run', facet_wrap('~samplename', ncol=3), len(seqruns))]:
    print("\nNumber of alignments, {0}:".format(plottype.replace('_', ' ')))
    nseqs_plot = os.path.join(aligndir, 'nsequences_{0}.pdf'.format(plottype))
    (plot_nseqs + facet).save(nseqs_plot, height=1 + 1.8 * math.ceil(n / ncol), 
                              width=0.8 + 2 * min(n, ncol))
    showPDF(nseqs_plot, width=80 + 200 * min(n, ncol))
```

```
Number of alignments, all runs:
```

```
Number of alignments, by run:
```

The plots above show the different distributions of genes per run, as expected as some runs tried to heavily load polymerase genes while others tried to load the other genes.
This is because the polymerase genes were rare when everything was loaded equally (2017-06-08 run), so all the other ones were attempting to balance things out.
But see the caveat about polymerase deletions that becomes apparent below...

Also, we see that there are many fewer alignments for M2 than M1, and for NS2 than NS1.
This is as expected based on the known ratios of these splice forms.

##### Distribution of alignment lengths¶

Not all of the alignments necessarily cover the entire gene, as there are often deletions in flu segments.
So we also plot the distribution of alignments by length for each segment, where we define the *alignment length* as the total number of aligned nucleotides, which can be either identities or mismatches (but not indels).
We plot this both across all sequencing runs, and for each run individually:

In [40]:

```
# add columnn with alignment lengths
df_ccs['alignment_length'] = df_ccs.gene_aligned_alignment.apply(
        lambda a: dms_tools2.minimap2.numAligned(a.cigar_str))

# base plot of alignment lengths
plot_align_len = (
    ggplot(df_ccs, aes('alignment_length')) +
        geom_histogram(aes(x='alignment_length'), bins=15) +
        theme(axis_text_x=element_text(angle=90, hjust=0.5)) +
        ylab("relative number of alignments") +
        xlab("alignment length")
        )

# plot for all runs and by run
for plottype, facet, n in [
        ('all_runs', facet_wrap('~ gene_aligned_target', nrow=1), 1),
        ('by_run', facet_grid('samplename ~ gene_aligned_target'), len(seqruns))]:
    print("\nAlignment lengths, {0}:".format(plottype.replace('_', ' ')))
    align_len_plot = os.path.join(aligndir, 'alignment_len_{0}.pdf'.format(plottype))
    (plot_align_len + facet).save(align_len_plot, width=15, height=2 * n)
    showPDF(align_len_plot)
```

```
Alignment lengths, all runs:
```

```
Alignment lengths, by run:
```

As can be seen above, for most genes the alignments are strongly peaked at the length expected for the full-length gene.
But for the polymerase genes (PB2, PB1, and PA), for most of the runs most of the reads are actually deletions that are much smaller than the full-length size.
The enrichment of deletions for the polymerase probably exceeds their actual biological frequency due to amplification bias that prefers shorter segments during PCR.
However, the circularization strategy did an excellent job of enriching for longer full-length polymerase genes.

For HA, there are also noticeable peaks at smaller sizes.

##### Near full-length alignments¶

Given the plots above that show that for some genes, a substantial number of alignments are for partially deleted genes, we also plot the number "near full-length" alignments.
We define an alignment as "full-length" if the alignment length (as defined immediately above) is no more than 20 nucleotides less than the expected length of the alignment target.

In [41]:

```
# assign target lengths and near full-length status
target_lengths = {name:len(seq) for name, seq in mapper.targetseqs.items()}
df_ccs = (
    df_ccs
    .assign(target_length=lambda x: x.gene_aligned_target.map(target_lengths))
    .assign(full_length=lambda x: (x.target_length <= 20 + x.alignment_length))
    )

plot_nseqs_full = (
    ggplot(df_ccs, aes('gene_aligned_target')) +
        geom_bar(aes(fill='full_length')) +
        theme(axis_text_x=element_text(angle=90, hjust=0.5)) +
        xlab("flu gene") +
        ylab("QC-ed aligned sequences") +
        scale_fill_manual(COLOR_BLIND_PALETTE_GRAY[1 : ],
                          name='full length')
    )

# plot total for all runs and by run
ncol = 3
for plottype, facet, n in [
        ('all_runs', geom_blank(), 1),
        ('by_run', facet_wrap('~samplename', ncol=ncol), len(seqruns))]:
    print("\nNumber of alignments, {0}:".format(plottype.replace('_', ' ')))
    nseqs_plot = os.path.join(aligndir, 'nseqs_full_len_{0}.pdf'.format(plottype))
    (plot_nseqs_full + facet).save(nseqs_plot,
            height=0.5 + 2 * math.ceil(n / ncol), width=2 * (min(n, ncol) + 0.8))
    showPDF(nseqs_plot, width=220 * (min(n, ncol) + 0.8))
```

```
Number of alignments, all runs:
```

```
Number of alignments, by run:
```

We see basically what is expected from the sets of plots in the previous two subsections.
For all runs except for the one that use the circularization strategy, the polymerase reads are highly biased towards short fragments.
Note that we have relatively little for M2 / NS2, but these sequences are contained within the M1 / NS1, so it's probably OK to have less for these.

##### Passes/accuracy for alignments¶

Back at the initial running of `ccs`, we set thresholds for how many passes of sequencing and the minimum accuracy required for our CCSs.
Are these limiting?
For instance, when we look at the full-length alignments of the long polymerase genes, are we losing lots of sequences because of overly stringent thresholds?
We can address this by looking at the actual distribution of number of passes and CCS accuracy for our QC-ed aligned genes, both for full-length and non-full-length (e.g., with deletions) viral genes.
Below we plot these, using dashed green lines to show the minimum cutoffs:

In [42]:

```
ncol = 3

npasses_plot = os.path.join(aligndir, 'npasses.pdf')
(ggplot(df_ccs, aes('gene_aligned_target', 'passes', fill='full_length')) +
    geom_boxplot(outlier_size=0.2) +
    scale_y_log10(limits=(minpasses, df_ccs.passes.max())) +
    scale_fill_manual(COLOR_BLIND_PALETTE_GRAY[1 : ]) +
    facet_wrap('~ samplename') +
    theme(axis_text_x=element_text(angle=90, hjust=0.5)) +
    geom_hline(yintercept=minpasses, linetype='dashed',
        color=COLOR_BLIND_PALETTE_GRAY[3])
    ).save(npasses_plot, height=0.5 + 2 * math.ceil(len(seqruns) / ncol),
            width=1.8 * (1 + min(ncol, len(seqruns))))
showPDF(npasses_plot)

error_rate_plot = os.path.join(aligndir, 'error_rate.pdf')
(ggplot(df_ccs, aes('gene_aligned_target', 'CCS_accuracy', fill='full_length')) +
    geom_boxplot(outlier_size=0.2) +
    scale_y_continuous(limits=(ccsminaccuracy, 1)) +
    scale_fill_manual(COLOR_BLIND_PALETTE_GRAY[1 : ]) +
    facet_wrap('~ samplename') +
    theme(axis_text_x=element_text(angle=90, hjust=0.5)) +
    geom_hline(yintercept=ccsminaccuracy, linetype='dashed',
        color=COLOR_BLIND_PALETTE_GRAY[3])
    ).save(error_rate_plot, height=0.5 + 2 * math.ceil(len(seqruns) / ncol),
            width=1.8 * (1 + min(ncol, len(seqruns))))
showPDF(error_rate_plot)
```

As is apparent from these plots, the vast majority of sequences are well above both minimum thresholds.
Although the full-length polymerase genes do tend to have less passes and lower accuracy, most of them are still well above the threshold.
This suggests that we are **not** losing lots of fairly good sequences to overly stringent thresholds.

#### Analyze viral barcodes¶

Each viral gene is either fully wildtype or "doubly synonymously barcoded" by a pair of synonymous mutations near each termini.
Note that these viral barcodes are distinct from the cell barcodes added by the 10X system.
These viral barcodes were called during the alignment using the dms\_tools2.minimap2.TargetVariants initialized above.

Here we examine these barcodes.
The most important reason to do this is to test for strand exchange during PCR, or pairing of different variants into the same dsDNA molecule during library preparation.
These would manifest as CCSs in which the different ends are assigned to different viral barcode variants, or in which the viral barcode sites are low accuracy.

##### Inspect viral barcodes by gene¶

First we just plot the viral barcodes that are called for each gene in each sequencing run.
This plot is below.
In addition to faceting on sequencing run, we also facet on all QC-ed aligned sequences, and just those that correspond to full-length viral genes.

In [43]:

```
# create a directory for the CCS analysis
analysisdir = os.path.join(pacbioresultsdir, 'variant_and_mutation_analysis')
os.makedirs(analysisdir, exist_ok=True)

# order viral barcode categories from most to least for plotting
viralbarcode_cats = (
    df_ccs
    .groupby(['gene_aligned_target_variant'])
    .CCS
    .agg('count')
    .sort_values(ascending=False)
    .index
    )
df_ccs['gene_aligned_target_variant'] = pandas.Categorical(
        df_ccs['gene_aligned_target_variant'], viralbarcode_cats)

viralbarcode_plot = os.path.join(analysisdir, 'viralbarcodeplot.pdf')
(ggplot(pandas.concat([
            df_ccs.assign(full_length='any length'),
            df_ccs.query('full_length').assign(full_length='full length')]),
        aes('gene_aligned_target')) +
    geom_bar(aes(fill='gene_aligned_target_variant')) +
    facet_grid('samplename ~ full_length', scales='free_y') +
    scale_fill_manual(COLOR_BLIND_PALETTE_GRAY, name='viral barcode') +
    theme(axis_text_x=element_text(angle=90, hjust=0.5), panel_spacing_x=0.5) +
    ylab('QC-ed aligned sequences\n') +
    xlab('flu gene')
    ).save(viralbarcode_plot, width=8, height=1.9 * (len(seqruns) + 0.5))
showPDF(viralbarcode_plot, width=700)
```

These plots look pretty good.
We want to avoid are *mixed* barcodes (indicating chimeras) or *low accuracy* barcodes (indicating possible chimeric dsDNA during library preparation).
These states are very rare for all genes on all sequencing runs.
With the exception of PB2 / PB1 / PA on a few runs, things look very consistent across genes and sequencing runs, with most barcodes either assigned to fully *wildtype* or fully *synonymously barcoded*.
The *synonymously barcoded* variant is clearly more common.

Things are a bit more complicated for the polymerase genes, where we can only partially call many of the barcodes on some runs.
This is because there are lots of internal deletions, some of which remove barcode variant sites.
In addition, the PCR enrichment selectively favored these deletions, so there is skewing in abundance.
But if we limit to just full-length genes, things look fairly similar for the polymerase as for other genes.

##### Estimate rate of PCR strand exchange¶

We will now use the above data to estimate the rate of PCR strand exchange.
In principle, this rate could (and probably does to some degree) vary among molecules and sequencing runs.
However, the plots in the previous subsection suggest that this variation is fairly small.
So will just make a single "average" estimate over all the data.

We make the estimate just using the counts of the full *wildtype* and *synonymously barcoded* variants, and ignore the partially barcoded ones.
The reason that we ignore the partially barcoded ones is that they often represent calling of the barcodes at only one end of the molecule, which might not identify strand exchange.
We consider variants to be *chimeric* if their barcoded sites are **either** *mixed* or *low accuracy*, since *low accuracy* can indicate mixed variants in the CCS sequencing.

First, we get and display these statistics:

In [44]:

```
chimera_stats = (
    df_ccs
    .rename(columns={'gene_aligned_target_variant':'viral_barcode'})
    .replace({'viral_barcode':{'mixed':'chimeric', 'low accuracy':'chimeric'}})
    .query('viral_barcode in ["chimeric", "wildtype", "synonymously barcoded"]')
    .assign(number=1) # dummy variable to count in `agg`
    .groupby(['viral_barcode'])
    .agg({'number':'count'})
    .assign(fraction=lambda x: x.number / x.number.sum())
    )

display(HTML(chimera_stats.to_html()))
```

|  | number | fraction |
| --- | --- | --- |
| viral\_barcode |  |  |
| chimeric | 4961 | 0.025896 |
| synonymously barcoded | 116191 | 0.606517 |
| wildtype | 70419 | 0.367587 |

Obviously, we don't expect to observe all the chimeric CCSs, because some will be chimeras of a wildtype with wildtype, or synonymously barcoded with synonymously barcoded.
The amplification was done using emulsion PCR, which places each molecule in its own droplet.
We can therefore use the equation for calculating the multiplet frequency described in Bloom (2018, *PeerJ*).
Below we take the function described there, and then calculate the multiplet frequency:

In [45]:

```
def multipletFreq(n1, n2, n12):
    """Estimated multiplet frequency from cell-type mixing experiment.

    `n1`, `n2`, `n12` (`int` or `numpy.ndarray` of integers)
        Number of droplets with at least one cell of type 1,
        at least one cell of type 2, or cells of both types.
    """
    n = numpy.array(n1 * n2 / n12).astype('float')
    mu1 = -numpy.log((n - n1) / n)
    mu2 = -numpy.log((n - n2) / n)
    mu = mu1 + mu2
    return 1 - mu * numpy.exp(-mu) / (1 - numpy.exp(-mu))

chimera_freq = multipletFreq(
        chimera_stats.number.ix['wildtype'],
        chimera_stats.number.ix['synonymously barcoded'],
        chimera_stats.number.ix['chimeric'])

print("The estimated rate at which CCSs are chimeric is {0:.4f}".format(
        chimera_freq))
```

```
The estimated rate at which CCSs are chimeric is 0.0572
```

So we see that about $\approx$5% of the molecules are estimated to be chimeras.
Of these, about half are chimeric between different barcodes, and so can be explicitly excluded as is done in the subsection that immediately follows.
That leaves about 2.5% of CCSs that will still be un-identified chimeras.

##### Figure with number CCSs and strand exchange¶

Now we make a figure that shows the number of aligned post-QC CCSs for each gene and the estimates of strand exchange.

We then copy this to the directory with the paper figures.

In [46]:

```
ccs_per_viralbarcode_fig = os.path.join(
        figsdir, 'ccs_per_viralbarcode.pdf')

(ggplot(pandas.concat([
            df_ccs.assign(full_length='any length'),
            df_ccs.query('full_length').assign(full_length='full length')])
            .assign(flu_gene=lambda x: pandas.Categorical(
                    x.gene_aligned_target.str.replace('flu', ''),
                    [g.replace('flu', '') for g 
                     in mapper.targetseqs.keys()])),
        aes('flu_gene')) +
    geom_bar(aes(fill='gene_aligned_target_variant')) +
    geom_text(aes(label='..count..'), stat='count', size=8,
              angle=90, va='bottom') +
    facet_wrap('~ full_length') +
    scale_y_continuous(expand=(0.05, 0, 0.2, 0)) +
    scale_fill_manual(COLOR_BLIND_PALETTE_GRAY, name='viral barcode') +
    theme(axis_text_x=element_text(angle=90, hjust=0.5)) +
    ylab('number of aligned QC-ed CCSs') +
    xlab('flu gene') +
    ggtitle(f'estimated frequency of PCR chimeras: '
            f'{chimera_freq:.3f}')
    ).save(ccs_per_viralbarcode_fig, width=6, height=3)

showPDF(ccs_per_viralbarcode_fig, width=700)
```

##### Filter chimeras, assign viral barcodes¶

We now remove any of the aligned CCSs with chimeric or low accuracy barcodes.
We also assign each CCS its *viral\_barcode* as *wt\_flu* if it is either fully or partially wildtype barcoded, and *flu\_syn* if it is either fully or partially synonymously barcoded.
This *viral\_barcode* is therefore an indication of which viral variant gave rise to this CCS:

In [47]:

```
nstart = len(df_ccs)

df_ccs = (
    df_ccs.assign(viral_barcode=lambda x: x.gene_aligned_target_variant.str
        .replace(re.compile('(partial ){0,1}wildtype'), 'flu_wt')
        .replace(re.compile('(partial ){0,1}synonymously barcoded'), 'flu_syn'))
    .query('viral_barcode in ["flu_wt", "flu_syn"]')
    )

print("Retained the {0} of {1} ({2:.2f}%) QC-ed aligned CCSs with valid viral barcodes."
      .format(len(df_ccs), nstart, 100 * len(df_ccs) / nstart))
```

```
Retained the 204955 of 209921 (97.63%) QC-ed aligned CCSs with valid viral barcodes.
```

#### 10X cell barcodes / UMIs¶

Now we examine the 10X cell barcodes and UMIs that were called for the aligned genes.
Recall that the cell barcode reports which cell the sequence came from, and the UMI is a unique identifier for molecules.

##### Get valid cell barcodes¶

Only some of the cell barcodes are actually assigned to cells that are called by the 10X `cellranger` pipeline, with the others typically being GEMs with no actual cell (or perhaps sometimes sequencing errors).

We have already called the valid cells with the align\_and\_annotate.ipynb notebook, and written them to `results/cellgenecounts/merged_humanplusflu_cells.tsv`.
First, we read in those barcodes, filtering for the ones from the *IFN\_enriched* sample and removing the suffix `-IFN_enriched` from them.
These represent our set of *valid* cell barcodes:

In [48]:

```
cellbarcodesfile = 'results/cellgenecounts/merged_humanplusflu_cells.tsv'
sample = 'IFN_enriched' # only keep barcodes for this sample

valid_cellbarcodes = set(
    pandas.read_csv(cellbarcodesfile, sep='\t')
    .query('Sample == @sample')
    .CellBarcode.str.replace('-' + sample, '')
    )

print("Read {0} valid cell barcodes for sample {1} from {2}"
        .format(len(valid_cellbarcodes), sample, cellbarcodesfile))
```

```
Read 1626 valid cell barcodes for sample IFN_enriched from results/cellgenecounts/merged_humanplusflu_cells.tsv
```

##### Filter alignments from valid cells¶

We really only care about the alignments that correspond to valid cells, so now we filter for those.
First, we annotate CCS alignments that correspond to a valid cell:

In [49]:

```
df_ccs['valid_cell'] = df_ccs.barcode.isin(valid_cellbarcodes)
```

Next we plot the fraction of aligned CCSs that correspond to valid cells:

In [50]:

```
valid_cells_plots = []
for pos, label in [('stack', 'number'), ('fill', 'fraction')]:
    valid_cells_plot = os.path.join(analysisdir, 'valid_cells_{0}.pdf'.format(label))
    (ggplot(df_ccs, aes('gene_aligned_target')) +
        geom_bar(aes(fill='valid_cell'), position=pos) +
        scale_fill_manual(COLOR_BLIND_PALETTE_GRAY[1 : ], name='valid cell') +
        xlab("flu gene") +
        ylab(label + ' aligned CCSs') +
        theme(axis_text_x=element_text(angle=90, hjust=0.5))
        ).save(valid_cells_plot, width=3, height=2.25)
    valid_cells_plots.append(valid_cells_plot)
showPDF(valid_cells_plots)
```

As can be seen above, a bit over half of the aligned CCSs correspond to valid cells, and there do not appear to be large differences among the flu genes.
The fact that almost half aren't in valid cells isn't too surprising, as there are many empty GEMs that have some RNA but aren't called as cells.

From here on out, we work just with the aligned CCSs in valid cells, so retain only those:

In [51]:

```
df_ccs = df_ccs.query('valid_cell')
```

##### Per-gene barcodes / UMI sampling¶

We now want to examine how many times the typical cell barcode and UMI is observed, and also see if we expect to gain additional cell barcodes / UMIs if we sequence more.

The UMIs are shorter than the cell barcodes, and in reality the cell barcodes can be thought of as an addition to the UMI since individual molecules must come from the same cell **and** have the same UMI.
So we create a new variable called *UMI\_long* that is the concatenation of the UMI and the cell barcode--and then use this in place of just the UMI for identifying molecules:

In [52]:

```
df_ccs['UMI_long'] = df_ccs.UMI + df_ccs.barcode
```

Now for both these longer UMIs and the cell barcodes, we plot how many times each is observed and rarefaction plots of the saturation for each:

In [53]:

```
max_obs = 4 # plot up to this many observations

for prop, proplabel in [('barcode', 'cell barcode'), ('UMI_long', 'UMI')]:
    
    print("Number of times each {0} observed:".format(proplabel))
    df = (df_ccs
         .groupby(['gene_aligned_target', prop], as_index=False)
         .agg({'CCS':'count'})
         .rename(columns={'CCS':'times_observed', prop:'number'})
         .groupby(['gene_aligned_target', 'times_observed'], as_index=False)
         .agg('count')
         .assign(times_observed=lambda x:
                 x.times_observed
                 .clip(0, max_obs)
                 .map(lambda n: str(n) if n < max_obs else '{0}+'.format(max_obs)))
        )
    nobservedplot = os.path.join(analysisdir, '{0}_nobserved.pdf'.format(proplabel))
    (ggplot(df, aes('times_observed', 'number')) +
        geom_bar(stat='identity') +
        facet_wrap('gene_aligned_target', nrow=1) +
        xlab('times {0} observed'.format(proplabel)) +
        ylab('number of {0}s'.format(proplabel))
        ).save(nobservedplot, width=13, height=3)
    showPDF(nobservedplot)
    
    print("Rarefaction plot for {0}s:".format(proplabel))
    rarefyplot = os.path.join(analysisdir, '{0}_rarefaction.pdf'.format(proplabel))
    dms_tools2.plot.plotRarefactionCurves(df_ccs, prop,
            rarefyplot, facet_col='gene_aligned_target',
            ylabel='number of {0}s'.format(proplabel), xlabel='number of CCSs',
            nrow=2)
    showPDF(rarefyplot)
```

```
Number of times each cell barcode observed:
```

```
Rarefaction plot for cell barcodes:
```

```
Number of times each UMI observed:
```

```
Rarefaction plot for UMIs:
```

These plots make clear that we are nowhere near saturation of the UMIs: very few are observed more than once and the rarefaction plots indicate no saturation.
Therefore, we aren't going to use the UMIs going forward, as typically very few are sample multiple time.

But for the cell barcodes, it appears that a majority of the barcodes that we observe are observed multiple times.
However, we are still not all that close to saturation of cell barcodes in the rarefaction plots--things are flattening off, but we could still clearly get more barcodes for all genes if we sequenced more.
Also, we would like multiple UMIs for individual cells to give error correction on calling that viral genotype.

##### Get just one sequence per UMI¶

Although the subsection below shows that most UMIs are only observed once, some are observed multiple times.
When we have multiple sequences for a UMI and we treat them all independently, this could skew our estimates since many of the sources of error (reverse transcription, PCR, etc) will be shared by all members of a UMI class.
We therefore want to get just **one** of each UMI for subsequent analysis.

Probably the best way to do this would be to take the consensus of the CCSs for each duplicated UMI.
However, we are going to take a simpler approach which should also work fine provided the PacBio sequencing accuracy is high:

1. Group all UMIs that differ by no more than one nucleotide and align, and keep only the one that is most abundant in the group.
2. Among the most-abundant UMIs in a group, if some correspond to full-length alignments and others don't, subset to the full-length alignment ones. The reason is that if anything is spuriously enriched during PCR, it will likely be a shorter segment.
3. After performing step 2, keep the one with the highest accuracy over the aligned gene.

Note that we do all this after grouping by the gene to which the CCS aligns, the viral barcode, and the cell barcode, since these are also a form of UMI information.
We also use the dms\_tools2.barcodes.almost\_duplicated function to group UMIs that have just one difference.
We also note how many UMIs are exactly duplicated, but do the actual grouping based on allowing one difference.
However, we report both numbers to give a sense of how the grouping increases the number of UMIs removed.

First, we tag the duplicate UMIs and plot the number of CCSs that correspond to the duplicate UMIs for each gene:

In [54]:

```
df_ccs = (
    df_ccs
    .sort_values(['full_length', 'gene_accuracy'], ascending=False)
    .assign(duplicate_UMI=lambda x: 
        x.groupby(
            ['gene_aligned_target', 'barcode', 'viral_barcode'])
        .UMI
        .apply(dms_tools2.barcodes.almost_duplicated, threshold=1))
    .assign(exact_duplicate_UMI=lambda x: x.duplicated(
        ['gene_aligned_target', 'barcode', 'viral_barcode', 'UMI']))
    )

print(f"{df_ccs.duplicate_UMI.sum()} of {len(df_ccs)} UMIS assigned as duplicates;"
      f" of these {df_ccs.exact_duplicate_UMI.sum()} are exact duplicates.")

duplicate_UMI_plot = os.path.join(analysisdir, 'duplicate_UMIs.pdf')
(ggplot(df_ccs, aes('gene_aligned_target', fill='duplicate_UMI')) +
    geom_bar(position=position_dodge()) +
    scale_fill_manual(COLOR_BLIND_PALETTE_GRAY[1 : ]) +
    theme(axis_text_x=element_text(angle=90, hjust=0.5)) +
    ylab('number of aligned CCSs')
    ).save(duplicate_UMI_plot, height=2.5, width=4)
showPDF(duplicate_UMI_plot, width=500)
```

```
28909 of 126136 UMIS assigned as duplicates; of these 25685 are exact duplicates.
```

We see that most UMIs are not duplicated, but a non-negligible number are.
Now we remove these duplicated UMIs from further analysis, meaning we will just keep the first CCS for each UMI (above we sorted so that highest accuracy full-length sequences are first for each UMI):

In [55]:

```
df_ccs = df_ccs.query('not duplicate_UMI')
```

##### Genes sequenced per cell¶

We will now examine how completely the different genes are sequenced in each cell.

Specifically, we want to determine how many genes are captured per cell, and which genes tend to be present or missing.
Note that this analysis is limited to only cells in which at least one gene is captured.

The M2 and NS2 isoforms are at very low abundance, and their sequences are actually included within the longer unspliced major M1 and NS2 isoforms.
We therefore perform the analysis both on all isoforms and just the major isoforms:

In [56]:

```
# analyze all flu genes, and just major isoforms
all_isoforms = set(mapper.targetseqs.keys())
major_isoforms = {sorted(isoforms)[0] for isoforms in mapper.target_isoforms.values()}
print("Analyzing all {0} flu genes and just the {1} major isoforms ({2})."
      .format(len(all_isoforms), len(major_isoforms), ', '.join(major_isoforms)))
```

```
Analyzing all 10 flu genes and just the 8 major isoforms (fluNP, fluPA, fluNS1, fluPB1, fluPB2, fluHA, fluM1, fluNA).
```

In [57]:

```
for isoforms, isoform_desc in [
        (all_isoforms, 'all'), (major_isoforms, 'major')]:

    print("\nAnalysis of {0} isoforms ({1} genes):"
          .format(isoform_desc, len(isoforms)))
    
    # calculate genes per cell
    genes_per_cell = (
        df_ccs
        .query('gene_aligned_target in @isoforms')
        .groupby(['barcode'])
        .agg({'gene_aligned_target':lambda x: len(x.unique())})
        .rename(columns={'gene_aligned_target':'number of flu genes'})
        )
    # calculate genes per cell conditioned on presence of each gene
    genes_per_cell_by_gene = (
        df_ccs
        .query('gene_aligned_target in @isoforms')
        .assign(has_gene=True,
                gene_aligned_target=lambda x: 
                    x.gene_aligned_target.cat.remove_unused_categories())
        [['barcode', 'gene_aligned_target', 'has_gene']]
        .groupby(['barcode', 'gene_aligned_target'])
        .agg('any').fillna(False)
        .join(genes_per_cell)
        .reset_index()
        )

    # plot distribution of genes per cell
    plotwidth = 1 + 0.2 * len(isoforms)
    genes_per_cell_plot = os.path.join(analysisdir,
            '{0}_genes_per_cell.pdf'.format(isoform_desc))
    (ggplot(genes_per_cell, aes('number of flu genes')) +
        geom_histogram(binwidth=1) +
        scale_x_continuous(breaks=range(1, len(isoforms) + 1)) +
        ylab('number of cells')
        ).save(genes_per_cell_plot, height=2.25, width=plotwidth)
    showPDF(genes_per_cell_plot, 80 * plotwidth)
    
    # plot distribution of genes per cell faceted by gene presence
    genes_per_cell_by_gene_plot = os.path.join(analysisdir,
            '{0}_genes_per_cell_by_gene.pdf'.format(isoform_desc))
    plotwidth=1 + len(isoforms)**2 * 0.1
    (ggplot(genes_per_cell_by_gene,
            aes('number of flu genes', fill='has_gene')) +
        geom_histogram(binwidth=1) +
        scale_x_continuous(breaks=range(1, len(isoforms) + 1)) +
        ylab('number of cells') +
        facet_wrap('~gene_aligned_target', nrow=2) +
        scale_fill_manual(COLOR_BLIND_PALETTE_GRAY, name='has gene')
        ).save(genes_per_cell_by_gene_plot, height=4.25, width=plotwidth)
    showPDF(genes_per_cell_by_gene_plot, 80 * plotwidth)
```

```
Analysis of all isoforms (10 genes):
```

```
Analysis of major isoforms (8 genes):
```

#### Call mutations¶

We have already technically called mutations far above when we passed a dms\_tools2.minimap2.MutationCaller to the dms\_tools2.pacbio.matchAndAlignCCS method used to create the data frame holding the results.
We now want to examine these mutations and impose criteria (such as being observed in multiple sequences) to determine which ones are real.

##### Number of mutations per gene¶

First, we simply calculate and plot the number of mutations called per aligned sequence for each gene and mutation type, also stratifying by the viral barcode:

In [58]:

```
max_muts = 3 # plot up to this many mutations

nmuts_df = (
    df_ccs
    .assign(substitution=lambda x: x.gene_aligned_mutations.apply(
                    dms_tools2.minimap2.Mutations.substitutions),
            insertion=lambda x: x.gene_aligned_mutations.apply(
                    dms_tools2.minimap2.Mutations.insertions),
            deletion=lambda x: x.gene_aligned_mutations.apply(
                    dms_tools2.minimap2.Mutations.deletions))
    .melt(id_vars=['gene_aligned_target', 'viral_barcode'],
          value_vars=['substitution', 'insertion', 'deletion'],
          var_name='mutation_type', value_name='mutation_list')
    # workaround melt / categorical bug: https://github.com/pandas-dev/pandas/issues/15853
    .assign(gene_aligned_target=lambda x: pandas.Categorical(
            x.gene_aligned_target, targetnames))
    .assign(number=lambda x: x.mutation_list.map(len).clip(0, max_muts)
            .map(lambda n: str(n) if n < max_muts else '{0}+'.format(max_muts)))
    .assign(nseqs=1) # dummy variable for counting
    .groupby(['gene_aligned_target', 'viral_barcode', 'mutation_type', 'number'])
    .agg({'nseqs':'count'})
    .assign(fraction=lambda x: # see https://stackoverflow.com/a/23377155
                x.div(df_ccs.assign(nseqs=1)
                              .groupby('gene_aligned_target')
                              .agg({'nseqs':'count'}),
                      'gene_aligned_target'))
    .reset_index()
    )

nmuts_plot = os.path.join(analysisdir, 'nmuts_plot.pdf')
(ggplot(nmuts_df, aes('number', 'fraction', fill='viral_barcode')) +
    geom_bar(stat='identity') + 
    facet_grid('mutation_type ~ gene_aligned_target') +
    xlab('number of mutations') +
    ylab('fraction of sequences') +
    scale_fill_manual(COLOR_BLIND_PALETTE_GRAY[1 : ])
    ).save(nmuts_plot, width=12, height=7)
showPDF(nmuts_plot)
```

We see above that we often call deletions for PB2, PB1, and PA.
This is expected since these genes often have internal deletions.
We also call a lot of deletions for HA, which is less expected.
For the other genes, deletions are rarely called.

We don't see many insertions (as perhaps expected).

We see quite a few point substitutions, with the number probably about proportional to gene length.
At our sequencing accuracy of 99.9% cutoff, we would expect about 1.5 point substitutions per gene for genes like HA, NP, and NA.
In reality it is a bit higher, but of course some of this is probably due to their also being **real** point substitutions.
In any case, the high substitution rate indicates that the sequences probably have an appreciable degree of errors from sequencing or PCR that we will need to correct somehow...

The behavior of the two different viral barcodes generally looks comparable in terms of the distribution of mutations for all genes except the polymerase ones (where we already have observed that selective enrichment of certain deletions leads to some biases) and M2 / NS2 (where the counts are very low).

##### Lengths of indels¶

Now we look at the length distribution of the insertion and deletion mutations, again stratifying by gene and viral barcode.
*A priori*, we might expect two kinds of mutations: very short indels (possibly due to sequencing errors), and longer internal deletions that are real.

In [59]:

```
indel_len_df = (
    df_ccs
    .assign(insertion=lambda x: x.gene_aligned_mutations.apply(
                    dms_tools2.minimap2.Mutations.insertions, returnval='length'),
            deletion=lambda x: x.gene_aligned_mutations.apply(
                    dms_tools2.minimap2.Mutations.deletions, returnval='length'))
    .melt(id_vars=['gene_aligned_target', 'viral_barcode'],
          value_vars=['insertion', 'deletion'],
          var_name='mutation_type', value_name='mutation_length')
    .groupby(['gene_aligned_target', 'viral_barcode', 'mutation_type'])
    .mutation_length
    .sum()
    # column of lists to long form: https://stackoverflow.com/a/27266225
    .apply(pandas.Series)
    .stack()
    .rename('length')
    .reset_index()
    # workaround melt / categorical bug: https://github.com/pandas-dev/pandas/issues/15853
    .assign(gene_aligned_target=lambda x: pandas.Categorical(
            x.gene_aligned_target, targetnames))
    ) 
    
indel_len_plot = os.path.join(analysisdir, 'indel_len_plot.pdf')
(ggplot(indel_len_df, aes('length', fill='viral_barcode')) +
    geom_histogram(aes(y='..density..')) +
    facet_grid('mutation_type ~ gene_aligned_target') +
    ylab('relative fraction') +
    scale_y_continuous(breaks=None) +
    scale_x_log10(labels=lambda x: x.astype('int') 
                  if all(x.astype('int') == x) else x) +
    scale_fill_manual(COLOR_BLIND_PALETTE_GRAY[1 : ])
    ).save(indel_len_plot, width=14, height=4)
showPDF(indel_len_plot)
```

These plots are consistent with virtually all of the insertions being very short, plausibly due to sequencing errors.
For most genes, the deletions are also very short and plausibly due to sequencing errors.
However, for some genes (especially the polymerase genes but also to some extent HA and a few others), many of the deletions are very long and unlikely to be sequencing errors.

##### Accuracy of mutations¶

Below we compute the distributions of accuracies for mutations from the Q-values, and then plot the error rate (1 minus the accuracy) for these mutations.
We also do this for **all** sites in the genes.
The goal here is to determine if the mutations are enriched at low accuracy sites where sequencing errors are more probable.

First, plot the accuracy of all sites in the aligned genes:

In [60]:

```
min_error_rate = 1e-6 # plot error rates down to this low

# data frame with accuracies of all sites
all_acc_df = (
    pandas.DataFrame({
        'accuracy':dms_tools2.pacbio.qvalsToAccuracy(df_ccs
                    .gene_qvals.apply(pandas.Series).stack().values,
                    no_avg=True)})
    .assign(error_rate=lambda x: 1 - x.accuracy.clip(upper=1 - min_error_rate))
    )

# cumulative fraction plot
all_acc_plot = os.path.join(analysisdir, 'all_acc_plot.pdf')
(ggplot(all_acc_df, aes('error_rate')) +
    stat_ecdf() +
    ylab('cumulative fraction') +
    scale_x_log10() +
    scale_y_continuous(limits=(0.5, 1)) 
    ).save(all_acc_plot, width=3, height=2)
showPDF(all_acc_plot, width=300)
```

As can be seen from the above plot, the vast majority of sites have very high accuracies.

Specifically, we will only consider mutations that have accuracies of at least 0.999 (error rates < 0.001).
This is the following fraction of all site calls:

In [61]:

```
mut_acc_threshold = 0.999
print("{0} of all called identities meet the accuracy threshold."
        .format(len(all_acc_df.accuracy[all_acc_df.accuracy >= mut_acc_threshold])
        / len(all_acc_df)))
```

```
0.9970566383706483 of all called identities meet the accuracy threshold.
```

Now we make the same plot for just the accuracy of the mutated sites (not yet applying the accuracy filter in the cell above):

In [62]:

```
mut_acc_df = (
    df_ccs
    .assign(insertion=lambda x: x.gene_aligned_mutations.apply(
                    dms_tools2.minimap2.Mutations.insertions, returnval='accuracy'),
            deletion=lambda x: x.gene_aligned_mutations.apply(
                    dms_tools2.minimap2.Mutations.deletions, returnval='accuracy'),
            substitution=lambda x: x.gene_aligned_mutations.apply(
                    dms_tools2.minimap2.Mutations.substitutions, returnval='accuracy'))
    .melt(id_vars=['gene_aligned_target', 'viral_barcode'],
          value_vars=['insertion', 'deletion', 'substitution'],
          var_name='mutation_type', value_name='accuracy')
    .groupby(['gene_aligned_target', 'viral_barcode', 'mutation_type'])
    .accuracy
    .sum()
    # column of lists to long form: https://stackoverflow.com/a/27266225
    .apply(pandas.Series)
    .stack()
    .rename('accuracy')
    .reset_index()
    # workaround melt / categorical bug: https://github.com/pandas-dev/pandas/issues/15853
    .assign(gene_aligned_target=lambda x: pandas.Categorical(
            x.gene_aligned_target, targetnames))
    .assign(error_rate=lambda x: 1 - x.accuracy.clip(upper=1 - min_error_rate))
    )

mut_acc_plot = os.path.join(analysisdir, 'mut_acc_plot.pdf')
(ggplot(mut_acc_df, aes('error_rate', color='viral_barcode')) +
    stat_ecdf() +
    facet_grid('mutation_type ~ gene_aligned_target') +
    ylab('cumulative fraction') +
    scale_x_log10() +
    scale_y_continuous(limits=(0.5, 1)) +
    scale_color_manual(COLOR_BLIND_PALETTE_GRAY[1 : ])
    ).save(mut_acc_plot, width=16, height=4)
showPDF(mut_acc_plot)
```

As can be seen above, for insertions and deletions, the accuracy of mutated sites is lower than for other sites.
This is also true, but to a much less degree, for the substitution mutations.

This provides evidence that at least a reasonable fraction of sites that we call as having mutations really are lower accuracy mis-calls, and so justifies imposing an accuracy threshold to avoid getting lots of signal for the low quality mutation calls.

#### Analyze flu sequences at cell level¶

Now we want to analyze the results at the level of cells.
In particular, we want to try to build up accurate calls of the full sequences of the flu genes in individual infected cells.
Because it appears that individual CCSs may not be sufficiently accurate to call these, we in particular look for cells where we have multiple CCSs to take the consensus of these for the cell.

Of course, this approach of building consensus of CCSs from the same cell depends on the assumption that the cell was only infected by a single virion (only sometimes true based on monocle\_analysis.ipynb...) and that *de novo* mutations did not arise rapidly in the first few rounds of genome replication after infection (probably usually true given what is known about the virus's mutation rate).

##### Aggregate data at the cell level¶

First, we aggregate the information for each cell barcode, flu gene, and viral barcode into a new data frame called *df\_cells*.
We aggregate by cell barcode and flu gene because we would like to build the consensus for each flu gene for each cell, and we aggregate by viral barcode because if genes come from different viral barcodes then we know that the cell must have been infected by multiple virions, one of each viral barcode (note that there could also be some cells infected by multiple virions of the same barcode that we won't catch).

For each cell barcode, flu gene, and viral barcode we calculate the number of valid aligned sequences and a list of the mutations called in each of these CCSs.
We also calculate the number of valid aligned sequences that are also **full length**, using the criterion for full length sequences defined earlier in this notebook (this is important since for some genes such as the polymerase ones, our sequencing may be highly biased against full-length sequences).

Also, because we know that we have very low levels of the minor isoforms M2 and NS2, and because sequences of the major isoforms M1 and NS1 contain the full sequences of M2 and NS2 as well, we also make a column called *major\_isoform* which is true only if a flu gene is a major isoform.
We will then perform many of our operations only on these major isoforms.

In [63]:

```
df_cells = (
    df_ccs
    .groupby(['barcode', 'gene_aligned_target', 'viral_barcode'])
    .apply(lambda x: pandas.Series({ # following https://stackoverflow.com/a/47103408
        'n_sequences':x.gene.count(),
        'n_sequences_full_length':x.full_length.sum(),
        'mutation_list':list(x.gene_aligned_mutations)
        }))
    .query('n_sequences > 0')
    .reset_index()
    .assign(major_isoform=lambda x: x.gene_aligned_target.isin(major_isoforms))
    )
```

##### Genes sequenced per cell¶

Now for each cell and viral barcode, we tally how many genes are observed at least N times for several values of the threshold N.
We do this counting **only** for the major isoforms, so the most genes that can be observed are eight.
The reason that we tally the number of genes observed for several thresholds is to figure out how many cells we'll retain if we impose various different criterion for how many sequences we take the consensus of to call mutations.

We do this both for all sequences and for full-length sequences only.

In [64]:

```
# tally how many genes observed >= this many times for each cell
n_obs = [1, 2, 3]
n_obs_labels = ['at least {0} sequence{1}'.format(
        n, 's' if n > 1 else '') for n in n_obs]
n_obs_cols = {
    'all_lengths':['n_genes_ge_{0}'.format(n) for n in n_obs],
    'full_length':['n_genes_ge_{0}_full_length'.format(n) for n in n_obs]
    }

for nseq_col, cols in [
        ('n_sequences', n_obs_cols['all_lengths']),
        ('n_sequences_full_length', n_obs_cols['full_length'])]:
    for nobs, col in zip(n_obs, cols):
        df_cells[col] = (
            df_cells
            .assign(count_gene=lambda x: # major isoform observed enough? 
                    x.major_isoform & (x[nseq_col] >= nobs))
            .groupby(['barcode', 'viral_barcode'])
            .count_gene
            .transform('sum')
            .astype('int')
            )
```

Now plot the number of cells with a given number of genes sequenced for several thresholds for how many sequences we require per gene.
We make these plots both for any sequence, and requiring full-length sequences.

In [65]:

```
# dataframe with number of gene and full-length genes per cell
gene_counts_df = (
    df_cells
    .melt(id_vars=['barcode', 'viral_barcode'],
          value_vars=n_obs_cols['all_lengths'] + n_obs_cols['full_length'],
          var_name='minimum_observations',
          value_name='number of flu genes')
    .assign(full_length_only=lambda x: numpy.where(
          x.minimum_observations.isin(n_obs_cols['full_length']),
          'full length only', 'all lengths'))
    .assign(minimum_observations=lambda x: x.minimum_observations
          .replace(dict(zip(n_obs_cols['all_lengths'], n_obs_labels)))
          .replace(dict(zip(n_obs_cols['full_length'], n_obs_labels))))
    )

gene_counts_plot = os.path.join(analysisdir, 'gene_counts.pdf')
(ggplot(gene_counts_df, aes('number of flu genes', fill='viral_barcode')) +
    geom_bar(position=position_dodge()) +
    facet_grid('full_length_only ~ minimum_observations') +
    scale_x_continuous(limits=(0.5, len(major_isoforms) + 0.5),
        breaks=list(range(1, len(major_isoforms) + 1))) +
    ylab("number of cells") +
    scale_fill_manual(COLOR_BLIND_PALETTE_GRAY[1 : ])
    ).save(gene_counts_plot, height=4.25, width=2 * (1 + len(n_obs)))
showPDF(gene_counts_plot)
```

We see that for a reasonable number of cells, we have all 8 genes sequenced multiple times.
This might be even better than it initially seems, since (as detailed more in monocle\_analysis.ipynb) some cells don't express all viral genes.
This bodes well for calling viral genome sequences from these cells using the consensus of multiple CCSs.

#### Call mutations for viruses in cells¶

Now we try to call the "consensus" sequence for cells from the sequences we have observed from that cell.
We are trying to take advantage of the fact that we often have multiple CCSs for the same cell barcode for a gene.

##### Call consensus mutations¶

First, we use a dms\_tools2.minimap2.MutationConsensus object to call mutations for each cell barcode.
Note that this calls substitutions, deletions, and insertions separately.
We impose the accuracy threshold of 0.999 described above, and only call mutations if they are present in >30% of sequences.

In [66]:

```
# object to call mutation consensus
mutcons = dms_tools2.minimap2.MutationConsensus(
        min_acc=mut_acc_threshold,
        min_sub_frac=0.3,
        min_indel_frac=0.3)

mutation_types = ['substitutions', 'deletions', 'insertions']

# call mutations with stats on number of sequences in which observed
for mutation_type in mutation_types:
    df_cells[mutation_type] = df_cells.mutation_list.apply(
            mutcons.callConsensus, mutation_type=mutation_type,
            include_stats=True)
```

Sequences are called as *unknown* if they are not wildtype and don't have enough sequences.
We see how many barcode / gene combinations are called *unknown*:

In [67]:

```
df_cells['unknown'] = (df_cells[mutation_types] == 'unknown').any(axis=1)

unknown_seq_plot = os.path.join(analysisdir, 'unknown_seqs.pdf')
(ggplot(df_cells
        .query('major_isoform')
        .assign(n=lambda x: numpy.where(x.n_sequences > 1, '>1', '1')),
        aes('n', fill='unknown')) +
    geom_bar() +
    facet_wrap('~gene_aligned_target', nrow=1) +
    xlab('number of sequences for gene in cell') +
    scale_fill_manual(COLOR_BLIND_PALETTE_GRAY[1 :])
    ).save(unknown_seq_plot, height=1.8, width=8)
showPDF(unknown_seq_plot)
```

As expected, when there is just one sequence for a cell in a gene, most of the time it is called *unknown* since it typically isn't wildtype and our prior is to not believe any mutation if we only see it once.
This is almost completely true for the longer genes, and mostly true for the shorter ones.
This last fact makes sense, as longer genes have more opportunity for mutations.

We will ignore from further consideration any *unknown* genes / cell barcodes, since we can't call mutations in these in a useful way.

##### Frequency of mutations¶

Now we look at the frequency of the consensus mutations in each segment:

In [68]:

```
mutfreq_df = (
    df_cells
    .query('major_isoform & (not unknown)')
    .melt(id_vars=['gene_aligned_target', 'viral_barcode', 'n_sequences'],
          value_vars=mutation_types,
          var_name='mutation_type',
          value_name='mutations')
    .assign(n_mutations=lambda x: x.mutations.str.split().apply(len))
    .assign(n_mutations_ceil=lambda x: x.n_mutations.clip(0, 3).map(
            lambda n: str(n) if n < 3 else '3+'))
    .assign(n_sequences_ceil=lambda x: x.n_sequences.clip(0, 9).astype('int')
            .map(lambda n: str(n) if n < 9 else '9+'))
    .assign(mutated=lambda x: x.mutations.astype('bool'))
    # workaround melt / categorical bug: https://github.com/pandas-dev/pandas/issues/15853
    .assign(gene_aligned_target=lambda x: pandas.Categorical(
            x.gene_aligned_target, targetnames))
    )

n_consensus_muts_plot = os.path.join(analysisdir, 'n_consensus_muts.pdf')
(ggplot(mutfreq_df, aes('n_mutations_ceil')) +
    geom_bar() +
    facet_grid('gene_aligned_target ~ mutation_type') 
    ).save(n_consensus_muts_plot, width=4, height=10)
showPDF(n_consensus_muts_plot, width=400)
```

The plots above show that, as expected, mutations are quite rare.
The most common type of mutation is deletions in the polymerase genes and to a lesser extent HA.

Above we see that most cells don't have deletions even in the polymerase genes.
At first, this seems surprising given the results way above showing that many polymerase sequences have large deletions.
We reconcile this by plotting the number of sequences per cell for those with and without deletions.
We see that the cells with deletions have their polymerase genes sequenced far more often than those without deletions, explaining the preponderance of deletions among the sequences even though there isn't a preponderance among the cells.

In [69]:

```
nseqs_per_cell_by_del_plot = os.path.join(analysisdir, 'nseqs_per_cell_by_del_plot.pdf')
(ggplot(df_cells
            .query('not unknown')
            .assign(deletion=lambda x: x.deletions.astype('bool'))
            .groupby(['gene_aligned_target', 'deletion'])
            .aggregate({'barcode':'count', 'n_sequences':'sum'})
            .assign(seqs_per_cell=lambda x: x.n_sequences / x.barcode)
            .reset_index(),
        aes('gene_aligned_target', 'seqs_per_cell', fill='deletion')) +
    geom_col(position=position_dodge()) +
    scale_fill_manual(COLOR_BLIND_PALETTE_GRAY[1 : ]) +
    theme(axis_text_x=element_text(angle=90, hjust=0.5))
    ).save(nseqs_per_cell_by_del_plot, width=3.5, height=2)
showPDF(nseqs_per_cell_by_del_plot, width=500)
```

##### Mutations vs number of sequences¶

A crucial issue is the criteria we use to call consensus mutations.
If these are too lax, you might imagine that we would call more mutations if we have fewer sequences, due to some just being errors.

To look at this, we plot the fraction of genes that are called that are mutated versus the number of sequences for that gene in that cell:

In [70]:

```
mutated_vs_nseqs_plots = os.path.join(analysisdir,'mutated_vs_nseqs.pdf')
(ggplot(mutfreq_df, aes('n_sequences_ceil', fill='mutated')) +
    geom_bar(position='fill') +
    facet_grid('gene_aligned_target ~ mutation_type') +
    xlab('number of sequences') +
    ylab('fraction') +
    scale_fill_manual(COLOR_BLIND_PALETTE_GRAY[1 : ]) +
    theme(legend_position='top')
    ).save(mutated_vs_nseqs_plots, width=6, height=9)
showPDF(mutated_vs_nseqs_plots, width=400)
```

There is no trend to call more mutations when there are fewer sequences.

#### Add to cell-gene matrix annotations¶

As mentioned above have already called the valid cells with the align\_and\_annotate.ipynb notebook, and written these cells with a bit of annotations to `results/cellgenecounts/merged_humanplusflu_cells.tsv`.
(Note that we are working with the *IFN\_enriched* sample, so in the file they barcodes all have the suffix `-IFN_enriched`).
Now we want to add the genotypes / mutations that we have called by PacBio to these cell annotations:

First, we read in the existing cell annotations

In [71]:

```
print("Reading cell annotations from {0}".format(cellbarcodesfile))
annotations = pandas.read_csv(cellbarcodesfile, sep='\t').set_index('CellBarcode')   
print("Read {0} annotated cells for the {1} sample.".format(
        len(annotations.query('Sample == @sample')), sample))
assert not annotations.isna().any().any(), "can't fill NaN values"
```

```
Reading cell annotations from results/cellgenecounts/merged_humanplusflu_cells.tsv
Read 1626 annotated cells for the IFN_enriched sample.
```

Now we merge our PacBio into these existing annotations.
First, we add the number of PacBio sequences for each flu gene (**all** isoforms), broken down by viral barcode.

In [72]:

```
cell_nseqs = (
    df_cells
    .assign(colname=lambda x: x.gene_aligned_target.astype('str') + 
            x.viral_barcode.str.replace('flu', '') + '_n_PacBio_seqs')
    .assign(CellBarcode=lambda x: x.barcode + '-' + sample)
    .pivot(index='CellBarcode', columns='colname', values='n_sequences')
    .fillna(0)
    )
```

Next, we add the consensus mutations for each flu gene (just major isoforms), broken down by viral barcode, breaking mutations down into substitutions, deletions, and insertions.
We harmonize the naming of the genes with those in the existing cell annotations by renaming the major isoforms NS1 to NS and M1 to M, since the rest of the cell-gene matrix doesn't do more than 3' counting.
Also, when there are no mutations we put the string *WT*, and when there is no data the entry is *None*.

In [73]:

```
cell_mutations = (
    pandas.concat([
        df_cells
        .query('(not unknown) & major_isoform')
        .assign(CellBarcode=lambda x: x.barcode + '-' + sample)
        .assign(colname=lambda x: x.gene_aligned_target.str
                .replace('fluM1', 'fluM').replace('fluNS1', 'fluNS') + 
                x.viral_barcode.str.replace('flu', '') + '_' + mut)
        .pivot(index='CellBarcode', columns='colname', values=mut)
        for mut in mutation_types[ : 3]], axis=1)
    .replace('', 'WT')
    )
```

Finally, we merge the original cell annotations and these new ones into a single new `*.tsv` file that is appropriate for input to monocle\_analysis.ipynb notebook.

In [74]:

```
annotatedfile = os.path.join(
        os.path.dirname(cellbarcodesfile),
        'PacBio_annotated_' + os.path.basename(cellbarcodesfile))
print("Writing PacBio annotated cell information to {0}".format(annotatedfile))

(annotations
    .join(cell_nseqs).fillna(0)
    .join(cell_mutations).fillna('None')
    ).to_csv(annotatedfile, sep='\t')
```

```
Writing PacBio annotated cell information to results/cellgenecounts/PacBio_annotated_merged_humanplusflu_cells.tsv
```
