## Supplementary material for "Single-cell virus sequencing of influenza infections that trigger innate immunity": File S6

monocle\_analysis


### Table of Contents

- Analyze viral features associated with IFN induction
  - Setup for analysis
    - Load / install packages
    - Notebook-wide variables / functions
  - Get cell-gene matrices
    - Specify cell types
    - Load cell-gene matrix
  - Count cells and annotate multiplets
    - Annotate cross-celltype multiplets
    - Number of cells and multiplet frequency
    - Plot summarizing cell counts and multiplets
  - Filter multiplets and low-quality cells
    - Remove cross-celltype multiplets
    - Number of cellular and flu mRNAs, bounds for filtering
    - Plot cellular / flu mRNAs with filters
    - Filter cells with extreme mRNA amounts
  - Call infection / gene presence from canine cell thresholds
    - Constant fraction or number of mRNAs from flu?
    - Confirm equal mix of flu barcodes in canine cells
    - Look at segment frequencies
    - Get human cells for infection-status calling
    - Compute P-value flu is above background
    - Call infected cells by amount of total flu
    - Call gene presence/absence
    - Number of flu genes per cell
    - Fraction of times each flu gene missing
    - Relative expression of flu genes among infected cells
    - Call viral barcode status (co-infection)
  - Annotate interferon status
    - Calculate IFN and ISG expression
    - Correlation between type I and III IFN
    - Call IFN+ and plot IFN expression
    - Plot ISG expression
  - Unsupervised tSNE clustering
    - Estimate size factors and dispersions
    - Unsupervised tSNE clustering
  - Get viral genotypes from PacBio
    - Look at NS1 / NS2 and M1 / M2 isoforms
    - For which cells can we call viral genotypes?
      - Annotate presence of barcoded viral genes
      - Identify genes missing PacBio sequences
      - Identify cells with complete genotypes
    - Mutation types in PacBio data
  - Plot and print complete genotypes
    - Prepare to plot genotypes
    - Plot genotypes
    - Print genotypes
  - Viral features that correlate with IFN induction
    - Data frames with relevant data
    - Viral genetics and flu burden heterogeneity
    - Viral genetics and IFN induction
    - Known co-infection and IFN induction
    - Gene absence / mutations and IFN induction
    - Amount of viral mRNA and IFN induction
  - Viral features the correlate with ISG expression
    - Viral genetics and ISG expression
    - Known co-infection and ISG expression
    - Viral genetics and ISG expression
    - Amount of viral mRNA and ISG expression
  - Figures for paper

### Analyze viral features associated with IFN induction¶

This R Jupyter notebook starts with the annotated cell-gene matrix that is produced by align\_and\_annotate.ipynb and then further agumented with the PacBio information from pacbio\_analysis.ipynb.

It performs some analyses using Monocle as well as many others just using `R`.

It analyzes IFN-enriched A549 cells infected with influenza (the enrichment was done by MACS sorting at 13 hours post-infection).
The analysis also looks at the canine (MDCK) cell spike-ins used to estimate background amounts of flu in non-infected cells.

The main goal is to identify viral features associated with IFN induction

Analysis by Alistair Russell and Jesse Bloom.

#### Setup for analysis¶

##### Load / install packages¶

Load or install the necessary `R` packages, print session information that describes the packages / versions used.

In [1]:

```
options(warn=-1) # suppress warnings that clutter output

# install R packages
r_packages <- c(
  "ggplot2", "ggExtra", "gridExtra", "cowplot", "xtable",
  "scales", "reshape2", "dplyr", "magrittr", "rmarkdown",
  "IRdisplay", "psych", "qlcMatrix", "colorRamps", "ggpubr",
  "tidyverse", "RColorBrewer", "naturalsort", "grid", 
  "DescTools", "stringr", "ggrepel", "assertr", "gggenes",
  "ggsignif")
suppressMessages(invisible(
  lapply(r_packages, library, character.only=TRUE)))

# install Bioconductor packages
bioc_packages <- c("monocle", "piano", "qvalue", "Biostrings",
  "genbankr")
suppressMessages(invisible(
  lapply(bioc_packages, library, character.only=TRUE)))

sessionInfo()
```

```
R version 3.4.3 (2017-11-30)
Platform: x86_64-conda_cos6-linux-gnu (64-bit)
Running under: Ubuntu 14.04.5 LTS

Matrix products: default
BLAS/LAPACK: /fh/fast/bloom_j/software/conda/envs/BloomLab/lib/R/lib/libRblas.so

locale:
 [1] LC_CTYPE=en_US.UTF-8       LC_NUMERIC=C              
 [3] LC_TIME=en_US.UTF-8        LC_COLLATE=en_US.UTF-8    
 [5] LC_MONETARY=en_US.UTF-8    LC_MESSAGES=en_US.UTF-8   
 [7] LC_PAPER=en_US.UTF-8       LC_NAME=C                 
 [9] LC_ADDRESS=C               LC_TELEPHONE=C            
[11] LC_MEASUREMENT=en_US.UTF-8 LC_IDENTIFICATION=C       

attached base packages:
 [1] splines   stats4    parallel  grid      stats     graphics  grDevices
 [8] utils     datasets  methods   base     

other attached packages:
 [1] genbankr_1.6.1      Biostrings_2.46.0   XVector_0.18.0     
 [4] IRanges_2.12.0      S4Vectors_0.16.0    qvalue_2.10.0      
 [7] piano_1.18.1        monocle_2.6.4       DDRTree_0.1.5      
[10] irlba_2.3.2         VGAM_1.0-6          Biobase_2.38.0     
[13] BiocGenerics_0.24.0 ggsignif_0.4.0      gggenes_0.3.1      
[16] assertr_2.5         ggrepel_0.8.0       DescTools_0.99.25  
[19] naturalsort_0.1.3   RColorBrewer_1.1-2  forcats_0.2.0      
[22] stringr_1.3.1       purrr_0.2.4         readr_1.1.1        
[25] tidyr_0.7.2         tibble_1.4.1        tidyverse_1.2.1    
[28] ggpubr_0.1.8        colorRamps_2.3      qlcMatrix_0.9.7    
[31] sparsesvd_0.1-4     slam_0.1-43         Matrix_1.2-12      
[34] psych_1.8.4         IRdisplay_0.4.4     rmarkdown_1.10     
[37] magrittr_1.5        dplyr_0.7.6         reshape2_1.4.3     
[40] scales_1.0.0        xtable_1.8-2        cowplot_0.9.3      
[43] gridExtra_2.3       ggExtra_0.8         ggplot2_3.1.0      

loaded via a namespace (and not attached):
  [1] readxl_1.0.0               uuid_0.1-2                
  [3] backports_1.1.2            fastmatch_1.1-0           
  [5] plyr_1.8.4                 igraph_1.2.2              
  [7] repr_0.12.0                lazyeval_0.2.1            
  [9] densityClust_0.3           BiocParallel_1.12.0       
 [11] GenomeInfoDb_1.14.0        fastICA_1.2-1             
 [13] digest_0.6.13              htmltools_0.3.6           
 [15] viridis_0.5.1              gdata_2.18.0              
 [17] memoise_1.1.0              BSgenome_1.46.0           
 [19] cluster_2.0.6              limma_3.34.9              
 [21] ggfittext_0.6.0            modelr_0.1.1              
 [23] matrixStats_0.54.0         docopt_0.6                
 [25] prettyunits_1.0.2          colorspace_1.3-2          
 [27] blob_1.1.0                 rvest_0.3.2               
 [29] haven_1.1.0                crayon_1.3.4              
 [31] RCurl_1.95-4.11            jsonlite_1.5              
 [33] bindr_0.1.1                VariantAnnotation_1.24.5  
 [35] glue_1.2.0                 gtable_0.2.0              
 [37] zlibbioc_1.24.0            DelayedArray_0.4.1        
 [39] pheatmap_1.0.10            mvtnorm_1.0-8             
 [41] DBI_0.7                    relations_0.6-8           
 [43] miniUI_0.1.1.1             Rcpp_0.12.18              
 [45] progress_1.2.0             viridisLite_0.3.0         
 [47] bit_1.1-12                 foreign_0.8-69            
 [49] httr_1.3.1                 fgsea_1.4.1               
 [51] FNN_1.1.2.1                gplots_3.0.1              
 [53] XML_3.98-1.9               pkgconfig_2.0.1           
 [55] manipulate_1.0.1           AnnotationDbi_1.40.0      
 [57] tidyselect_0.2.3           rlang_0.2.2               
 [59] munsell_0.5.0              cellranger_1.1.0          
 [61] tools_3.4.3                cli_1.0.0                 
 [63] RSQLite_2.0                broom_0.4.3               
 [65] evaluate_0.10.1            bit64_0.9-7               
 [67] knitr_1.18                 caTools_1.17.1.1          
 [69] RANN_2.6                   bindrcpp_0.2.2            
 [71] nlme_3.1-131               mime_0.5                  
 [73] xml2_1.1.1                 biomaRt_2.34.2            
 [75] compiler_3.4.3             rstudioapi_0.7            
 [77] marray_1.56.0              stringi_1.2.4             
 [79] GenomicFeatures_1.30.3     lattice_0.20-35           
 [81] HSMMSingleCell_0.112.0     pillar_1.0.1              
 [83] combinat_0.0-8             data.table_1.10.4-3       
 [85] bitops_1.0-6               rtracklayer_1.38.3        
 [87] GenomicRanges_1.30.3       httpuv_1.3.5              
 [89] R6_2.2.2                   RMySQL_0.10.15            
 [91] KernSmooth_2.23-15         boot_1.3-20               
 [93] MASS_7.3-48                gtools_3.8.1              
 [95] assertthat_0.2.0           SummarizedExperiment_1.8.1
 [97] rprojroot_1.3-1            withr_2.1.1               
 [99] GenomicAlignments_1.14.2   Rsamtools_1.30.0          
[101] mnormt_1.5-5               GenomeInfoDbData_1.0.0    
[103] expm_0.999-3               hms_0.4.0                 
[105] IRkernel_0.8.11            Rtsne_0.13                
[107] pbdZMQ_0.3-0               sets_1.0-18               
[109] shiny_1.0.5                lubridate_1.7.1
```

##### Notebook-wide variables / functions¶

Define some variables and functions that are used throughout the rest of the notebook.

In [2]:

```
# http://www.cookbook-r.com/Graphs/Colors_(ggplot2)/#a-colorblind-friendly-palette
# The palette with grey:
cbPalette <- c("#999999", "#E69F00", "#56B4E9", "#009E73", "#F0E442", 
               "#0072B2", "#D55E00", "#CC79A7")
# The palette with black
cbbPalette <- c("#000000", "#E69F00", "#56B4E9", "#009E73", "#F0E442", 
                "#0072B2", "#D55E00", "#CC79A7")

# plots will be saved here
plotsdir <- './results/plots/'
if (!dir.exists(plotsdir)) 
  dir.create(plotsdir)    
    
# figures for paper will be saved here
figsdir <- './paper/figures/single_cell_figures'
if (!dir.exists(figsdir))
  dir.create(figsdir)

saveShowPlot <- function(p, width, height, isfig=FALSE) {
  # save plot with filename of variable name with dots replaced by _, then show
  # if *isfig* is TRUE, then also saves a PDF to *figsdir*
  pngfile <- file.path(plotsdir, sprintf("%s.png", 
    gsub("\\.", "_", deparse(substitute(p)))))
  figfile <- file.path(figsdir, sprintf("%s.pdf", 
    gsub("\\.", "_", deparse(substitute(p)))))
  ggsave(pngfile, plot=p, width=width, height=height, units="in")
  if (isfig)
    ggsave(figfile, plot=p, width=width, height=height, units="in")
  display_png(file=pngfile, width=width * 90)
}
    
fancy_scientific <- function(x, parse.str=TRUE, digits=NULL) {
  # scientific notation formatting, based loosely on https://stackoverflow.com/a/24241954
  # if `parse.str` is TRUE, then we parse the string into an expression
  # `digits` indicates how many digits to include
  x %>% format(scientific=TRUE, digits=digits) %>% gsub("^0e\\+00","0", .) %>%
    gsub("^1e\\+00", "1", .) %>% gsub("^(.*)e", "'\\1'e", .) %>% 
    gsub("e\\+","e", .) %>% gsub("e", "%*%10^", .) %>%
    gsub("^\'1\'\\%\\*\\%", "", .) %>% {if (parse.str) parse(text=.) else .}
}
```

#### Get cell-gene matrices¶

##### Specify cell types¶

We have two cell types in the experiments, and there is a separate cell-gene matrix for each cell type.
The two cell types are:

- *humanplusflu*: A549 cells infected with influenza
- *canine*: MDCK cells spiked in as a control to estimate leakage / contamination rate.

For most analyses, the *humanplusflu* cell types is the one of interest, the *canine* cells are the "other" cell type used as a control.
We therefore specify variables giving the celltype of interest and the other cell type.

We also specify which cell type has PacBio annotations (which are processed when reading the cell-gene matrix).
This is only the *humanplusflu* cells.

In [3]:

```
celltype_interest <- "humanplusflu"
celltype_other <- "canine"

celltypes <- c(celltype_interest, celltype_other)

celltype_pacbio_annotated <- setNames(c(TRUE, FALSE), celltypes)

# prettier cell-type names for plots
celltypes_pretty <- setNames( 
  lapply(celltypes, function (c) gsub("plus", " + ", c)),
  celltypes) %>% unlist
```

##### Load cell-gene matrix¶

We load the cell-gene matrices that were created previously by align\_and\_annotate.ipynb, with the cells further annotated with mutations identified by pacbio\_analysis.ipynb.
These matrices include all cells identified by cellranger.

Note that the cells are annotated by how many flu reads contain the synonymous viral barcoces near the 3' ends by align\_and\_annotate.ipynb after the cellranger analysis.
The cell type of interest is then further annotated with mutations identified by the PacBio analysis in pacbio\_analysis.ipynb.

In [4]:

```
# Cell gene matrices in this directory
matrixdir <- "./results/cellgenecounts/"

# Read cell-gene matrices into a vector named by cell type.
# These are in `matrixdir` with names like "merged_humanplusflu_matrix.mtx",
# and for PacBio annotated cells "PacBio_annotated_merged_humanplusflu_cells.tsv"
all_cells <- lapply(
  setNames(celltypes, celltypes),
  function(celltype) {
    if (celltype_pacbio_annotated[[celltype]]) {
      cellsfile <- paste("PacBio_annotated_merged", celltype, "cells.tsv", sep="_")
    } else {
      cellsfile <- paste("merged", celltype, "cells.tsv", sep="_")
    }
    newCellDataSet(
      readMM(file.path(matrixdir,
        paste("merged", celltype, "matrix.mtx", sep="_"))),
      phenoData=new(
        "AnnotatedDataFrame", data=read.delim(file.path(matrixdir, cellsfile),
          na.strings=c("None"))), # entries of None indicate NA
      featureData=new(
        "AnnotatedDataFrame",
        data=read.delim(file.path(matrixdir,
          paste("merged", celltype, "genes.tsv", sep="_")))
        ),
      expressionFamily=negbinomial.size()
      ) 
    }
  )
```

#### Count cells and annotate multiplets¶

We determine the number of cells of each type as called by the 10X pipeline in align\_and\_annotate.ipynb.
We separate the cells by whether they are "pure" of one type, or whether they are a cross-celltype multiplet (mix of *humanplusflu* and *canine*).

##### Annotate cross-celltype multiplets¶

We first look for cell barcodes that are called as both *humanplusflu* and *canine* cells.
There are cross-species multiplets, and should be filtered from downstream analyses.
Their identification also provides an estimate of the rate of multiplets.

We annotate whether a cell is human / canine by adding a new *pData* attribute called *cross\_celltype\_multiplet*.

In [5]:

```
# Annotate cross-celltype multiplets
for (celltype in celltypes) {
  # get all barcodes for the other cell type  
  other_cellbarcodes <- lapply(
      celltypes,
      function (other_celltype) {
        if (celltype != other_celltype) { 
          pData(all_cells[[other_celltype]])$CellBarcode %>% as.character
        }
      }
      ) %>%
    unlist 
  # mark shared barcodes as multiplets  
  pData(all_cells[[celltype]])$cross_celltype_multiplet <- all_cells[[celltype]] %>%
    pData %$% 
    CellBarcode %in% other_cellbarcodes
}
```

##### Number of cells and multiplet frequency¶

Tabulate the number of cells that are pure of each type, plus the cross-celltype multiplets.
The table below shows that most of the cells are *humanplusflu*.
This is expected, since the canine cells were spiked in at a relatively low fraction (targeting about 5 to 10%).
There are a modest number of cross-celltype multiplets.

In [6]:

```
tab_cellcounts <- lapply(
    celltypes,
    function (celltype) {
      all_cells[[celltype]] %>% 
      pData %>% 
      mutate(celltype=celltype) %>%
      group_by(celltype, cross_celltype_multiplet) %>% 
      summarize(ncells=n()) %>%
      ungroup
    }) %>%
  bind_rows %>%
  mutate(celltype=ifelse(cross_celltype_multiplet, "cell-type mix", celltype),
         celltype_pretty=ifelse(cross_celltype_multiplet, "cell-type mix",
                                celltypes_pretty[celltype])) %>%
  select(-cross_celltype_multiplet) %>%
  unique %>%
  transform(celltype_pretty=factor(celltype_pretty,
            c(celltypes_pretty, "cell-type mix")))

tab_cellcounts
```

| celltype | ncells | celltype\_pretty |
| --- | --- | --- |
| humanplusflu | 1614 | human + flu |
| cell-type mix | 12 | cell-type mix |
| canine | 50 | canine |

We estimate the multiplet frequency from the number of *canine* cells, *humanplusflu* cells, and cross-celltype multiplets using the function described in Bloom (2018, DOI 10.1101/293639).
This calculation estimates the multiplet frequency at $\approx$10%.

In [7]:

```
# Function to compute multiplet frequency from:
# https://github.com/jbloomlab/multiplet_freq/blob/master/calcmultiplet_R.ipynb
# Note that for this function, n1 and n2 are droplets with any cells from that
# host (including multiplets).
multiplet_freq <- function(n1, n2, n12) {
  n <- n1 * n2 / n12
  mu1 <- -log((n - n1) / n)
  mu2 <- -log((n - n2) / n)
  mu <- mu1 + mu2
  return (1 - mu * exp(-mu) / (1 - exp(-mu)))
}

tab_multiplet <- tab_cellcounts %>% 
  mutate(celltype=str_replace(celltype, "cell-type ", "celltype_")) %>%
  select(-celltype_pretty) %>%
  spread(celltype, ncells) %>%
  mutate(n1=get(celltype_interest) + celltype_mix,
         n2=get(celltype_other) + celltype_mix,
         n12=celltype_mix,
         multiplet_freq=multiplet_freq(n1, n2, n12))

tab_multiplet
```

| canine | celltype\_mix | humanplusflu | n1 | n2 | n12 | multiplet\_freq |
| --- | --- | --- | --- | --- | --- | --- |
| 50 | 12 | 1614 | 1626 | 62 | 12 | 0.1071366 |

##### Plot summarizing cell counts and multiplets¶

Finally, we make a plot that shows number of pure cells of each type and cross-celltype multiplets, and is annnotated with the estimated multiplet frequency:

In [8]:

```
p_cellcounts <- ggplot(
    tab_cellcounts,
    aes(x=celltype_pretty, y=ncells)) +
  geom_bar(width=0.75, position="dodge", stat="identity") +
  geom_text(aes(label=ncells), hjust=-0.1) +
  scale_y_continuous(name="number of cells",
    expand=expand_scale(c(0.02, 0.23))) +
  scale_x_discrete(name=NULL) + 
  theme(axis.text.x=element_blank(),
        axis.ticks.x=element_blank(),
        axis.text=element_text(size=14)) +
  coord_flip() +
  annotate("text", x=2.5, y=400,
           label=sprintf("multiplet rate = %.0f%%",
             100 * tab_multiplet$multiplet_freq),
           hjust=0,
           fontface="italic")

saveShowPlot(p_cellcounts, width=3.5, height=0.5 * (1 + length(celltypes)))
```

#### Filter multiplets and low-quality cells¶

In the previous section, we estimated the multiplet frequency.

However, we can also try to filter the multiplets and other low-quality cells. We do this in two ways:

1. We can immediately remove any of the cross-celltype multiplets, as those are obviously multiplets.
2. We remove cells that have much more or less than the typical number of cellular mRNAs detected for that cell type. This procedure is recommended in the Monocle documentation, and will remove some multiplets (since they will typically have lots of mRNA counts since they represent multiple cells), as well as some low-quality cells with little mRNA detected.

##### Remove cross-celltype multiplets¶

First, we remove all of the cross-celltype multiplets from the dataset:

In [9]:

```
all_cells <- lapply(
  all_cells,
  function (c) {
      c[, ! pData(c)$cross_celltype_multiplet]
  }
  )
```

##### Number of cellular and flu mRNAs, bounds for filtering¶

Annotate cells with the number of total mRNAs, cellular mRNAs, total flu mRNAs, flu mRNAs for each gene, and fraction of total mRNA from flu.
Note that for the *canine* cells, the flu genes are **not** in the reference genome, as the canine cells weren't infected. Nonetheless, they will have a small fraction of flu reads due to leakage.
We get their number of flu reads from the pre-annotation of the cell-gene matrix performed by align\_and\_annotate.ipynb.
For the *humanplusflu* cells, we calculate the flu reads from the cell-gene matrix.

In [10]:

```
# names of the flu genes
# use "flu" prefix so "NA" isn't interpreted as not any
flugenes <- c("fluPB2", "fluPB1", "fluPA", "fluHA",
               "fluNP", "fluNA", "fluM", "fluNS")
flugenes_noprefix <- sapply(flugenes, function (x) gsub("flu", "", x)) %>% as.vector

# annotate by number of total, cellular, and flu mRNAs, and frac from flu
all_cells <- sapply(
  names(all_cells),
  function (name) {
    c <- all_cells[[name]]
    pData(c)$total_mRNAs <- Matrix::colSums(exprs(c))
      
    pData(c)$cell_mRNAs <- Matrix::colSums(exprs(c[
      row.names(subset(fData(c), !(gene_short_name %in% flugenes))),]))
      
    flu_indices <- subset(fData(c), (gene_short_name %in% flugenes))
    if (name == celltype_interest) {
      # flu genes in cell-gene matrix, so calculate flu mRNAs from that
      stopifnot(nrow(flu_indices) == length(flugenes))
      pData(c)$flu_mRNAs <- Matrix::colSums(exprs(c[
        row.names(flu_indices),]))
      for (g in flugenes) {
        pData(c)[[paste0(g, "_mRNAs")]] = Matrix::colSums(
          exprs(c[row.names(subset(fData(c), gene_short_name == g)), ]))    
      }
    } else {
      # use flu count set during custom annotation of cell-gene matrix
      stopifnot(name == celltype_other)
      pData(c)$flu_mRNAs <- pData(c)$total_annotated_flu
      for (g in flugenes) {
        pData(c)[[paste0(g, "_mRNAs")]] = pData(c)[[
          paste0("annotated_", g)]]
      }
    }

    pData(c)$frac_mRNA_from_flu <- pData(c)$flu_mRNAs / pData(c)$total_mRNAs

    return(c)
  },
  simplify=FALSE,
  USE.NAMES=TRUE
  )
```

Below we calculate the fraction of **all** mRNA that is from flu:

In [11]:

```
lapply(
  all_cells,
  function (c) {
    pData(c) %>% 
      summarize(n_flu_mRNAs=sum(flu_mRNAs),
                n_total_mRNAs=sum(total_mRNAs),
                total_frac_flu=n_flu_mRNAs / n_total_mRNAs)
  }) %>%
  bind_rows(.id="cell_type")
```

| cell\_type | n\_flu\_mRNAs | n\_total\_mRNAs | total\_frac\_flu |
| --- | --- | --- | --- |
| humanplusflu | 2393061 | 41801983 | 0.05724755 |
| canine | 3472 | 516304 | 0.00672472 |

Now we compute the median number of cellular mRNAs and exclude cells with substantially more or less than this median by setting upper and lower bounds.
We set a *filtered* flag for all cells outside these bounds, as well as for any cells that are cross-celltype multiplets:

In [12]:

```
bottom_bound = 2 # exclude if this many fold less than median
top_bound = 2 # exclude if this many fold more than median

# annotate cells with bounds
all_cells <- lapply(
  all_cells,
  function (c) {
    pData(c)$median_cell_mRNAs <- c %>%
      pData %>%
      mutate(median=median(cell_mRNAs)) %>%
      select(median) %>%
      unlist
    pData(c)$lower_bound_cell_mRNAs <- pData(c)$median_cell_mRNAs / bottom_bound
    pData(c)$upper_bound_cell_mRNAs <- pData(c)$median_cell_mRNAs * top_bound
    pData(c)$filtered <- (
      pData(c)$cell_mRNAs < pData(c)$lower_bound_cell_mRNAs |
      pData(c)$cell_mRNAs > pData(c)$upper_bound_cell_mRNAs)
    return(c)
    }
    )

# print the bounds and number of filtered cells
lapply(
  all_cells,
  function (c) {
    pData(c) %>% 
      summarize(median_cell_mRNAs=mean(median_cell_mRNAs),
                lower_bound_cell_mRNAs=mean(lower_bound_cell_mRNAs),
                upper_bound_cell_mRNAs=mean(upper_bound_cell_mRNAs),
                n_retained=sum(! filtered),
                n_filtered=sum(filtered))
  }
  ) %>%
  bind_rows(.id="cell_type")
```

| cell\_type | median\_cell\_mRNAs | lower\_bound\_cell\_mRNAs | upper\_bound\_cell\_mRNAs | n\_retained | n\_filtered |
| --- | --- | --- | --- | --- | --- |
| humanplusflu | 23638.0 | 11819.00 | 47276 | 1490 | 124 |
| canine | 10210.5 | 5105.25 | 20421 | 49 | 1 |

##### Plot cellular / flu mRNAs with filters¶

Plot the number of cellular and viral mRNAs and show the filtering.
Each point is a cell, the green lines show the density over number of cellular mRNAs per cell, and the magenta dotted lines show the lower and upper bounds for filtering.
This plot does **not** show the cross-celltype multiplets, which have already been filtered above.

In [13]:

```
# combine data for cell types to plot
mRNA_counts_data <- all_cells %>%
  lapply(function (c) {pData(c)}) %>%
  bind_rows(.id="celltype") %>%
  mutate(celltype_pretty=celltypes_pretty[celltype]) %>%
  transform(celltype_pretty=factor(celltype_pretty, celltypes_pretty)) 

# create scatter plot
scatterplot <- ggplot(mRNA_counts_data,
    aes(cell_mRNAs, flu_mRNAs), color=cbbPalette[[1]]) +
  geom_point(alpha=0.3) 

# density plot, get value of maxy as here: https://stackoverflow.com/a/10659563
densityplot <- eval(substitute(
  {stat_density(aes(x=cell_mRNAs, y=maxy*(..scaled..)),
                geom="line", color=cbbPalette[[4]])},
  list(maxy=layer_scales(scatterplot)$y$range$range[2])))

# combine into faceted plot
p_flu_vs_cell <- scatterplot + densityplot + 
  theme(axis.text.x=element_text(angle=90, vjust=0.5, hjust=1)) +  
  geom_vline(aes(xintercept=lower_bound_cell_mRNAs), 
                 color=cbbPalette[[8]], linetype='dashed') +
  geom_vline(aes(xintercept=upper_bound_cell_mRNAs),
                 color=cbbPalette[[8]], linetype='dashed') +
  facet_wrap(~ celltype_pretty, scales='free_x') + 
  xlab("cellular mRNAs per cell") + 
  ylab("flu mRNAs per cell") +
  theme(strip.text=element_text(size=14), legend.position='none')

saveShowPlot(p_flu_vs_cell, width=6, height=3)
```

A key point in the above filtering is that we filtered based on the number of **cellular** mRNAs rather than total mRNAs.
That approach makes sense if the flu RNAs are extra in additional to the cellular mRNAs, but less sense if they replace the cellular mRNAs so that the total mRNA of infected cells is similar to uninfected cells.
In the latter case, filtering out low cellular mRNA cells would preferentially remove infected cells.

In our prior work (Russell et al (2018)), it seemed that viral mRNAs were mostly additional to cellular ones, supporting the idea of filtering on cellular mRNAs.
To confirm that it is also the case here, we correlate the fraction of mRNA from flu with total and cellular mRNA.
We see that for the infected cells (*humanplusflu*), it is much more correlated with total mRNA, with infected cells (lots of mRNA from flu) tending to have more total mRNA.
Therefore, it makes more sense to filter on cellular mRNA as we have done above since it appears that highly infected cells tend to have similar amounts of cellular mRNA but more total mRNA.

In [14]:

```
# creates string of correlation (R and P-value)
cor_string <- function (x1, x2) {
  cor <- cor.test(x1, x2, method='pearson')
  sprintf("R = %.2f (P = %.1g)", cor$estimate, cor$p.value)
}

mRNA_counts_data %>%
  group_by(celltype) %>%
  summarize(fracflu_vs_total_mRNA=cor_string(
              frac_mRNA_from_flu, total_mRNAs),
            fracflu_vs_cellular_mRNA=cor_string(
              frac_mRNA_from_flu, cell_mRNAs))
```

| celltype | fracflu\_vs\_total\_mRNA | fracflu\_vs\_cellular\_mRNA |
| --- | --- | --- |
| canine | R = -0.19 (P = 0.2) | R = -0.19 (P = 0.2) |
| humanplusflu | R = 0.26 (P = 4e-27) | R = -0.12 (P = 3e-06) |

##### Filter cells with extreme mRNA amounts¶

We now remove the cells that we have defined above for filtering based on having far more or less mRNA than the typical cell of that type:

In [15]:

```
all_cells <- lapply(
  all_cells,
  function (c) {
      c[, ! pData(c)$filtered]
  }
  )
```

#### Call infection / gene presence from canine cell thresholds¶

We want to determine which cells we think are truly infected with flu various versus just having flu mRNA they have picked up from lysed cells.
We also want to call presence of each individual gene in infected cells.
The *canine* cells provide a way to examine this, since they are **not** infected with flu and so should provide a baseline for how much viral mRNA to expect in non-infected cells.

##### Constant fraction or number of mRNAs from flu?¶

We can imagine two relatively simple scenarios to explain the amount of contaminating flu mRNA that cells simply pick up from the environment:

1. Cells get a constant number of contaminating flu reads.
2. Cell get a constant fraction of number of contaminating flu reads.

In the first case, the number of flu mRNAs should be independent of the number of total mRNAs.
In the second case, the number of flu mRNAs should increase linearly with the number of total mRNAs.

First, we make a plot of the relationship between the number of flu mRNAs and total mRNAs for the canine cells.
As that plot shows, the relationship appears linear, suggesting the constant fraction model:

In [16]:

```
threshold_data <- all_cells[[celltype_other]] %>%
  pData %>% 
  rename_all(funs(str_replace(., "annotated_", ""))) 
         
p_threshold_linear <- ggplot(
    threshold_data, aes(total_mRNAs, flu_mRNAs)) +
  geom_point() +
  geom_smooth(method='lm')

saveShowPlot(p_threshold_linear, width=3.5, height=3)
```

Next, we estimate the number or fraction of mRNA in these *canine* cells that comes from flu.

Under the constant number of contaminating flu mRNAs model, this is just the average number of flu mRNAs per canine cell.

Under the constant fraction of contaminating flu mRNAs model, there are two ways we could compute the average fraction:

1. Simply averaging over all cells
2. Taking a weighted average over cells, so that cells with more mRNA are weighted more strongly.

Below we see that these approaches for the constant fraction model given essentially identical results, so we just use the first approach:

In [17]:

```
threshold_data %>% 
  summarize(
    mean_frac_flu=mean(frac_mRNA_from_flu),
    weighted_mean_frac_flu=sum(flu_mRNAs) / sum(total_mRNAs),
    mean_n_flu=mean(flu_mRNAs)
    )
```

| mean\_frac\_flu | weighted\_mean\_frac\_flu | mean\_n\_flu |
| --- | --- | --- |
| 0.006768012 | 0.006724974 | 70.34694 |

We now annotate cells by the predicted number of mRNAs from flu under both the constant number and constant fraction model.
We also plot the relationship between the predicted and observed flu mRNAs in each model.
The plot again suggests that the constant fraction model might be better:

In [18]:

```
threshold_data <- threshold_data %>% 
  mutate(mean_frac_flu=mean(frac_mRNA_from_flu),
         mean_n_flu=mean(flu_mRNAs)) %>%
  mutate(predicted_flu_mRNAs_constant_frac=mean_frac_flu * total_mRNAs,
         predicted_flu_mRNAs_constant_number=mean_n_flu)

p_actual_predicted_flu <- ggplot(
    threshold_data %>% 
      mutate(constant_fraction=predicted_flu_mRNAs_constant_frac,
             constant_number=predicted_flu_mRNAs_constant_number) %>%
      gather(key='predict_method', value='predicted_flu_mRNAs',
             constant_fraction, constant_number),
    aes(predicted_flu_mRNAs, flu_mRNAs)) +
  geom_point() +
  facet_wrap(~ predict_method) 

saveShowPlot(p_actual_predicted_flu, width=5, height=3)
```

We now test if the predictions fit as well as expected if the data really are Poisson distributed from the predicted values under either the constant fraction or constant number model.
We do this using the Poisson deviance goodness of fit test described here.
The larger the deviance, the worse the fit.
The P-value indicates if we can reject we can reject the Poisson distributed numbers around the predicted values.

Below, we see that both models can be rejected (neither fit the data adequately).
However, going by both the deviance and the P-value, as well as the qualitative plots above, the predicted values are better under the constant fraction model.

So this provides a rationale by identifying infected *humanplusflu* cells based on the fraction of their mRNA from flu rather than the total number of mRNAs from flu.

In [19]:

```
threshold_data %>%
  mutate(constant_fraction=predicted_flu_mRNAs_constant_frac,
         constant_number=predicted_flu_mRNAs_constant_number) %>%
  gather(key='predict_method', value='predicted_flu_mRNAs',
         constant_fraction, constant_number) %>%
  mutate(dev_term=flu_mRNAs * log(flu_mRNAs / predicted_flu_mRNAs) -
                  (flu_mRNAs - predicted_flu_mRNAs)) %>%
  group_by(predict_method) %>%
  summarize(deviance=2 * sum(dev_term),
            ndegrees=n() - 1,
            P=pchisq(deviance, ndegrees, lower.tail=FALSE))
```

| predict\_method | deviance | ndegrees | P |
| --- | --- | --- | --- |
| constant\_fraction | 80.22651 | 48 | 2.427470e-03 |
| constant\_number | 159.91999 | 48 | 5.912465e-14 |

##### Confirm equal mix of flu barcodes in canine cells¶

The human cells were infected with a mix of wildtype and synonymously barcoded virus.

If the *canine* cells are just picking up environmental viral mRNA, then each should have a mix of mRNAs from these two viruses that should match the average composition of the *humanplusflu* cells.

Below we verify that this is true: we see that the fraction of flu reads with annotated wildtype or synonymous barcodes is about the same in *canine* and *humanplusflu* cells.
Therefore, for the rest of this section, we will just use the fraction estimated from the *canine* cells.

Interestingly, the table below shows that the fraction of flu reads that have an annotated barcode is different between *canine* and *humanplusflu* cells, however.
Not sure of the reason (it could have something to do with preferential leakage of defective particles), but this means that we will only extrapolate from *canine* to *humanplusflu* cells the fraction of annotated reads that are wildtype or synonymous, not the fraction of annotated reads.

In [20]:

```
lapply(all_cells, function (c) pData(c)) %>%
  bind_rows(.id="celltype") %>%
  group_by(celltype) %>%
  summarize(n_wt_flu=sum(annotated_flu_wt),
            n_syn_flu=sum(annotated_flu_syn),
            frac_flu_wt=n_wt_flu / (n_wt_flu + n_syn_flu),
            n_annotated=n_wt_flu + n_syn_flu,
            frac_annotated=n_annotated / sum(flu_mRNAs))
```

| celltype | n\_wt\_flu | n\_syn\_flu | frac\_flu\_wt | n\_annotated | frac\_annotated |
| --- | --- | --- | --- | --- | --- |
| canine | 378 | 618 | 0.3795181 | 996 | 0.2889469 |
| humanplusflu | 347502 | 625675 | 0.3570800 | 973177 | 0.4674827 |

Now we examine how well the fraction of flu reads from each barcode actually follows the predicted value assuming equal mixing.
We obviously expect more statistical noise for cells with fewer reads with annotated barcodes, so we plot the fraction from each barcode as a function of the number of barcode-annotated reads.
In the plots below, the horizontal line indicates the expected fraction from the averages.
As the plot shows, the actual fractions are clustered around the horizontal line, with the clustering getting tighter as we increase the number of flu reads.
This is as expected.

In [21]:

```
threshold_data <- threshold_data %>%
  mutate(avg_frac_flu_wt=sum(flu_wt) /
          (sum(flu_wt) + sum(flu_syn))) %>%
  mutate(n_annotated_flu=flu_wt + flu_syn,
         frac_flu_wt=flu_wt / n_annotated_flu,
         predicted_flu_wt=avg_frac_flu_wt * n_annotated_flu)

p_barcode_fraction <- ggplot(
    threshold_data, aes(n_annotated_flu, frac_flu_wt)) +
  geom_point(alpha=0.33) +
  geom_hline(aes(yintercept=avg_frac_flu_wt), color=cbbPalette[[2]]) 

saveShowPlot(p_barcode_fraction, width=3.5, height=3)
```

We now determine whether the observed "noise" in the fraction of flu with each barcode is consistent with a binomial distribution parameterized by the average fraction flu across cells.
We do this using the binomial deviance goodness of fit test, calculating the deviance using the formula provided here.
As can be seen below, we cannot reject the null hypothesis of a binomial distribution, suggesting we can interpret the binomial as a good fit, and assume equal mixing of barcodes in the *canine* cells.

In [22]:

```
threshold_data %>%
  filter(n_annotated_flu > 0) %>%
  mutate(dev_term1=ifelse(flu_wt == 0,
                          0,
                          flu_wt * log(flu_wt / predicted_flu_wt)),
         dev_term2=ifelse(flu_syn == 0,
                          0,
                          log(flu_syn / (n_annotated_flu - 
                                         predicted_flu_wt))),
         dev_term=dev_term1 + dev_term2) %>%
  summarize(deviance=2 * sum(dev_term),
            ndegrees=n() - 1,
            P=pchisq(deviance, ndegrees, lower.tail=FALSE))
```

| deviance | ndegrees | P |
| --- | --- | --- |
| 31.28426 | 48 | 0.9704415 |

##### Look at segment frequencies¶

Now we look at the individual segment frequencies to come up with thresholds to call whether each cell is expressing each segment.

First, annotate all cells by the fraction of flu mRNA that comes from each segment:

In [23]:

```
all_cells <- lapply(
  all_cells,
  function (c) {
    pData(c)[sapply(flugenes, function (x) paste0(x, "_frac"))] <- (
      pData(c)[sapply(flugenes, function (x) paste0(x, "_mRNAs"))] /
      pData(c)$flu_mRNAs)
    c    
  }    
  )
```

Now let's get the total number of flu mRNAs for each gene:

In [24]:

```
flu_gene_freqs <- lapply(all_cells, function (c) pData(c)) %>%
  bind_rows(.id="celltype") %>%
  group_by(celltype) %>%
  mutate_at(sapply(flugenes, function (x) paste0(x, "_mRNAs")),
            return) %>%
  summarize_at(c(flugenes, "flu_mRNAs"), sum)
            
flu_gene_freqs
```

| celltype | fluPB2 | fluPB1 | fluPA | fluHA | fluNP | fluNA | fluM | fluNS | flu\_mRNAs |
| --- | --- | --- | --- | --- | --- | --- | --- | --- | --- |
| canine | 102 | 73 | 81 | 247 | 672 | 647 | 780 | 845 | 3447 |
| humanplusflu | 90710 | 40843 | 54199 | 155122 | 407576 | 426672 | 378018 | 528599 | 2081739 |

Now we plot these data to see if the fractions from each gene are similar among the *canine* and *humanplusflu* cell types.
We estimate the statistical noise on each estimate from counting statistics, and plot error bars that show 1.96 times the estimated standard deviation (95% confidence interval).

Overall, the plot below shows that *humanplusflu* cells look fairly similar to the *canine* ones in the distributions of reads among flu genes.
The overall expression profiles for genes is also roughly similar to those from our previous study (Russell et al, 2018), with NS, M, NA and NP being highest, and then HA, and then the three polymerase genes.

Based on the plot below, we will use the gene frequencies in the *canine* cells to estimate cutoffs for whether each cell in *humanplusflu* is expressing a gene.

In [25]:

```
p_flu_gene_freqs <- ggplot(
    flu_gene_freqs %>% 
      gather(gene, counts, -celltype, -flu_mRNAs) %>%
      mutate(frac=counts / flu_mRNAs,
             frac_sd=1.96 * sqrt(counts) / flu_mRNAs) %>%
      transform(gene=factor(gsub("flu", "", gene), flugenes_noprefix)),
    aes(gene, weight=frac, fill=celltype)) +
  geom_bar(position=position_dodge()) +
  geom_errorbar(aes(ymin=frac - frac_sd,
                    ymax=frac + frac_sd,
                    fill=celltype),
                width=0.3, position=position_dodge(width=0.9)) +
  ylab("fraction of flu mRNAs") +
  theme(axis.text.x=element_text(angle=90,hjust=1, vjust=0.5)) +
  scale_fill_manual(values=c(cbPalette[1], cbPalette[3])) 

saveShowPlot(p_flu_gene_freqs, width=5, height=3)
```

##### Get human cells for infection-status calling¶

We will now call infection status and presence of each flu gene from the background thresholds determined from the *canine* cells.
We will do this just for the *humanplusflu* cells, as these are what we will use for downstream analyses.
So we get those cells into a new variable called `cells`.

In [26]:

```
cells <- all_cells[[celltype_interest]]
```

##### Compute P-value flu is above background¶

Above, we've determined that the *canine* cells provide a good indication background fraction of contaminating reads from flu and for each flu gene.
Now, we need to get the exact numbers for this background expection.
Cells will be called as "infected" and also for the presence of individual genes if they have significantly more than this fraction of mRNA from flu or the indicated flu gene.

In [27]:

```
thresholds <- threshold_data %>%
  mutate(flu=flu_mRNAs, total=total_mRNAs) %>%
  summarize_at(c(flugenes, "flu", "total"), sum) %>%
  gather(gene, frac, -total) %>%
  mutate(frac=frac / total,
         percent=frac * 100) %>%
  select(-total)

thresholds
```

| gene | frac | percent |
| --- | --- | --- |
| fluPB2 | 0.0001989984 | 0.01989984 |
| fluPB1 | 0.0001424204 | 0.01424204 |
| fluPA | 0.0001580281 | 0.01580281 |
| fluHA | 0.0004818882 | 0.04818882 |
| fluNP | 0.0013110481 | 0.13110481 |
| fluNA | 0.0012622740 | 0.12622740 |
| fluM | 0.0015217523 | 0.15217523 |
| fluNS | 0.0016485650 | 0.16485650 |
| flu | 0.0067249745 | 0.67249745 |

Above we noted number of flu reads per cell in the *canine* cells appear vaguely consistent with Poisson sampling relative to the expected number from a constant contaminating fraction.
So for each cell, we can calculate a P-value to reject the null hypothesis that the number of observed flu reads is consistent with them just being from the contaminating background.
This function does that test:

In [28]:

```
poisson_P <- function(n, mu){
  # Probability of observing >= `n` given Poisson with mean `mu`
  poisson.test(n, r=mu, alternative="greater")$p.value
}
```

We compute the P-value with which we can reject the null hypothesis that the amount of total flu mRNA is consistent with the background rates, both for total flu and for each gene individually, and also for all flu and by viral barcode.

In [29]:

```
for (g in c(flugenes, "flu")) {
  # get thresholds for this gene
  threshold <- thresholds %>% filter(gene == g)
  # annotate cells with P values for all flu
  pData(cells)[[paste0("has_", g, "_Pval")]] <- map2(
      pData(cells)[[paste0(g, "_mRNAs")]],
      pData(cells)$total_mRNAs * threshold$frac, 
      poisson_P) %>%
    unlist
}
```

##### Call infected cells by amount of total flu¶

Now we categorize cells as whether they are infected.
We have a lot of cells, so we compute a Q-value for whether the cell is infected.
This will allow us to call infection status at some specified false discovery rate (FDR).

In [30]:

```
pData(cells)$infected_Q <- pData(cells) %>%
  mutate(infected_Q=qvalue(has_flu_Pval)$qvalue) %$%
  infected_Q
```

Now let's plot the distribution of Q values for cells being infected (actually, we plot the $\log$ Q-value).
As is immediately apparent, the majority of cells have Q values close to 1 ($\log$s close to 0), meaning we can reject that they are infected.
But a moderate fraction of cells have very low Q-values, indicating that they are clearly infected.

In [31]:

```
p_qvals <- ggplot(
    pData(cells) %>% mutate(logQ=pmax(-100, log10(infected_Q))), 
    aes(logQ)) +
  stat_ecdf() + 
  coord_cartesian(xlim=c(-12, 0)) +
  ylab("cumulative fraction") +
  xlab("log10(Q)")

saveShowPlot(p_qvals, 4, 2.5)
```

The Q-value distributions above indicate that it doesn't matter much where we set the FDR for calling infected cells, as long as we choose some reasonably small value (e.g., $< 0.1$).
Since for our analyses it will be worse to falsely call infected cells than to miss a few, and since the distribution of flu reads per cell was not quite Poisson (and overdispersion will make the P-values liberal), we will be conservative and call infected cells at a FDR of $10^{-8}$, which corresponds to -8 on the log Q-value plot above.
Note that this is a ridiculously low FDR; however, since the number of flu reads per cell used to estimate the background isn't exactly Poisson, we wanted to be cautious and only get very clearly infected cells.
We then print some basic statistics on the infected and uninfected cells:

In [32]:

```
fdr_cutoff <- 1e-8
pData(cells)$infected <- pData(cells)$infected_Q < fdr_cutoff

pData(cells) %>%
  group_by(infected) %>%
  summarize(n=n(),
            mean_cellular_mRNAs=mean(cell_mRNAs),
            mean_total_mRNAs=mean(total_mRNAs),
            min_frac_mRNA_from_flu=min(frac_mRNA_from_flu),
            max_frac_mRNA_from_flu=max(frac_mRNA_from_flu))
```

| infected | n | mean\_cellular\_mRNAs | mean\_total\_mRNAs | min\_frac\_mRNA\_from\_flu | max\_frac\_mRNA\_from\_flu |
| --- | --- | --- | --- | --- | --- |
| FALSE | 1200 | 24792.62 | 24872.71 | 0.001678704 | 0.01048556 |
| TRUE | 290 | 25453.73 | 32300.74 | 0.010097493 | 0.65112671 |

Now we get the cutoff minimum fraction of flu at which we call cells infected.
Note that the maximimum fraction mRNA from flu in the *uninfected* cells and the minimum fraction in *infected* cells are actually very slightly different, since the statistical calling method above also takes into account the total number of reads in a cell in calling the P-value.
But they are very close, and we will make some plots above using the minimum fraction of flu above which all cells are called infected:

In [33]:

```
min_frac_flu <- pData(cells) %>% 
  filter(! infected) %$% 
  frac_mRNA_from_flu %>% 
  max

cat("All cells with >", min_frac_flu, "mRNA from flu are called infected")
```

```
All cells with > 0.01048556 mRNA from flu are called infected
```

In [34]:

```
# get data with infection status into `all_cells`
all_cells[[celltype_interest]] <- cells
 
# data for histograms
hist_data <- lapply(
  names(all_cells),
  function (celltype) {
    all_cells[[celltype]] %>%
      pData %>%
      mutate(celltype=celltype,
             celltype_pretty=celltypes_pretty[celltype],
             infected=ifelse(celltype == celltype_interest,
                      infected,
                      frac_mRNA_from_flu > min_frac_flu)) %>%
      select(celltype, celltype_pretty, frac_mRNA_from_flu, infected)
    }
  ) %>% bind_rows

# histograms for all cells
p_frac_flu_histograms <- ggplot(
    hist_data,
    aes(frac_mRNA_from_flu, fill=infected)) + 
  geom_histogram(bins=33) +
  scale_x_continuous() +
  scale_fill_manual(values=c(cbPalette[4], cbPalette[3]),
    name="classified\nas infected?") +
  ylab("number of cells") +
  xlab("frac mRNA from flu") +
  facet_wrap(~celltype_pretty, ncol=1, scales='free_y') +
  theme(strip.text=element_text(size=14), legend.position='none')

# histogram for infected cells
p_frac_flu_infected_histogram <- ggplot(
    hist_data %>% filter(celltype == celltype_interest & infected),
    aes(frac_mRNA_from_flu)) +
  geom_histogram(bins=32, fill=cbPalette[3]) +
  scale_x_continuous(name=NULL, limits=c(0, NA)) +
  ylab(NULL) +
  theme(plot.title=element_text(face='plain', vjust=-2))

# counts of cells infected / uninfected
cellcounts <- hist_data %>%
  transform(infected=factor(infected)) %>%
  group_by(celltype, infected) %>%
  summarize(n=n()) %>%
  complete(infected) %>%
  mutate(n=ifelse(is.na(n), 0, n)) %>%
  mutate(infected=infected %>% as.logical)

# merge into one figure
p_frac_flu <- ggdraw() +
  draw_plot(p_frac_flu_histograms, 0, 0, 1, 1) +
  draw_plot(p_frac_flu_infected_histogram, 0.4, 0.17, 0.6, 0.28) +
  draw_label(
    paste(cellcounts %>% filter(celltype == celltype_other & ! infected) %$% n,
      "uninfected"),
    x=0.96, y=0.91, colour=cbPalette[4], size=12, hjust=1, fontface='italic') +
  draw_label(
    paste(cellcounts %>% filter(celltype == celltype_other & infected) %$% n,
      "infected"),
    x=0.96, y=0.87, colour=cbPalette[3], size=12, hjust=1, fontface='italic') +
  draw_label(
    paste(cellcounts %>% filter(celltype == celltype_interest & ! infected) %$% n,
      "uninfected"),
    x=0.96, y=0.48, colour=cbPalette[4], size=12, hjust=1, fontface='italic') +
  draw_label(
    paste(cellcounts %>% filter(celltype == celltype_interest & infected) %$% n,
      "infected"),
    x=0.96, y=0.44, colour=cbPalette[3], size=12, hjust=1, fontface='italic')

saveShowPlot(p_frac_flu, width=3, height=4.5)
```

##### Call gene presence/absence¶

Now we want to call whether each infected cell is expressing each individual flu gene above background.
We've already used a method to control for multiple hypotheses (controlling the FDR) above, so now we make the call for each gene cell individually using just the P-values without a further multiple hypothesis test.
The reason that we do not need another multiple hypothesis test is that we are conditioning on the cells already having been called as infected.

First, we plot the distribution of P-values **among infected cells**:

In [35]:

```
pvals <- pData(cells) %>% 
  filter(infected) %>%
  select(CellBarcode, matches('_Pval')) %>%
  gather(gene, P, -CellBarcode) %>%
  mutate(gene=gsub("has_|_Pval", "", gene)) %>%
  filter(gene != "infected", gene != "flu") 

p_pvals <- ggplot(
    pvals %>% mutate(logP=pmax(-100, log10(P))), 
    aes(logP)) +
  stat_ecdf() + 
  coord_cartesian(xlim=c(-3, 0)) +
  facet_wrap(~ gene, nrow=1) +
  ylab("cumulative fraction") +
  xlab("log10(P)")

saveShowPlot(p_pvals, 10, 2.5)
```

The plot above clearly shows that it will be harder to set a clean P-value cutoff for whether cells have a specific flu gene.
For some of the more highly expressed genes (e.g., NS, M) it is pretty clear.
But for lower-expressed genes, it's harder to make a clean division.
So we'll choose a P-value cutoff of 0.05, which seems plausible given above and also means that we're pretty sure that most of the time we're not falsely calling cells as having a flu gene that they really are missing.

In [36]:

```
p_has_gene_cutoff <- 0.05

pData(cells)[map_chr(flugenes, function (g) paste0("has_", g))] <- (
  (pData(cells)[map_chr(flugenes, function(g) paste0("has_", g, "_Pval"))]
     < p_has_gene_cutoff) & pData(cells)$infected)
```

##### Number of flu genes per cell¶

Compute the number of distinct flu genes per cell based on the gene presence / absence calling above:

In [37]:

```
pData(cells)['n_flu_genes'] <- pData(cells) %>% 
  select_(.dots=lapply(flugenes, function (g) paste0("has_", g))) %>% 
  rowSums
```

Now plot the distribution of the number of flu genes per cell among the cells we have previously called as infected.

In [38]:

```
p_flugene_count <- ggplot(
    pData(cells) %>% filter(infected), aes(n_flu_genes)) +
  geom_bar(fill=cbPalette[3]) +
  ylab("number of cells") +
  geom_text(stat='count', aes(label=..count..), vjust=-0.3) +
  expand_limits(y=c(0, 190)) +
  theme(axis.text.x=element_blank(),
        axis.title.x=element_blank(),
        axis.ticks.x=element_blank(),
        axis.text.y=element_blank(),
        axis.ticks.y=element_blank())

p_fracflu_by_ngenes <- ggplot(
    pData(cells) %>% filter(infected),
    aes(factor(n_flu_genes), frac_mRNA_from_flu)) +
  geom_boxplot(outlier.size=1, outlier.alpha=0.4,
    fill=cbPalette[3], outlier.color=cbPalette[3]) +
  xlab("number of flu genes") +
  ylab("frac mRNA from flu") 

p_nflu_genes <- plot_grid(p_flugene_count, p_fracflu_by_ngenes,
  ncol=1, align='v', rel_heights=c(1, 1.3))
saveShowPlot(p_nflu_genes, width=3.5, height=4)
```

##### Fraction of times each flu gene missing¶

Here we plot and show statistics on the fraction of infected cells that have each gene.
Typically genes are present in infected cells >85% of the time but <95% of the time, which is comparable to prior estimates.

The exception is that we call almost all cells as having NP, which may just be because we have a hard time detecting infection if cells lack NP due to failure of secondary transcription.

In [39]:

```
tab_frac_has_gene <- pData(cells) %>%
  filter(infected) %>%
  rename_all(funs(str_replace_all(., 'has_flu', ''))) %>%
  select(flugenes_noprefix) %>%
  summarize_all(mean)

p_frac_has_gene <- ggplot(
    tab_frac_has_gene %>% 
      gather(gene, frac) %>%
      transform(gene=factor(gene, flugenes_noprefix)),
    aes(x=gene, y=frac)) +
  geom_bar(stat='identity', fill=cbPalette[3]) +
  geom_text(aes(label=sprintf("%0.2f", frac)), vjust=1.5, color='white') +
  theme(axis.text.x=element_text(angle=90, hjust=1, vjust=0.5),
        axis.text.y=element_blank(),
        axis.ticks.y=element_blank()) +
  ylab("fraction cells with gene") +
  geom_hline(yintercept=1, linetype='dashed', color=cbPalette[1])

saveShowPlot(p_frac_has_gene, width=3.5, height=2.75, isfig=TRUE)
```

##### Relative expression of flu genes among infected cells¶

We plot the relative expression of flu genes among the infected *humanplusflu* cells.
Note that this distribution of expressions is reasonably similar to that in Russell et al (2018) in terms of the heirarchy of which genes are most highly expressed.
Reasons that the values are not exactly identical could include: we have applied slightly different filters to call infected cells, are looking at a different timepoint, and have enriched for IFN+ cells.

In [40]:

```
p_flu_rel_expr <- ggplot(
    pData(cells) %>%
      filter(infected) %>%
      mutate_at(sapply(flugenes, function (x) paste0(x, "_mRNAs")),
                return) %>%
      select(CellBarcode, flu_mRNAs, flugenes) %>%
      gather(gene, counts, flugenes) %>%
      transform(gene=factor(gsub("flu", "", gene), flugenes_noprefix)) %>%
      mutate(frac=counts / flu_mRNAs),
    aes(gene, frac)) +
  geom_boxplot(notch=TRUE, outlier.size=1, outlier.alpha=0.4,
    fill=cbPalette[3], outlier.color=cbPalette[3]) +
  ylab("frac viral mRNA") +
  theme(axis.text.x=element_text(angle=90, hjust=1, vjust=0.5),
        axis.title.x=element_blank())
    
saveShowPlot(p_flu_rel_expr, width=3.5, height=3)
```

##### Call viral barcode status (co-infection)¶

Cells were infected with a mix of wildtype and synonymously barcoded viruses.
Here we call whether they are infected with viruses with just one barcode, or are co-infected.
Note that we expect to only identify about half (actually a few less, since the barcode mixing is not exactly even, see above) of the truly co-infected cells this way, since some will be co-infected with the same barcode.

First, we annotated the number of called flu barcodes and the "purity" of these barcoded flu reads.
The purity if the fraction of annotated barcodes that come from the more abundant variant (wildtype or synonymously barcoded).
The purity can therefore range from 0.5 (equal mix of both virus barcode variants) to 1 (only one barcode).

We then calculate the P-value with which we can reject the hypothesis that the barcodes are drawn from a distribution with a mean purity of at least 95%.
We then call as co-infected all cells that are infected and for which this P-value is $<10^{-3}$.
We also annotate each cell by whether it is infected with each barcode.

Finally, we print a table of the number of cells that we can call as coinfected this way, and also plot the cells that we called as coinfected versus their number of annotated barcodes and purity:

In [41]:

```
pData(cells)$n_annotated_flu <- pData(cells)$annotated_flu_wt +
    pData(cells)$annotated_flu_syn
pData(cells)$n_major_barcode <- pmax(pData(cells)$annotated_flu_wt,
    pData(cells)$annotated_flu_syn)
pData(cells)$purity <- pData(cells)$n_major_barcode / 
    pData(cells)$n_annotated_flu

pData(cells)$coinfected_P <- map2(
  pData(cells)$n_major_barcode, 
  pData(cells)$n_annotated_flu, 
  function(x, n) ifelse(n == 0, 1, binom.test(x, n, 0.95, "less")$p.value)
  )

# annotate whether coinfected
pData(cells)$coinfected <- pData(cells)$infected & 
  (pData(cells)$coinfected_P < 1e-3)

# annotate whether infected by each barcode
pData(cells)$infected_wt <- pData(cells)$infected &
  (pData(cells)$coinfected | 
  (pData(cells)$annotated_flu_wt >= pData(cells)$annotated_flu_syn))
pData(cells)$infected_syn <- pData(cells)$infected &
  (pData(cells)$coinfected | 
  (pData(cells)$annotated_flu_syn >= pData(cells)$annotated_flu_wt))

pData(cells) %>%
  filter(infected) %>%
  group_by(coinfected) %>%
  summarize(ncells=n(),
            mean_frac_flu=mean(frac_mRNA_from_flu))

p_coinfected <- ggplot(pData(cells) %>% filter(infected), 
    aes(x=n_annotated_flu, y=purity, color=coinfected)) + 
  geom_point() + 
  scale_color_manual(values=cbbPalette[2:3]) 

saveShowPlot(p_coinfected, 4.5, 3)
```

| coinfected | ncells | mean\_frac\_flu |
| --- | --- | --- |
| FALSE | 179 | 0.1779311 |
| TRUE | 111 | 0.2091390 |

We obviously only observe the co-infected cells with a mix of both barcodes.
We can use the `multiplet_freq` function defined above (and taken from Bloom (2018, DOI 10.1101/293639)) to estimate the actual co-infection frequency assuming Poisson infection:

In [42]:

```
tab_coinfect <- pData(cells) %>% 
  filter(infected) %>%
  mutate(viral_barcode=ifelse(coinfected, "both",
    ifelse(infected_wt, "wt", "syn"))) %>%
  group_by(viral_barcode) %>%
  summarize(ncells=n()) %>%
  spread(key=viral_barcode, value=ncells) %>%
  mutate(coinfection_freq=multiplet_freq(wt + both, syn + both, both))
```

Plot the numbers and coinfection frequency:

In [43]:

```
p_coinfect <- ggplot(
    tab_coinfect %>%
      select(-coinfection_freq) %>%
      gather(barcode, ncells) %>%
      transform(barcode=factor(barcode, c("wt", "syn", "both"))) %>%
      mutate(coinfected=ifelse(barcode == "both", TRUE, FALSE)),
    aes(x=barcode, y=ncells, fill=coinfected)) +
  geom_bar(width=0.75, position="dodge", stat="identity") +
  geom_text(aes(label=ncells), hjust=0.5, vjust=1.4, color='white') +
  scale_fill_manual(values=c(cbPalette[3], cbPalette[6])) +
  scale_y_continuous(name="number of cells",
    expand=expand_scale(c(0.02, 0.2))) +
  scale_x_discrete(name="viral barcode") + 
  theme(axis.text.y=element_blank(),
        axis.ticks.y=element_blank(),
        text=element_text(size=14),
        axis.text=element_text(size=12),
        legend.position='none') +
  annotate("text", x=3.5, y=180,
           label=sprintf(
             paste("dual-barcode coinfection: %.0f%%",
                   "estimated coinfection rate: %.0f%%",
                   sep='\n'),
             100 * tab_coinfect$both / (tab_coinfect$wt +
               tab_coinfect$syn + tab_coinfect$both),
             100 * tab_coinfect$coinfection_freq),
           hjust=1,
           vjust=0.4,
           fontface="italic")

saveShowPlot(p_coinfect, width=2.9, height=2.2)
```

An interesting feature of this analysis is that even though the results above indicate that less than a quarter of cells are called *infected*, more than half are coinfected.
This indicates that there is some enrichment for coinfected cells beyond what would be expected under a Poisson process.
This could be because infection is not Poisson, because coinfection increases our ability to detect coinfection (such as by increasing the mRNA from an infecting virus with a defective polymerase), or because the interferon sorting enriched for coinfected cells among the infected cells.

#### Annotate interferon status¶

We now want to identify the cells that are interferon (IFN) positive.

##### Calculate IFN and ISG expression¶

A549 cells induce both type I and type III IFN in response to viral infection.
In addition, autocrine and paracrine signaling induce expression of various IFN-stimulated genes (ISGs).

We get all type I and type III interferons by their names, and select a few ISGs that we know have high expression.
Below we print the names of the genes these IFNs and ISGs:

In [44]:

```
isg_genes <- c("IFIT1", "ISG15", "CCL5", "MX1")

ifn_genes <- fData(cells) %>% 
  select(gene_short_name) %>%
  filter(str_detect(gene_short_name, "IFN[ABL]") | gene_short_name %in% isg_genes,
         !str_detect(gene_short_name, "IFN.R")) %>%
  mutate(genetype=ifelse(gene_short_name %in% isg_genes, "ISG",
                         ifelse(str_detect(gene_short_name, "IFNL"),
                         "type_III_IFN", "type_I_IFN"))) %>%
  arrange(genetype, gene_short_name)

ifn_genes %>%
  group_by(genetype) %>%
  mutate(genes=paste0(gene_short_name, collapse=", ")) %>%
  select(genetype, genes) %>%
  filter(row_number() == 1)
```

| genetype | genes |
| --- | --- |
| ISG | CCL5, IFIT1, ISG15, MX1 |
| type\_I\_IFN | IFNA1, IFNA10, IFNA13, IFNA14, IFNA16, IFNA17, IFNA2, IFNA21, IFNA4, IFNA5, IFNA6, IFNA7, IFNA8, IFNB1 |
| type\_III\_IFN | IFNL1, IFNL2, IFNL3 |

Now for each cell, we calculate:

1. The number mRNA from each of these categories of genes.
2. The fraction and log10 fraction of all cellular mRNA from each of these categories of genes.

For the fraction calculations, we add a pseudocount to the number of IFN or ISG mRNAs.
This fraction is therefore an **upper bound** on the amount of each type of mRNA, and also allows us to avoid taking the logarithm of zero.

In [45]:

```
pseudocount <- 1

for (gtype in ifn_genes$genetype %>% unique) {
  genes <- ifn_genes %>% filter(genetype == gtype) %$% gene_short_name
  pData(cells)[[gtype]] <- Matrix::colSums(exprs(
      cells[row.names(subset(fData(cells), gene_short_name %in% genes)), ]))
  pData(cells)[[paste0(gtype, "_frac")]] <- (pData(cells)[[gtype]] + 
      pseudocount) / pData(cells)$cell_mRNAs
  pData(cells)[[paste0("log_", gtype, "_frac")]] <- log10(pData(cells)[[paste0(gtype, "_frac")]])
}
```

##### Correlation between type I and III IFN¶

We plot the correlation between relative expression measure (fraction of cellular mRNA) for type I and type III IFN:

In [46]:

```
p_ifn_genes_corr <- ggplot(
    pData(cells) %>%
      mutate(infected=ifelse(infected, "infected", "not infected")),
    aes(x=type_III_IFN_frac, y=type_I_IFN_frac, color=infected)) +
  geom_point(alpha=0.5, size=2) +
  stat_cor(color='black', fontface='italic') +
  scale_color_manual(values=rev(rev(cbPalette[3:4]))) +
  facet_wrap(~infected, nrow=1) +
  theme(legend.position="none", strip.text=element_text(size=14)) +
  xlab("frac mRNA from IFN III") +
  ylab("frac mRNA from IFN I")

saveShowPlot(p_ifn_genes_corr, width=4.7, height=2.5, isfig=TRUE)
```

The plot above shows that the expression of type I and type III IFNs is highly correlated.
Also, most cells expressing either of these IFNs are clearly flu infected (the others must either not be infected or just have such low amounts of flu that we failed to call them as infected).

Because type I and type III IFN are so correlated, we will define a new variable which is just the sum of type I and type III IFN expressions, and use this to call IFN status.
Note that this variable *mostly* just reflects type III IFN levels since these are $\approx$10-fold higher than type I levels.
We also compute the fraction and log10 fraction of mRNA from IFN, again adding a pseudocount to avoid logs of zero:

In [47]:

```
pData(cells)$IFN <- pData(cells)$type_I_IFN + 
  pData(cells)$type_III_IFN

pData(cells)$IFN_frac <- (pData(cells)$IFN + 
  pseudocount) / pData(cells)$cell_mRNAs

pData(cells)$log_IFN_frac <- log10(pData(cells)$IFN_frac)
```

##### Call IFN+ and plot IFN expression¶

Based on the plots above, we will call IFN status based just on the total fraction of mRNA from IFN (type I and type III combined).
We don't use the ISGs because we are interested in cells that directly activate IFN due to viral infection, not cells that express ISGs possibly due to paracrine signaling.

We will classify cells into three IFN status groups:

- *IFN-*: cells are called interferon negative if the fraction of their mRNA from IFN is $\le$ that expected in the lowest-mRNA cell if it had 2 IFN mRNAs.
- *IFN+*: cells are called interferon positive if the fraction of their mRNA from IFN is $\ge$ that expected in the lowest-mRNA cell if it had 5 IFN mRNAs.
- *IFNambiguous*: cells that fall between *IFN-* and *IFN+* are called ambiguous because it is harder to determine if they are truly IFN positive or just picked up IFN mRNA from environmental mRNA.

In [48]:

```
pData(cells)$log_IFNneg_frac_threshold <- pData(cells) %>%
  mutate(threshold=log10(2 / min(cell_mRNAs))) %$%
  threshold

pData(cells)$log_IFNpos_frac_threshold <- pData(cells) %>%
  mutate(threshold=log10(5 / min(cell_mRNAs))) %$%
  threshold

cat(sprintf(paste(
    "Calling IFN- if <=%.2e%% of mRNA from IFN",
    "and IFN+ if >=%.2e%% from IFN."),
  10**(pData(cells) %>% head(n=1) %$% log_IFNneg_frac_threshold),
  10**(pData(cells) %>% head(n=1) %$% log_IFNpos_frac_threshold)))

pData(cells)$IFN_state <- pData(cells) %>%
  mutate(IFN_state=ifelse(log_IFN_frac >= log_IFNpos_frac_threshold, "IFN+",
                   ifelse(log_IFN_frac <= log_IFNneg_frac_threshold, "IFN-",
                   "IFNambiguous"))) %$%
  IFN_state
```

```
Calling IFN- if <=1.69e-04% of mRNA from IFN and IFN+ if >=4.21e-04% from IFN.
```

Below we plot cells in these classifications.
It is immediately obvious that there are many more IFN+ cells among the infected ones, which is as we expect if infection is what is inducing IFN.

In [49]:

```
# histograms of IFN expression
p_ifn_dist <- ggplot(
    pData(cells) %>% 
      mutate(
        infected=ifelse(infected, "infected", "not infected"),
        fill=ifelse(IFN_state %in% c("IFN+", "IFNambiguous"), IFN_state,
             ifelse(infected == "infected", "IFN-infected", "IFN-uninfected"))
        ),
    aes(log_IFN_frac, fill=fill)) + 
  geom_histogram(bins=37) +
  ylab("number of cells") +
  scale_fill_manual(values=c(cbPalette[3], cbPalette[4], 
                             cbPalette[2], cbPalette[1])) +
  geom_vline(aes(xintercept=log_IFNneg_frac_threshold), 
                 color=cbbPalette[[8]], linetype='dashed') +
  geom_vline(aes(xintercept=log_IFNpos_frac_threshold), 
                 color=cbbPalette[[8]], linetype='dashed') +
  facet_wrap(~ infected, scales="free_y", ncol=1) +
  theme(legend.position="none", strip.text=element_text(size=14)) +
  scale_x_continuous(name="frac mRNA from IFN",
    labels=function (x) fancy_scientific(10**x))

# numbers to annotate plot
n_infected <- pData(cells) %>% filter(infected) %>% nrow
n_uninfected <- pData(cells) %>% filter(! infected) %>% nrow
n_infected_IFN <- pData(cells) %>% 
  filter(infected & IFN_state == "IFN+") %>% nrow
n_uninfected_IFN <- pData(cells) %>% 
  filter(! infected & IFN_state == "IFN+") %>% nrow 

# annotate plot
p_ifn_dist <- ggdraw() +
  draw_plot(p_ifn_dist, 0, 0, 1, 1) +
  draw_label(
    paste0(n_infected_IFN, " of ", n_infected, "\nIFN+"),
    x=0.95, y=0.85, size=12, hjust=1, fontface='italic') +
  draw_label(
    paste0(n_uninfected_IFN, " of ", n_uninfected, "\nIFN+"),
    x=0.95, y=0.43, size=12, hjust=1, fontface='italic')
                     
saveShowPlot(p_ifn_dist, 3, 3.5)

# print statistics
pData(cells) %>%
  group_by(infected, IFN_state) %>%
  summarize(cells=n())
```

| infected | IFN\_state | cells |
| --- | --- | --- |
| FALSE | IFN- | 1175 |
| FALSE | IFN+ | 15 |
| FALSE | IFNambiguous | 10 |
| TRUE | IFN- | 217 |
| TRUE | IFN+ | 60 |
| TRUE | IFNambiguous | 13 |

The plots above show that (as we expect), there is a much higher fraction of IFN+ cells among the clearly virally infected ones.

What about the rare handful of IFN+ cells that we did not call as infected?
Maybe our experiment sorted out cells that are spontaneously activating IFN or activating it in response to some damage-associated molecular pattern (DAMP) from the infected cells--or maybe some of the cells we call as un-infected are fact infected at very low levels that induces IFN, but just didn't pass our filters for being clearly infected.

##### Plot ISG expression¶

Now we make a plot similar to the one above but for expression of ISGs rather than IFN.
Note that by "ISG" we mean the total mRNA counts for the four prototype ISGs defined above.

Since our downstream analyses don't actually do anything with ISG expression, we spend less time worrying about how this is called.
We just make a heuristic cutoff of saying a cell is ISG+ if it has more than 0.1% of its mRNA from the four ISG genes.

We then plot the distribution of ISG expression for both infected and uninfected cells.
We see that for both infected and uninfected there are more ISG+ than IFN+ cells, but the increase is much larger (10-fold) for uninfected cells than it is for infected ones (2-fold).
In both cases, presumably paracrine signaling is leading to ISG expression in the absence of IFN.

In [50]:

```
# heuristic cutoff for calling ISG+
ISG_cutoff <- 1e-3

# classify cells based on heuristic threshold
pData(cells)$ISG_state <- ifelse(pData(cells)$ISG_frac > ISG_cutoff, "ISG+", "ISG-")

# make histogram
p_isg_dist <- ggplot(
    pData(cells) %>% 
      mutate(
        infected=ifelse(infected, "infected", "not infected"),
        fill=ifelse(ISG_state == "ISG+", "ISG+",
             ifelse(infected == "infected", "ISG-infected", "ISG-uninfected"))
        ),
    aes(log_ISG_frac, fill=fill)) + 
  geom_histogram(bins=25) +
  geom_vline(aes(xintercept=log10(ISG_cutoff)), 
                 color=cbbPalette[[8]], linetype='dashed') +
  ylab("number of cells") +
  scale_fill_manual(values=c(cbPalette[3], cbPalette[4], cbPalette[7])) +
  facet_wrap(~ infected, scales="free_y", ncol=1) +
  theme(legend.position="none", strip.text=element_text(size=14)) +
  scale_x_continuous(name="frac mRNA from ISG",
    labels=function (x) fancy_scientific(10**x))
                     
# numbers to annotate plot
n_infected_ISG <- pData(cells) %>% 
  filter(infected & ISG_frac > ISG_cutoff) %>% nrow
n_uninfected_ISG <- pData(cells) %>% 
  filter(! infected & ISG_frac > ISG_cutoff) %>% nrow 
                     
# annotate plot
p_isg_dist <- ggdraw() +
  draw_plot(p_isg_dist, 0, 0, 1, 1) +
  draw_label(
    paste0(n_infected_ISG, " of ", n_infected, "\nISG+"),
    x=0.95, y=0.85, size=12, hjust=1, fontface='italic') +
  draw_label(
    paste0(n_uninfected_ISG, " of ", n_uninfected, "\nISG+"),
    x=0.95, y=0.43, size=12, hjust=1, fontface='italic')
                     
saveShowPlot(p_isg_dist, 3, 3.5)
```

We next plot the correlation between ISG and IFN expression:

In [51]:

```
p_isg_corr <- ggplot(
    pData(cells) %>%
      mutate(infected=ifelse(infected, "infected", "not infected"),
             fill=ifelse(ISG_state == "ISG+", "ISG+",
                  ifelse(infected == "infected", "ISG-infected",
                         "ISG-uninfected"))),
    aes(x=log_ISG_frac, y=log_IFN_frac, color=fill)) +
  geom_point(alpha=0.5, size=2) +
  stat_cor(color='black', fontface='italic', label.sep='\n') +
  scale_color_manual(values=c(cbPalette[3], cbPalette[4], cbPalette[7])) +
  facet_wrap(~infected, ncol=1) +
  theme(legend.position="none", strip.text=element_text(size=14)) +
  scale_x_continuous(name="frac mRNA from ISG",
    labels=function (x) fancy_scientific(10**x)) +
  scale_y_continuous(name="frac mRNA from IFN",
    labels=function (y) fancy_scientific(10**y))

saveShowPlot(p_isg_corr, width=2.7, height=4.5)
```

This plot confirms that ISG expression is reasonably correlated with IFN expression in infected cells (although there are some high ISG cells without IFN), but that it is not well correlated in uninfected cells.
Therefore, most of the uninfected cells expression ISGs are probably doing so due to paracrine signaling.

We merge the two plots above into a single figure for the paper:

In [52]:

```
p_isg <- plot_grid(p_isg_dist, p_isg_corr,
          nrow=1, align='h', scale=0.9, rel_widths=c(1.25, 1),
          labels=c("A", "B"), label_size=18, hjust=-1)

saveShowPlot(p_isg, width=6.3, height=4.5, isfig=TRUE)
```

#### Unsupervised tSNE clustering¶

Now we are going to use monocle's capabilities for clustering the cells.
The goal is simply to visualize what genes explain the structure of the data.

##### Estimate size factors and dispersions¶

First, we estimate the size factors and dispersions.
Note that for host differential gene expression of factors associated with influenza burden, there is some question of whether this should be done on cellular mRNAs only or on all mRNAs, since the viral mRNAs displace cellular mRNAs and so perhaps should not be considered for size factors.
But for clustering, we want to use all the genes including the viral ones, so we estimate size factors on everything.
We also annotate the number of cells in which each gene is detectably expressed.
All of these operations are directly out of the monocle vignette.

In [53]:

```
cells <- estimateSizeFactors(cells)
cells <- estimateDispersions(cells)
cells <- detectGenes(cells, min_expr=0.1)
```

```
Removing 115 outliers
```

We also make sure that the expression values are lognormally distributed as recommended by the monocle vignette.
The plot below shows that yes, they are close.

In [54]:

```
p_expr_lognorm <- ggplot(
  exprs(cells) %>%
      log %>%
      (Matrix::t) %>% 
      scale %>% 
      (Matrix::t) %>%
      melt,
    aes(value)) +
  geom_density(color=cbbPalette[[2]]) +
  stat_function(fun=dnorm, size=0.5, color=cbbPalette[[3]]) +
  xlab("standardized log(mRNAs)")

saveShowPlot(p_expr_lognorm, 3, 2)
```

##### Unsupervised tSNE clustering¶

We now select reasonalby highly expressed genes to use for clustering, plot their expression and dispersion, and plot the variance explained by each component.
This is all paralleling the monocle vignette.

In [55]:

```
unsup_clustering_genes <- dispersionTable(cells) %>% subset(mean_expression > 0.5)
cells <- setOrderingFilter(cells, unsup_clustering_genes$gene_id)

p_ordering_genes <- plot_ordering_genes(cells)
saveShowPlot(p_ordering_genes, 3.5, 3)

p_variance_explained <- plot_pc_variance_explained(cells, return_all=FALSE)
saveShowPlot(p_variance_explained, 3.5, 3)
```

Now we do the tSNE clustering.

In [56]:

```
cells <- reduceDimension(cells, max_components=2, num_dim=6, reduction_method='tSNE')
cells <- clusterCells(cells)
```

```
Distance cutoff calculated to 3.270851
```

And now we plot the cells color by the amount of flu mRNA, IFN expression, and expression of our selected ISGs.
We see that flu expression and ISG expression explain a large amount of the structure in the data, as there is a distinct group of flu positive cells, and then a distinct group of ISG-expressing cells which partially overlaps with flu positive cells.

In [57]:

```
pData(cells)$log_frac_mRNA_from_flu <- log10(pData(cells)$frac_mRNA_from_flu)

tsne_plotlist <- map(
  c("log_frac_mRNA_from_flu", "log_IFN_frac", "log_ISG_frac",
    "frac_mRNA_from_flu", "IFN_frac", "ISG_frac"),
  function (attr) {
    plot_cell_clusters(cells, color=attr) +
      scale_color_gradient(
        low=cbPalette[1],
        high=ifelse(str_detect(attr, "flu"), cbPalette[3],
             ifelse(str_detect(attr, "IFN"), cbPalette[2], cbPalette[7])),
        name=gsub("frac mRNA from flu", "flu frac",
             gsub("_", " ", 
             gsub("log", "log10", attr)))) +
      guides(color=guide_colorbar(title.position="top", title.hjust=0.5,
        barwidth=10, barheight=0.8), alpha=NULL)
  }
  )

ncol <- 3
p_tsne <- plot_grid(plotlist=tsne_plotlist, ncol=ncol, scale=0.95)

saveShowPlot(p_tsne,
             width=3.5 * min(length(tsne_plotlist), ncol), 
             height=4.2 * ceiling(length(tsne_plotlist) / ncol),
             isfig=TRUE)
```

#### Get viral genotypes from PacBio¶

The PacBio sequencing allows us to analyze viral genotypes, using the cell annotations added by the pacbio\_analysis.ipynb notebook.
This is different than the 10X data above, which simply counts the 3' ends of transcripts.

We obtain viral genotypes only for infected cells, so get these:

In [58]:

```
infected_cells <- cells[, pData(cells)$infected]
```

##### Look at NS1 / NS2 and M1 / M2 isoforms¶

One piece of information that the PacBio sequencing provides that the 10X does not is the identity of different isoforms of the two alternatively spliced genes (M1 / M2 and NS1 / NS2).

Note that the PacBio sequencing counts are expected to be much less good at quantitating levels of different transcripts due to the many rounds of PCR that are performed to get enough material for library prep.

Nonetheless, we can check if the levels M1 / M2 and NS1 / NS2 (or any other PacBio counted gene) is associated with IFN state, again only looking at cells that we can unambiguously call as IFN+ or IFN-.
We do that below.
We repeat the finding from 10X that there is a (non-significant) association between low NS1 and IFN+, and of high PB2 and IFN+.
But there is nothing obvious with NS2 and M2, so it does **not** appear that the abundance of these different isoforms is a major factor:

In [59]:

```
# the isoforms of all flu genes called by PacBio
isoforms <- c("fluPB2", "fluPB1", "fluPA", "fluHA", "fluNP",
              "fluNA", "fluM1", "fluM2", "fluNS1", "fluNS2")

# data frame giving fraction of PacBio from each isoform
isoform_counts <- pData(infected_cells) %>%
  filter(IFN_state %in% c("IFN+", "IFN-")) %>%
  gather(gene_barcodes, n_PacBio,
         lapply(isoforms, function (i) c(paste0(i, "_wt_n_PacBio_seqs"),
                                         paste0(i, "_syn_n_PacBio_seqs"))) 
         %>% unlist) %>%
  separate(gene_barcodes, "isoform", sep="_", extra="drop") %>%
  transform(isoform=factor(isoform, isoforms)) %>%
  group_by(CellBarcode, isoform, IFN_state) %>%
  summarize(n_PacBio_isoform=sum(n_PacBio)) %>%
  group_by(CellBarcode) %>%
  mutate(n_PacBio_cell=sum(n_PacBio_isoform),
         frac_PacBio=n_PacBio_isoform / n_PacBio_cell) %>%
  ungroup

p_isoformfrac_IFN <- ggplot(isoform_counts, aes(IFN_state, frac_PacBio)) +
  geom_boxplot(outlier.color=NA) + 
  geom_jitter(width=0.25, alpha=0.2, size=1) +
  facet_wrap(~isoform, nrow=1) +
  stat_compare_means(method="wilcox.test",
    aes(label = paste0("P = ", ..p.format..)), vjust=0.5) 

saveShowPlot(p_isoformfrac_IFN, 11, 3)
```

We also take a detailed look at the relationship just between the two alternate isoform pairs (M1 / M2 and NS1 / NS2) in the IFN+ and IFN- cells.
That plot is below.
Again, it indicates that altered ratios of these two isoforms does **not** appear to explain IFN induction in our experiments, and the relationship looks similarly linear between the two isoforms for both IFN+ and IFN- cells:

In [60]:

```
isoform_scatter_plots <- lapply(
  list(c("fluM1", "fluM2"), c("fluNS1", "fluNS2")),
  function (gene_isoforms)
    ggplot(
      isoform_counts %>%
        select(-frac_PacBio) %>%
        filter(isoform %in% gene_isoforms) %>%
        spread(key=isoform, value=n_PacBio_isoform),
      aes_string(gene_isoforms[[1]], gene_isoforms[[2]], color='IFN_state')) +
    geom_point(alpha=0.4) +
    geom_smooth(method='lm', se=FALSE) +
    scale_color_manual(values=rev(cbbPalette[2:3]))
  )

p_isoform_scatter <- plot_grid(plotlist=isoform_scatter_plots, nrow=1)
    
saveShowPlot(p_isoform_scatter, 7, 2.5)
```

Given these results, we don't use the PacBio to try to quantify any type of gene levels any further, as beyond M1 / M2 and NS1 / NS2, the 10X data should be vastly better at quantifying gene expression levels.

##### For which cells can we call viral genotypes?¶

We now want to determine for which cells we can call the full viral genotype.
By calling the genotype, we mean using the PacBio sequencing to identify the mutations present in all viral genes that the 10X sequencing says are present in that cell.

###### Annotate presence of barcoded viral genes¶

First, we need to determine from the 10X sequencing for which viral genes each cell should have.
We have already narrowed ourselves down to infected cells, and then called each these cells by whether they are co-infected both viral barcodes, and whether they have each flu gene.
However, we haven't yet called whether each cell has each flu gene from each viral barcode.

We do this below.
We say that a cell should have a particular gene and viral barcode if:

- The call has been called as being infected with that viral barcode variant.
- The cell has been called as having that particular viral gene.
- For that gene, the viral barcode variant in question is >= 25% of the total observed counts of that gene **or** the P-value that the number of annotated reads for that gene exceed background expected under the Poisson model defined above is < 0.05 (the cutoff used earlier in this notebook to define gene presence / absence).

In [61]:

```
# the two viral barcodes
viral_barcodes <- c("wt", "syn")

# has gene barcode if coinfected, has gene, and gene barcode frac > this
has_barcode_min_frac <- 0.25
   
# get total number mRNAs 
n_mRNA <- pData(infected_cells)$total_mRNAs

# annotate cells by whether they have barcoded gene
for (gene in flugenes) {
    
  # get background frac flu for this gene
  gene_background_frac <- thresholds$frac[thresholds$gene == gene]
    
  # get number of barcode annotated reads for gene
  n_gene <- pData(infected_cells)[[paste('annotated', gene, 'wt', sep='_')]] +
            pData(infected_cells)[[paste('annotated', gene, 'syn', sep='_')]]
            
  # fraction annotated
  frac_annotated <- pData(infected_cells)$n_annotated_flu / pData(infected_cells)$flu_mRNAs 
    
  for (bc in viral_barcodes) {
    # get number reads for barcoded gene
    n_gene_bc <- pData(infected_cells)[[paste('annotated', gene, bc, sep='_')]]    
    # call whether barcoded gene present
    pData(infected_cells)[paste('has', gene, bc, sep='_')] <- (
      pData(infected_cells)[paste('infected', bc, sep='_')] &
      pData(infected_cells)[paste('has', gene, sep='_')] &
      (((n_gene_bc / n_gene) > has_barcode_min_frac) |
        (map2(n_gene_bc, gene_background_frac * n_mRNA * frac_annotated, poisson_P) 
              < p_has_gene_cutoff))
      ) %>% as.vector
  }
}
```

Now we make a tidy data frame (called *genotypes*) with the gene presence and mutation data.
We add the following columns for each cell:

- *flu\_gene*: name of that viral gene
- *viral\_barcode*: viral barcode variant
- *barcoded\_gene\_present*: whether that barcoded viral gene is present according to 10X sequencing
- *mutation\_type*: the type of mutation (substitutions, insertions, deletions)
- *mutations*: string giving mutations of this type, or *NA* if not data available.

We do this only for cells for which we can unambiguously call the IFN state, as we care about analyzing how mutations associate with IFN state.

In [62]:

```
# types of mutations
mutation_types <- c("substitutions", "deletions", "insertions")

# vector with all combinations of flu genes and viral barcodes
genes_barcodes <- lapply(flugenes, function (g) lapply(viral_barcodes,
  function (bc) paste(g, bc, sep='_'))) %>% unlist

# vector with all combinations of flu genes, viral barcodes, mutations
genes_barcodes_mutations <- lapply(genes_barcodes, function(g_bc) lapply(
  mutation_types, function(m) paste(g_bc, m, sep='_'))) %>% unlist

# tidy data frame with genotype information
genotypes <- pData(infected_cells) %>%
  # make tidy for barcoded genes expected by 10X
  gather(key="gene_barcode", value="barcoded_gene_present",
    lapply(genes_barcodes, function (x) paste0('has_', x)) %>% unlist) %>%
  separate(gene_barcode, c("dummy", "flu_gene", "viral_barcode")) %>%
  select(-dummy) %>%
  # make tidy for PacBio data for barcoded genes
  gather(key="gene_barcode_mutation", value="mutations",
    genes_barcodes_mutations) %>%
  separate(gene_barcode_mutation, c("flu_gene2", "viral_barcode2", "mutation_type")) %>%
  filter(flu_gene == flu_gene2 & viral_barcode == viral_barcode2) %>%
  select(-flu_gene2, -viral_barcode2) %>%
  # make flu_gene a factor
  transform(flu_gene=factor(flu_gene, flugenes)) %>%
  # add nice label for coinfection state
  mutate(coinfection_state=factor(
    ifelse(coinfected, "dual-barcode coinfection", "single viral barcode"),
    c("single viral barcode", "dual-barcode coinfection")))
           
# make sure we have expected number of rows
stopifnot(nrow(genotypes) == (nrow(pData(infected_cells)) * 
  length(flugenes) * length(viral_barcodes) * length(mutation_types)))
           
# only cells with unambiguous IFN_state
genotypes <- genotypes %>% filter(IFN_state %in% c("IFN+", "IFN-"))
```

###### Identify genes missing PacBio sequences¶

Now we want to identify gene / barcode combinations that based on the 10X data we called as being present, but for which we do not have a PacBio consensus.
First, annotate these with the *missing\_PacBio* column in the data frame:

In [63]:

```
genotypes <- genotypes %>%
  mutate(missing_PacBio=barcoded_gene_present & is.na(mutations)) %>%
  # add number of missing gene / barcode combinations per cell
  group_by(CellBarcode) %>%
  mutate(n_missing_gene_barcode=sum(missing_PacBio) / length(mutation_types)) %>%
  ungroup
```

Now plot the distribution of the number of gene / barcode combinations that are missing PacBio data.
Obviously, our goal is to get as many cells as possible to have zero expected genes missing:

In [64]:

```
p_n_missing <- ggplot(
    genotypes %>% group_by(CellBarcode) %>% summarize_all(funs(dplyr::first)),
    aes(n_missing_gene_barcode)) +
  geom_bar() +
  scale_x_continuous(breaks=pretty_breaks()) +
  ylab("number of cells") +
  xlab("number of PacBio genes / barcodes missing") +
  facet_grid(IFN_state ~ coinfection_state, scales='free_y')

saveShowPlot(p_n_missing, width=5.5, height=4)
```

We also look to see **which** flu genes tend to be missing.
Below we count up the number of times each flu genes is missing **among the cells that are missing just one or two genes**.
We see that the genes that are missing most often are HA and NA.
Therefore, if we were to do more PacBio sequencing, those would be the genes that it would be most productive to target more.

In [65]:

```
p_missing_by_gene <- ggplot(
    genotypes %>% 
      filter(n_missing_gene_barcode <= 2) %>%
      group_by(flu_gene, viral_barcode, IFN_state, coinfection_state) %>%
      summarize(n_missing=sum(missing_PacBio / length(mutation_types))),
    aes(flu_gene, n_missing, fill=viral_barcode)) +
  geom_bar(position=position_dodge(), stat='identity') +
  scale_fill_manual(values=cbPalette[2:3]) +
  facet_grid(IFN_state ~ coinfection_state, scales='free_y') +
  theme(axis.text.x=element_text(angle=90, hjust=1, vjust=0.5))

saveShowPlot(p_missing_by_gene, width=7, height=3.5)
```

###### Identify cells with complete genotypes¶

Now we annotate as *complete* the cells for which we can use the PacBio to call the complete viral genotype.
By complete, we don't mean that all viral genes are present in the cell, but rather that we can use the PacBio to call the genotype of all genes that the 10X sequencing says is present.

We then plot the number of such cells below.
As can be seen below, we are worse at calling these genotypes in dual-barcode co-infections.

In [66]:

```
genotypes <- genotypes %>%
  mutate(complete=(n_missing_gene_barcode == 0),
         complete_state=ifelse(complete,
           "complete", "incomplete"))

n_complete_genotypes <- genotypes %>%
  group_by(IFN_state, coinfection_state, complete_state) %>%
  summarize(n_cells=length(unique(CellBarcode))) 

p_n_complete_genotype <- ggplot(
    genotypes %>%
      group_by(IFN_state, coinfection_state, complete_state) %>%
      summarize(n_cells=length(unique(CellBarcode))),
    aes(complete_state, n_cells, fill=IFN_state, label=n_cells)) +
  geom_bar(stat='identity', position=position_dodge()) +
  geom_text(position=position_dodge(width=1), vjust=-0.3) +
  facet_wrap(~ coinfection_state) +
  scale_fill_manual(values=rev(cbPalette[2:3]), name=NULL) +
  theme(
    axis.ticks.y=element_blank(),
    axis.text.y=element_blank(),
    strip.text=element_text(size=14, margin=margin(b=3))) +
  scale_y_continuous(name="number of cells", 
    limits=c(0, 1.15 * (n_complete_genotypes$n_cells %>% max))) +
  xlab("viral genes in cell completely sequenced?")

saveShowPlot(p_n_complete_genotype, 6.1, 2.5)
```

In [67]:

```
p_frac_flu_complete_genotype <- ggplot(
    genotypes %>% 
      group_by(CellBarcode, complete_state, 
               frac_mRNA_from_flu, IFN_state) %>%
      summarize_all(funs(dplyr::first)),
    aes(complete_state, frac_mRNA_from_flu)) +
  geom_boxplot(aes(color=IFN_state), position=position_dodge(),
    outlier.size=1, outlier.alpha=0.4) +
  geom_boxplot(aes(fill=IFN_state), position=position_dodge(),
    outlier.color=NA) +
  facet_wrap(~ coinfection_state) +
  scale_fill_manual(values=rev(cbPalette[2:3]), name=NULL) +
  scale_color_manual(values=rev(cbPalette[2:3]), guide=FALSE) +
  theme(strip.text=element_text(size=14, margin=margin(b=3))) +
  scale_y_continuous(name="frac mRNA from flu") + 
  xlab("viral genes in cell completely sequenced?")

saveShowPlot(p_frac_flu_complete_genotype, 6.3, 2.35)
```

Merge the two plots above into a single plot for a figure:

In [68]:

```
p_cells_complete <- plot_grid(
  p_n_complete_genotype + 
    theme(axis.text.x=element_blank(),
          axis.ticks.x=element_blank(),
          axis.title.x=element_blank()),
  p_frac_flu_complete_genotype +
    theme(strip.background=element_blank(),
          strip.text.x=element_blank()),
  ncol=1, align='v', rel_heights=c(1, 1.25),
  labels=c("A", "B"), label_size=18, vjust=1)

saveShowPlot(p_cells_complete, 6.3, 4.75, isfig=TRUE)
```

##### Mutation types in PacBio data¶

Now we examine which types of mutations are present as annotated in pacbio\_analysis.ipynb.
For all gene / barcode / mutation combinations, we tally up the number of sequences with mutations, both among cells that do and do not have complete sequencing of the viral genomes.
We see below that substititutions are by far the most common type of mutation, then deletions, then insertions.

In [69]:

```
genotypes %>% 
  filter(barcoded_gene_present & ! is.na(mutations)) %>%
  mutate(has_mutation=mutations != "WT") %>%
  group_by(complete, mutation_type) %>%
  summarize(wildtype=sum(! has_mutation), mutated=sum(has_mutation)) %>%
  mutate(frac=mutated / wildtype) %>%
  arrange(-complete, -mutated)
```

| complete | mutation\_type | wildtype | mutated | frac |
| --- | --- | --- | --- | --- |
| TRUE | substitutions | 1280 | 104 | 0.081250000 |
| TRUE | deletions | 1372 | 12 | 0.008746356 |
| TRUE | insertions | 1377 | 7 | 0.005083515 |
| FALSE | substitutions | 708 | 73 | 0.103107345 |
| FALSE | deletions | 766 | 15 | 0.019582245 |
| FALSE | insertions | 775 | 6 | 0.007741935 |

We create a *gene\_state* variable that indicates if each barcoded gene variant is wildtype, absent, mutated, or has an unknown sequence:

In [70]:

```
genotypes <- genotypes %>%
  mutate(gene_state=
    ifelse(! barcoded_gene_present, "absent",
    ifelse(missing_PacBio, "unknown",
    ifelse(mutations == "WT", "wildtype", "mutated")))) %>%
  transform(gene_state=factor(gene_state,
    c("wildtype", "mutated", "absent", "unknown")))
```

#### Plot and print complete genotypes¶

We now want to plot and print the complete genotypes of all cells for which we could call all genes that are present.

##### Prepare to plot genotypes¶

This plot is fairly complex, so we have to do a lot of data cleaning to make it.

First, we number the cells with complete genomes (to provide succinct labels) in the *genotypes\_complete* data frame, and arrange by the amount of viral mRNA:

In [71]:

```
genotypes_complete <- genotypes %>%
  filter(complete) %>%
  arrange(-frac_mRNA_from_flu) %>%
  separate(CellBarcode, "CellBarcode_only", extra="drop", remove=FALSE) %>%
  mutate(cell=paste("cell", match(CellBarcode_only, unique(CellBarcode_only)))) %>%
  transform(cell=factor(cell, rev(unique(cell))))
```

Annotate each cell / gene by the number of viral barcodes for it (0 if gene not present, 1 if one barcoded variant, 2 if both barcoded variants):

In [72]:

```
n_viral_variants_df <- genotypes_complete %>%
  filter(complete) %>%
  select(cell, flu_gene, viral_barcode, barcoded_gene_present) %>%
  distinct %>%
  spread(viral_barcode, barcoded_gene_present) %>%
  mutate(n_viral_variants=factor(as.integer(wt) + as.integer(syn))) 

genotypes_complete <- merge(genotypes_complete, 
                           n_viral_variants_df,
                           by=c('cell', 'flu_gene'))
```

Create a dataframe (*genotypes\_by\_gene*) that aggregates the data for viral barcodes for each gene and cell.
In this dataframe, we no longer track the wildtype and synonymous barcodes separately, but rather simply track what is the genotype of all genes for that virus in that cell.

In [73]:

```
# get genotypes by gene (aggregating mutation types and viral barcodes)
genotypes_by_gene <- genotypes_complete %>%
  dplyr::rename(cell_barcode=CellBarcode_only) %>%
  # select just columns we care about
  select(cell, cell_barcode, frac_mRNA_from_flu, IFN_frac, IFN_state,
    coinfection_state, flu_gene, viral_barcode, mutation_type, mutations,
    barcoded_gene_present, n_viral_variants) %>%
  # annotate mutations into string
  mutate(mutations=recode(mutations, "WT" = "")) %>%
  spread(mutation_type, mutations) %>%
  mutate(mutation_str=ifelse(! barcoded_gene_present,
    paste(flu_gene, "missing", sep='-'),
    ifelse(is.na(substitutions) | is.na(insertions) | is.na(deletions), "unknown",
    ifelse(substitutions == "" & insertions == "" & deletions == "",
    paste(flu_gene, "WT", sep='-'),
    paste(substitutions, insertions, deletions))))) %>%
  select(-barcoded_gene_present, -substitutions, -insertions, -deletions) %>%
  verify(mutation_str != "unknown") %>%
  # aggregate both viral barcodes and label mutations with long and short str
  spread(viral_barcode, mutation_str) %>%
  mutate(mutation_str_long=ifelse(syn == wt, wt,
    ifelse(grepl("missing", syn), wt, ifelse(grepl("missing", wt), syn,
    paste(wt, syn, sep=" / "))))) %>%
  mutate(mutation_str_long=ifelse(mutation_str_long == paste(flu_gene, "WT", sep='-'),
    "", mutation_str_long)) %>%
  mutate(mutation_str_short=gsub("_\\(\\d+/\\d+\\)", "", mutation_str_long))
```

In preparation for plotting, create a data frame (*flu\_seq\_coords*) that gives the coordinates of all sites in the flu genome if we treat them as one continuous chromosome demarcated by gene boundaries.
We do this reading in the vRNA lengths from ./data/flu\_sequences/flu-wsn.fasta, and arrange the genes from longest to shortest as this is the convention for numbering flu gene segments:

In [74]:

```
# read in flu genes
flu_seqs <- readDNAStringSet("data/flu_sequences/flu-wsn.fasta") 

# get start / end coords of genes
flu_seq_coords <- data.frame(flu_gene=names(flu_seqs), gene_length=width(flu_seqs)) %>%
  separate(flu_gene, "flu_gene") %>%
  transform(flu_gene=factor(flu_gene, flugenes)) %>%
  mutate(gene_end=cumsum(gene_length), gene_start=gene_end - gene_length + 1,
    gene_center=(gene_start + gene_end) / 2)

flu_seq_coords
```

| flu\_gene | gene\_length | gene\_end | gene\_start | gene\_center |
| --- | --- | --- | --- | --- |
| fluPB2 | 2341 | 2341 | 1 | 1171.0 |
| fluPB1 | 2341 | 4682 | 2342 | 3512.0 |
| fluPA | 2233 | 6915 | 4683 | 5799.0 |
| fluHA | 1775 | 8690 | 6916 | 7803.0 |
| fluNP | 1565 | 10255 | 8691 | 9473.0 |
| fluNA | 1409 | 11664 | 10256 | 10960.0 |
| fluM | 1027 | 12691 | 11665 | 12178.0 |
| fluNS | 890 | 13581 | 12692 | 13136.5 |

Merge the *genotypes\_by\_gene* data frame with the sequence coordinates to create a new data frame (*genotypes\_by\_plot*) that gives the genotypic information in the coordinates used for the plotting below:

In [75]:

```
genotypes_to_plot <- genotypes_by_gene %>%
  filter(! grepl("missing", mutation_str_short)) %>%
  merge(flu_seq_coords, by="flu_gene") %>%
  transform(IFN_state=factor(IFN_state))
```

Get the coordinates of substitution mutations into a data frame (*substitution\_coords*) for use in the plotting below.
We also get the frequency of each mutation in the PacBio CCSs so we can use that to adjust the plotting size:

In [76]:

```
substitution_coords <- genotypes_complete %>%
  filter(barcoded_gene_present &
         (! is.na(mutations)) & 
         mutations != "WT" & 
         mutation_type == "substitutions") %>%
  mutate(mutations=strsplit(mutations, " ")) %>%
  unnest(mutations) %>%
  mutate(mutations=str_replace(mutations, paste0(flu_gene, '-'), '')) %>%
  tidyr::extract(mutations, 
                 c("site", "aa_subs", "freq_numerator", "freq_denominator"), 
                 '[ACGT](\\d+)[ACTG]_?(.*)_\\((\\d+)/(\\d+)\\)',
                 remove=FALSE) %>%
  mutate(aa_subs=aa_subs %>% 
    str_replace_all("flu", "") %>%
    str_replace_all("_", " ") %>%
    str_replace_all(c(Ala="A", Arg="R", Asn="N", Asp="D", Cys="C", Glu="E", Gln="Q",
                      Gly="G", His="H", Ile="I", Leu="L", Lys="K", Met="M", Phe="F",
                      Pro="P", Ser="S", Thr="T", Trp="W", Tyr="Y", Val="V"))
    ) %>%
  mutate(
    substitution_type=ifelse(nchar(aa_subs), "nonsynonymous", "synonymous"),
    frequency=as.numeric(freq_numerator) / as.numeric(freq_denominator)
    ) %>%
  merge(flu_seq_coords, by="flu_gene") %>%
  mutate(site=gene_start + as.integer(site) - 1) %>%
  select(cell, flu_gene, IFN_state, viral_barcode, n_viral_variants, site,
         aa_subs, substitution_type, gene_start, gene_end, frequency, mutations)
```

Get the coordinates of deletion mutations into a data frame (*deletion\_coords*) for use in the plotting below.
Importantly, deletions are shown at widths representing their actual size **unless they are very small**, because in that case they are two narrow to see. So very small deletions are "padded" which makes them no longer at actual size (they are slightly larger), but at least visible.

In [77]:

```
deletion_coords <- genotypes_complete %>%
  filter(barcoded_gene_present &
         (! is.na(mutations)) & 
         mutations != "WT" & 
         mutation_type == "deletions") %>%
  mutate(mutations=strsplit(mutations, " ")) %>%
  unnest(mutations) %>%
  mutate(mutations=str_replace(mutations, paste0(flu_gene, '-'), '')) %>%
  tidyr::extract(mutations,
                 c("del_start", "del_end", "freq_numerator", "freq_denominator"), 
                 'del(\\d+)to(\\d+)_\\((\\d+)/(\\d+)\\)',
                 remove=FALSE) %>%
  merge(flu_seq_coords, by="flu_gene") %>%
  mutate(del_start=as.integer(del_start) + gene_start - 1,
         del_end=as.integer(del_end) + gene_start - 1) %>%
  mutate(
    mutation_type='deletion',
    frequency=as.numeric(freq_numerator) / as.numeric(freq_denominator)
    ) %>%
  select(cell, flu_gene, viral_barcode, n_viral_variants, IFN_state,
         del_start, del_end, mutation_type, frequency, mutations)

# pad small deletions
min_width <- 15 # minimum width at which a deletion is shown
deletion_coords <- deletion_coords %>%
  mutate(del_len=del_end - del_start,
         del_padding=max(0, min_width - del_len),
         del_start=del_start - del_padding / 2,
         del_end=del_end + del_padding / 2)
```

The plotting is not currently set up to handle insertions since we do not have any in the current data set.
The next cell throws an error and stops the notebook if an insertion is present.
If we end up identifying an insertion in one of the genes, this cell and the plotting that follows it will need to be adjusted:

In [78]:

```
insertion_coords <- genotypes_complete %>% 
  filter(barcoded_gene_present &
         (! is.na(mutations)) & 
         mutations != "WT" & 
         mutation_type == "insertions") %>%
  mutate(mutations=strsplit(mutations, " ")) %>%
  unnest(mutations) %>%
  mutate(mutations=str_replace(mutations, paste0(flu_gene, '-'), '')) %>%
  tidyr::extract(mutations,
                 c("ins_site", "ins_len", "freq_numerator", "freq_denominator"), 
                 'ins(\\d+)len(\\d+)_\\((\\d+)/(\\d+)\\)',
                 remove=FALSE) %>%
  merge(flu_seq_coords, by="flu_gene") %>%
  mutate(ins_site=as.integer(ins_site) + gene_start - 1,
         ins_len=as.integer(ins_len)) %>%
  mutate(mutation_type='insertion',
         frequency=as.numeric(freq_numerator) / as.numeric(freq_denominator)
         ) %>%
  select(cell, flu_gene, viral_barcode, n_viral_variants, IFN_state,
         ins_site, ins_len, mutation_type, frequency)

# pad small insertions
min_width <- 15 # minimum width at which an insertion is shown
insertion_coords <- insertion_coords %>%
  mutate(ins_len=max(ins_len, min_width))
```

##### Plot genotypes¶

Now use gggenes to make a plot that visually displays the genotypes for all cells for which these are called.
We also indicate the amount of total mRNA in the cell that comes from virus and IFN.
This is a complex plot, so the code that creates it is fairly long.

First, write a function that plots the genotypes given a few arguments that will subsequently be customized:

In [79]:

```
arrow_height <- 0.3 # vertical space per arrow in inches
arrow_frac_height <- 0.55 # arrow takes up this much of available height
max_shape_size <- 4.6 # max size of substitutions, deletions, insertions
sub_size <- 0.55 # size of substitutions relative to `max_shape_size`
del_size <- 1 # size deletions relative to `max_shape_size`
ins_size <- del_size # size of segments for insertions

# Function to plot genotypes filtering for `col_name` %in% `col_val`.
# Expand y limits to make height for `ncells` cells. 
# Add title `title`.
# Returns named list with `percent_flu`, `percent_IFN`, `genotypes`
plot_genotypes <- function(col_name, col_val, ncells=NULL, title='') {
    
  # filter for cells of interest
  p_genotypes_to_plot <- genotypes_to_plot %>%
    filter(UQ(as.name(col_name)) %in% col_val)
  p_substitution_coords <- substitution_coords %>%
    filter(UQ(as.name(col_name)) %in% col_val)
  p_deletion_coords <- deletion_coords %>%
    filter(UQ(as.name(col_name)) %in% col_val)
  p_insertion_coords <- insertion_coords %>%
    filter(UQ(as.name(col_name)) %in% col_val)
    
  # get number of cells if not specified
  p_ncells <- p_genotypes_to_plot$cell %>% unique %>% length
  if (is.null(ncells))
    ncells <- p_ncells
    
  # make space for title
  if (nchar(title)) {
    ncells <- ncells + 2
    }
    
  # make plot of genotypes
  p_genes <- ggplot() +
    # plot arrows for genes
    geom_gene_arrow(data=p_genotypes_to_plot %>%
                         mutate(n_viral_variants=factor(
                           ifelse(n_viral_variants==1,
                           "one viral variant", "both viral variants"),
                           c("one viral variant", "both viral variants"))),
      aes(xmin=gene_start, xmax=gene_end, y=cell, fill=n_viral_variants), 
      arrow_body_height=unit(arrow_frac_height * arrow_height, 'in'), 
      arrowhead_height=unit(arrow_frac_height * arrow_height, 'in'),
      arrowhead_width=unit(0.3 * arrow_height, 'in'),
      size=0.4) +
    scale_fill_manual(values=rep(c(cbPalette[[3]], cbPalette[[6]]), 4),
      name="dual-barcode co-infection?") +
    
    # plot points for substitutions
    geom_point(data=p_substitution_coords %>%
        filter(n_viral_variants == 1),
      aes(x=site, y=cell, shape=substitution_type,
          size=frequency * sub_size),
      color=cbPalette[[7]], stroke=0.8) +
    geom_point(data=p_substitution_coords %>%
        filter(n_viral_variants == 2 & viral_barcode == "wt"),
      aes(x=site, y=cell, shape=substitution_type, 
          size=0.5 * frequency * sub_size),
      color=cbPalette[[7]], stroke=0.8,
      position=position_nudge(y=0.4 * arrow_height)) +
    geom_point(data=p_substitution_coords %>%
        filter(n_viral_variants == 2 & viral_barcode == "syn"),
      aes(x=site, y=cell, shape=substitution_type,
          size=0.5 * frequency * sub_size),
      color=cbPalette[[7]], stroke=1,
      position=position_nudge(y=-0.4 * arrow_height)) +
    scale_shape_manual(values=c(19, 1), name="point mutation") +
    scale_size(range=c(0, max_shape_size)) +
    guides(shape=guide_legend(override.aes=list(size=max_shape_size * sub_size)),
           size='none') +
    
    # plot lines for deletions
    geom_segment(data=p_deletion_coords %>% 
        filter(n_viral_variants == 1),
      aes(x=del_start, y=cell, xend=del_end, yend=cell,
          color=mutation_type, size=frequency * del_size)) +
    geom_segment(data=p_deletion_coords %>%
        filter(n_viral_variants == 2 & viral_barcode == "wt"),
      aes(x=del_start, y=cell, xend=del_end, yend=cell,
          color=mutation_type, size=0.38 * frequency * del_size),
      position=position_nudge(y=0.4 * arrow_height)) +
    geom_segment(data=p_deletion_coords %>%
        filter(n_viral_variants == 2 & viral_barcode == "syn"),
      aes(x=del_start, y=cell, xend=del_end, yend=cell,
          color=mutation_type, size=0.38 * frequency * del_size),
      position=position_nudge(y=-0.4 * arrow_height)) +
    scale_color_manual(values=c(cbPalette[[5]], cbPalette[[8]]),
                       name=NULL) +
    guides(color=guide_legend(
      override.aes=list(size=del_size * max_shape_size))) +
    
    # plot lines for insertions
    geom_segment(data=p_insertion_coords %>% 
        filter(n_viral_variants == 1),
      aes(x=ins_site, y=cell, xend=ins_site + ins_len, yend=cell,
          color=mutation_type, size=frequency * ins_size)) +
    geom_segment(data=p_insertion_coords %>%
        filter(n_viral_variants == 2 & viral_barcode == "wt"),
      aes(x=ins_site, y=cell, xend=ins_site + ins_len, yend=cell,
          color=mutation_type, size=0.38 * frequency * ins_size),
      position=position_nudge(y=0.4 * arrow_height)) +
    geom_segment(data=p_insertion_coords %>%
        filter(n_viral_variants == 2 & viral_barcode == "syn"),
      aes(x=ins_site, y=cell, xend=ins_site + ins_len, yend=cell,
          color=mutation_type, size=0.38 * frequency * ins_size),
      position=position_nudge(y=-0.4 * arrow_height)) +
    
    # label amino-acid substitutions
    geom_gene_label(data=p_substitution_coords %>% 
        group_by(cell, flu_gene, gene_start, gene_end) %>% 
        summarize(aa_subs=paste(aa_subs, collapse= " ")),
      aes(label=aa_subs, xmin=gene_start, xmax=gene_end, y=cell),
      size=6, min.size=2, reflow=TRUE, color='white',
      padding.y=unit(0.02 * arrow_height, "in")) +
    
    # other plot formatting
    theme(axis.ticks.y=element_blank(), axis.line.y=element_blank(),
      axis.text.y=element_blank(),
      panel.grid.major.y=ggplot2::element_line(colour = "grey", size = 1),
      plot.margin=unit(arrow_height * c(0, 1.2, 0.1, 0.4), "in")) +
    scale_x_continuous(expand=c(0.001, 0), breaks=flu_seq_coords$gene_center,
      labels=gsub("flu", "", flugenes)) +
    expand_limits(y=c(0, ncells)) +
    ylab(NULL) +
    xlab("viral gene") 
    
  # add title
  if (nchar(title))
    p_genes <- p_genes + annotate('text',
      x=(min(flu_seq_coords$gene_start) + max(flu_seq_coords$gene_end)) / 2,
      y=p_ncells + 1, label=title, size=6)
    
  # make plot of % flu
  p_flu <- ggplot(p_genotypes_to_plot %>% 
        mutate(x="flu", percent=frac_mRNA_from_flu * 100), 
      aes(x=x, y=cell, fill=percent, label=sprintf("%.0f", percent))) +
    geom_tile(color=cbPalette[[4]], height=arrow_frac_height, width=0.95, size=0.5) +
    geom_text(size=3) +
    scale_fill_gradient(low='white', high=cbPalette[[4]],
      limits=c(0, 100 * max(genotypes_to_plot$frac_mRNA_from_flu)),
      name="% mRNA from flu") +
    expand_limits(y=c(0, ncells)) +
    xlab("% mRNA") +
    theme(axis.title.x=element_text(hjust=0.2)) +
    ylab(NULL) +
    theme(axis.line.y=element_blank(),
      axis.ticks.y=element_blank(),
      plot.margin=unit(arrow_height * c(0, 0, 0, 0.3), "in"))
    
  # make plot of % IFN
  p_IFN <- ggplot(p_genotypes_to_plot %>% 
        mutate(x="IFN", percent=IFN_frac * 100), 
      aes(x=x, y=cell, fill=percent, color=IFN_state, label=sprintf("%.1f", percent))) +
    geom_tile(height=arrow_frac_height, width=0.95, size=0.5) +
    geom_text(size=3, color=cbbPalette[1]) +
    scale_fill_gradient(low='white', high=cbPalette[[2]],
      limits=c(0, 100 * max(genotypes_to_plot$IFN_frac)),
      name="% mRNA from IFN") +
    scale_color_manual(values=c(cbPalette[1], cbPalette[2]), drop=FALSE, guide=FALSE) +
    expand_limits(y=c(0, ncells)) +
    xlab(NULL) +
    ylab(NULL) +
    theme(axis.line.y=element_blank(),
      axis.text.y=element_blank(), axis.ticks.y=element_blank(),
      plot.margin=unit(arrow_height * c(0, 0, 0, 0), "in"))
    
  return(list(percent_flu=p_flu, percent_IFN=p_IFN, genotypes=p_genes))
  }
```

First make a plot that shows all cells wrapped into a specified number of columns:

In [80]:

```
p_blank <- ggplot(data.frame(c(1, 1))) + geom_blank()
```

In [81]:

```
ncols <- 2

cell_names <- genotypes_to_plot$cell %>% unique %>% sort(decreasing=TRUE)
ncells <- cell_names %>% length
ncells_per_col <- ceiling(ncells / ncols)

p_cells_list <- lapply(
  1:ncols,
  function (i) {
    col_val <- cell_names[((i - 1) * ncells_per_col + 1):
                          min(ncells, (i * ncells_per_col))]
    plot_genotypes('cell', col_val, ncells_per_col, '')
    }
  )

p_genotypes <- plot_grid(
  plotlist=
    append(lapply(1:ncols,
      function (i) plot_grid(
        plotlist=lapply(c('percent_flu', 'percent_IFN', 'genotypes'),
          function (p) p_cells_list[[i]][[p]] + theme(legend.position='none')
          ),
        align='h',
        rel_widths=c(0.12, 0.04, 1),
        nrow=1
        )
      ),
      list(plot_grid(
        p_blank,
        get_legend(p_cells_list[[1]][['percent_flu']]),
        get_legend(p_cells_list[[1]][['percent_IFN']]),
        get_legend(p_cells_list[[ncols]]['genotypes']),
        p_blank,
        cols=1,
        rel_heights=c(4, 1, 1, 1.5, 4)
        ))
    ),
  rel_widths=c(rep(1, ncols), 0.23),
  nrow=1
  )

saveShowPlot(p_genotypes,
             width=9 * ncols + 2,
             height=arrow_height * (2 + ncells_per_col),
             isfig=TRUE)
```

Now make a plot that splits into two columns, one for IFN+ and one for IFN- cells:

In [82]:

```
ncells_per_col_ifn <- max(
                genotypes_to_plot %>% filter(IFN_state == 'IFN+') %$%
                  cell %>% unique %>% length, 
                genotypes_to_plot %>% filter(IFN_state == 'IFN-') %$%
                  cell %>% unique %>% length)

p_ifn_pos <- plot_genotypes("IFN_state", c("IFN+"), ncells_per_col, "")
p_ifn_neg <- plot_genotypes("IFN_state", c("IFN-"), ncells_per_col, "")

p_genotypes_by_ifn <- plot_grid(
  plot_grid(
    p_ifn_neg[['percent_flu']] + theme(legend.position='none'),
    p_ifn_neg[['percent_IFN']] + theme(legend.position='none'),
    p_ifn_neg[['genotypes']] + theme(legend.position='none'),
    align='h',
    rel_widths=c(0.115, 0.039, 1),
    nrow=1
    ),
  plot_grid(
    p_ifn_pos[['percent_flu']] + theme(legend.position='none'),
    p_ifn_pos[['percent_IFN']] + theme(legend.position='none'),
    p_ifn_pos[['genotypes']] + theme(legend.position='none'),
    align='h',
    rel_widths=c(0.115, 0.039, 1),
    nrow=1
    ),
  get_legend(p_ifn_neg[['genotypes']]),
  rel_widths=c(1, 1, 0.23),
  nrow=1
  )

saveShowPlot(p_genotypes_by_ifn, width=20,
  height=arrow_height * (3 + ncells_per_col),
  isfig=TRUE)
```

The plots above summarize essentially all of the genotypic data.
They look better if you open them and view at higher resolution.

##### Print genotypes¶

We also have a table that gives the same information displayed in the plot above.
However, it also has a few pieces of additional relevant data: each mutation is suffixed by the number of PacBio CCSs in which it is observed, and the exact identities of all mutations are shown.
Below we show the first few lines, and then write the table to a file for the paper:

In [83]:

```
genotype_table <- genotypes_by_gene %>%
  arrange(flu_gene) %>%
  group_by(cell, cell_barcode, frac_mRNA_from_flu, IFN_frac, IFN_state, coinfection_state) %>%
  summarize(mutations=paste0(trimws(mutation_str_long), collapse="; "),
         mutations=gsub('(; )+', '; ', mutations),
         mutations=gsub('; $', '', mutations),
         mutations=gsub('^; ', '', mutations)) %>%
  arrange(desc(cell))

genotype_table %>% head

genotype_table %>% write.csv(file.path(figsdir, 'genotypes.csv'), row.names=FALSE)
```

| cell | cell\_barcode | frac\_mRNA\_from\_flu | IFN\_frac | IFN\_state | coinfection\_state | mutations |
| --- | --- | --- | --- | --- | --- | --- |
| cell 1 | CAAGAAATCGTGACAT | 0.6511267 | 1.140186e-04 | IFN- | single viral barcode |  |
| cell 2 | TGTGTTTTCAACGGGA | 0.6234279 | 4.164238e-05 | IFN- | dual-barcode coinfection | fluPA-WT / fluPA-del207to1867\_(8874/8951); fluHA-WT / fluHA-A1338G\_fluHA-Thr436Ala\_(24/46) |
| cell 3 | TTCTACAAGCCCAATT | 0.6095883 | 7.283321e-05 | IFN- | dual-barcode coinfection | fluPB1-del427to1876\_(1083/1089) / fluPB1-WT |
| cell 4 | CCTACACCAGATAATG | 0.5901598 | 2.178649e-03 | IFN+ | single viral barcode | fluNA-missing; fluM-missing |
| cell 5 | TTATGCTAGCTACCTA | 0.5657166 | 5.889282e-04 | IFN+ | dual-barcode coinfection | fluPB2-WT / fluPB2-T1143G\_(16/48); fluPB1-del385to2163\_(3353/3378) / fluPB1-WT; fluNA-WT / fluNA-A836G\_fluNA-Met273Val\_(15/27) |
| cell 6 | GACCAATGTGGACGAT | 0.5653513 | 1.324416e-04 | IFN- | single viral barcode | fluPB1-G1467A\_(14/33); fluHA-A1375G\_fluHA-Glu448Gly\_(13/40); fluNA-A379C\_fluNA-Gln120His\_(27/46) |

Even more extensive information (for instance, is the *wt* or *syn* viral barcode variant present in each cell) is in the full *genotypes\_by\_gene* data frame.
That is too long to print in this notebook, but the cell below shows the first few lines as an illustration:

In [84]:

```
genotypes_by_gene %>% 
  select(-n_viral_variants, -mutation_str_short) %>% 
  arrange(desc(cell)) %>% 
  head
```

| cell | cell\_barcode | frac\_mRNA\_from\_flu | IFN\_frac | IFN\_state | coinfection\_state | flu\_gene | syn | wt | mutation\_str\_long |
| --- | --- | --- | --- | --- | --- | --- | --- | --- | --- |
| cell 1 | CAAGAAATCGTGACAT | 0.6511267 | 0.0001140186 | IFN- | single viral barcode | fluHA | fluHA-WT | fluHA-missing |  |
| cell 1 | CAAGAAATCGTGACAT | 0.6511267 | 0.0001140186 | IFN- | single viral barcode | fluM | fluM-WT | fluM-missing |  |
| cell 1 | CAAGAAATCGTGACAT | 0.6511267 | 0.0001140186 | IFN- | single viral barcode | fluNA | fluNA-WT | fluNA-missing |  |
| cell 1 | CAAGAAATCGTGACAT | 0.6511267 | 0.0001140186 | IFN- | single viral barcode | fluNP | fluNP-WT | fluNP-missing |  |
| cell 1 | CAAGAAATCGTGACAT | 0.6511267 | 0.0001140186 | IFN- | single viral barcode | fluNS | fluNS-WT | fluNS-missing |  |
| cell 1 | CAAGAAATCGTGACAT | 0.6511267 | 0.0001140186 | IFN- | single viral barcode | fluPA | fluPA-WT | fluPA-missing |  |

Finally, although we think the two viral barcodes are equivalent, we just confirm that among the IFN+ cells infected with wildtype virus, we don't see any bias towards one viral barcode.
The analysis below confirms that there is no such bias: among IFN+ cells infected with wildtype, 3 have the wildtype viral variant, 4 have the synonymous viral variant, and 2 are co-infected:

In [85]:

```
genotypes_by_gene %>% 
  mutate(has_syn=! str_detect(syn, "missing"),
         has_wt=! str_detect(wt, "missing")) %>%
  group_by(cell, IFN_state) %>%
  summarize(
    has_syn=any(has_syn),
    has_wt=any(has_wt),
    mutations=paste0(trimws(mutation_str_long), collapse=""),
    is_wildtype=('' == mutations)
    ) %>%
  group_by(IFN_state, has_syn, has_wt, is_wildtype) %>%
  summarize(ncells=n()) %>%
  arrange(IFN_state, is_wildtype, has_syn, has_wt)
```

| IFN\_state | has\_syn | has\_wt | is\_wildtype | ncells |
| --- | --- | --- | --- | --- |
| IFN- | FALSE | TRUE | FALSE | 16 |
| IFN- | TRUE | FALSE | FALSE | 38 |
| IFN- | TRUE | TRUE | FALSE | 22 |
| IFN- | FALSE | TRUE | TRUE | 1 |
| IFN- | TRUE | FALSE | TRUE | 23 |
| IFN- | TRUE | TRUE | TRUE | 10 |
| IFN+ | FALSE | TRUE | FALSE | 6 |
| IFN+ | TRUE | FALSE | FALSE | 18 |
| IFN+ | TRUE | TRUE | FALSE | 7 |
| IFN+ | FALSE | TRUE | TRUE | 3 |
| IFN+ | TRUE | FALSE | TRUE | 4 |
| IFN+ | TRUE | TRUE | TRUE | 2 |

#### Viral features that correlate with IFN induction¶

Examine the genetic features of viruses (gene absence, mutations) that associated with IFN expression.

##### Data frames with relevant data¶

Get a data frame (*infection\_data*) that has all information of potential interest about each infected cell, and then has a row for each flu gene and mutation type (amino-acid substitutions, insertions, deletions, gene absence) that has a column *has\_mutation* that is:

- *NA*: if sequence for **any** viral barcode present in that cell is unknown in a way that prevents us from calling mutation
- *TRUE*: if mutation is present. For gene absence, both barcodes must be absent. For other mutations, either can be mutated.
- *FALSE*: if mutation is not present.

In [86]:

```
# data frame annotated with all mutation types
infection_data <- genotypes %>%
  # get call barcode without suffix
  separate(CellBarcode, "cell_barcode", extra="drop") %>%
  # call mutation status, disregard synonymous mutations
  tidyr::extract(mutations, c("aa_subs"), paste0('[ACGT]\\d+[ACTG]_',
    '([:alnum:]+\\-+[:upper:][:lower:]{2}\\d+[:upper:][:lower:]{2})_\\(\\d+/\\d+\\)'),
    remove=FALSE) %>%
  mutate(has_mutation=
    ifelse(! barcoded_gene_present, FALSE,
    ifelse(is.na(mutations), NA,
    ifelse(mutations == "WT", FALSE,
    ifelse(mutation_type != "substitutions", TRUE,
    ifelse(mutation_type == "substitutions" & is.na(aa_subs), FALSE, TRUE)))))) %>%
  # get columns of interest
  select(cell_barcode, total_mRNAs, frac_mRNA_from_flu,
         sapply(flugenes, function (g) paste0(g, "_frac")) %>% as.vector,
         sapply(flugenes, function (g) paste0("has_", g)) %>% as.vector,
         n_flu_genes, ISG_frac, ISG_state, IFN_frac, IFN_state,
         coinfection_state, complete_state, mutation_type, flu_gene,
         viral_barcode, has_mutation, barcoded_gene_present) %>%
  # aggregate data for both viral barcodes into gene_status
  group_by(cell_barcode, flu_gene, mutation_type) %>%
  mutate(has_mutation=
    ifelse(any(is.na(has_mutation)), NA,
    ifelse(any(has_mutation), TRUE, FALSE))) %>%
  # gene presence / absence aggregated across viral barcodes
  group_by(cell_barcode, flu_gene) %>%
  mutate(gene_present=any(barcoded_gene_present)) %>%
  select(-barcoded_gene_present, -viral_barcode) %>%
  unique %>%
  ungroup 
                
# add gene absence as a mutation type
infection_data <- rbind(
  infection_data,
  infection_data %>% 
    mutate(mutation_type="gene absent",
           has_mutation=!gene_present) %>%
    unique
  )

# indicate cells that are infected by fully wildtype virus
infection_data <- infection_data %>%
  group_by(cell_barcode) %>%
  mutate(wildtype_virus=ifelse(any(is.na(has_mutation)) | any(has_mutation),
    "incomplete / mutated", "full wildtype")) %>%
  ungroup
```

Write data frame to a file for paper.
This data frame has information on mutation presence with a row for each gene and mutation type:

In [87]:

```
infection_data %>% head

infection_data %>% write.csv(file.path(figsdir, 'mutations.csv'), row.names=FALSE)
```

| cell\_barcode | total\_mRNAs | frac\_mRNA\_from\_flu | fluPB2\_frac | fluPB1\_frac | fluPA\_frac | fluHA\_frac | fluNP\_frac | fluNA\_frac | fluM\_frac | ⋯ | ISG\_state | IFN\_frac | IFN\_state | coinfection\_state | complete\_state | mutation\_type | flu\_gene | has\_mutation | gene\_present | wildtype\_virus |
| --- | --- | --- | --- | --- | --- | --- | --- | --- | --- | --- | --- | --- | --- | --- | --- | --- | --- | --- | --- | --- |
| AAAGTAGTCGTGGACC | 54025 | 0.12807034 | 0.013152190 | 0.011562365 | 0.007226478 | 0.10203787 | 0.1524787 | 0.18933372 | 0.04321434 | ⋯ | ISG+ | 2.122872e-05 | IFN- | dual-barcode coinfection | complete | substitutions | fluPB2 | FALSE | TRUE | incomplete / mutated |
| AACACGTGTCTAGTGT | 24428 | 0.13689209 | 0.074461722 | 0.007177033 | 0.040370813 | 0.07505981 | 0.3666268 | 0.20155502 | 0.20574163 | ⋯ | ISG+ | 2.798330e-03 | IFN+ | single viral barcode | complete | substitutions | fluPB2 | FALSE | TRUE | incomplete / mutated |
| AACGTTGTCATCACCC | 31015 | 0.17020796 | 0.006061754 | 0.008145482 | 0.007956052 | 0.04641030 | 0.1812843 | 0.09812464 | 0.30327714 | ⋯ | ISG- | 7.771215e-05 | IFN- | dual-barcode coinfection | incomplete | substitutions | fluPB2 | NA | TRUE | incomplete / mutated |
| AACTCAGAGATAGCAT | 21021 | 0.02668760 | 0.023172906 | 0.026737968 | 0.021390374 | 0.03743316 | 0.5793226 | 0.09269162 | 0.02495544 | ⋯ | ISG+ | 1.559140e-02 | IFN+ | single viral barcode | incomplete | substitutions | fluPB2 | NA | TRUE | incomplete / mutated |
| AACTCTTCACGGATAG | 46934 | 0.01717305 | 0.044665012 | 0.029776675 | 0.002481390 | 0.12779156 | 0.1848635 | 0.17990074 | 0.16004963 | ⋯ | ISG- | 2.167881e-05 | IFN- | dual-barcode coinfection | incomplete | substitutions | fluPB2 | FALSE | TRUE | incomplete / mutated |
| AACTCTTGTCTAAACC | 20095 | 0.39995024 | 0.004728132 | 0.022396417 | 0.015553067 | 0.11646137 | 0.1383601 | 0.09655344 | 0.22919000 | ⋯ | ISG- | 8.293249e-05 | IFN- | dual-barcode coinfection | complete | substitutions | fluPB2 | FALSE | TRUE | incomplete / mutated |

Then get a smaller data frame (*cell\_infection\_data*) that just has the cell-specific data in *infection\_data* with one cell per row:

In [88]:

```
cell_infection_data <- infection_data %>%
  select(-mutation_type, -flu_gene, -has_mutation, -gene_present) %>%
  unique
```

##### Viral genetics and flu burden heterogeneity¶

We've already shown that expression of flu mRNA is highly heterogeneous across infected cells.
Is part of the heterogeneity due to viral genetics?
To assess this, we plot the distribution of viral mRNA among cells for which we obtained complete viral genomes, faceting by whether those cells are infected by fully wildtype virus.
We quantify the heterogeneity in flu mRNA production by the Gini coefficient, which we here refer to using the alternative nomenclature of *Gini index* since it is shorter and fits on the plot better.
Note that these plots are already conditioned on us obtaining a full genome of the infecting virus (which preferentially happens for cells with lots of viral mRNA), so this already reduced the heterogeneity below that observed in Russell et al (2018).

Nonetheless, conditioning on whether we have a full viral genome, the plots below shows that cells infected by wildtype viruses have less heterogeneity than cells infected by viruses with mutations / indel, indicating that viral genetics contributes to the heterogeneity.
The plots show the 95% confidence intervals for the Gini coefficients:

In [89]:

```
# data frame with complete infections and Gini coefficients on frac flu
frac_flu_df <- cell_infection_data %>%
  filter(complete_state == "complete") %>%
  group_by(wildtype_virus) %>%
  mutate(gini=Gini(frac_mRNA_from_flu, conf.level=0.95) %>% 
    (function (x) sprintf("%.2f (%.2f, %.2f)", x[1], x[2], x[3]))) %>%
  ungroup

# make histogram
p_frac_flu_wt_hist <- ggplot(
    frac_flu_df,
    aes(frac_mRNA_from_flu)) +
  geom_histogram(bins=20, fill=cbPalette[3]) +
  facet_wrap(~ wildtype_virus, ncol=1) +
  xlab("frac mRNA from flu") +
  scale_y_continuous(name="number of cells", breaks=pretty_breaks(4)) +
  expand_limits(y=c(0, 14)) +
  theme(strip.text=element_text(size=14, margin=margin(b=3)))

# annotate with Gini coefficients
p_frac_flu_wt <- ggdraw() +
  draw_plot(p_frac_flu_wt_hist, 0, 0, 1, 1) +
  draw_label(
    paste('Gini index',
          frac_flu_df %>%
            filter(wildtype_virus != "full wildtype") %>%
            head(n=1) %$%
            gini),
    x=0.97, y=0.47, size=12, hjust=1, fontface='italic') +
  draw_label(
    paste('Gini index',
          frac_flu_df %>%
            filter(wildtype_virus == "full wildtype") %>%
            head(n=1) %$%
            gini),
    x=.97, y=0.89, size=12, hjust=1, fontface='italic')
     
saveShowPlot(p_frac_flu_wt, 3, 3.8)
```

##### Viral genetics and IFN induction¶

We now look to see if IFN induction is less common in cells infected with wildtype virus.
We will then do a similar analysis with other variables, such as ISG levels and coinfection.
So we create a general function that does the analysis by making an annotated histogram, generating a contingency table, and performing a Fisher's exact test.

In [90]:

```
# Plot faceted histogram for viruses with complete sequenced genomes,
# with numerical annotations on histogram.
# Returns list of plot, contingeny table, and Fisher's exact test result.
# Designed to show how IFN or ISG levels depend on a faceting variable.
# `ifn_or_isg`: plot IFN or ISG levels on x-axis
# `facet_var`: variable to facet on, `wildtype_virus` or `coinfection_state`
plot_hist <- function (ifn_or_isg, facet_var) {
  
  df <- cell_infection_data %>%
    filter(complete_state == "complete") %>%
    mutate(frac=if (ifn_or_isg == "IFN") IFN_frac else ISG_frac,
           state=if (ifn_or_isg == "IFN") IFN_state else ISG_state,
           facet_var=if (facet_var == "wildtype_virus") wildtype_virus
                     else coinfection_state)
               
  p_hist <- ggplot(
      df,
      aes(frac, fill=state)) +
    geom_histogram(bins=42) +
    facet_wrap(~ facet_var, ncol=1) +
    xlab(paste("frac mRNA from", ifn_or_isg)) +
    scale_y_continuous(name="number of cells", breaks=pretty_breaks(4)) +
    scale_fill_manual(
      values=if (ifn_or_isg == "IFN") rev(cbPalette[2:3]) else
                    c(cbPalette[3], cbPalette[7]),
      name=NULL) +
    theme(strip.text=element_text(size=14, margin=margin(b=3)),
          legend.position=c(0.65, 0.89))
           
  facet_vals <- df %>% arrange(facet_var) %$% facet_var %>% unique

  # annotate with fractions
  p <- ggdraw() +
    draw_plot(p_hist, 0, 0, 1, 1) +
    draw_label(
      paste(df %>%
              filter(facet_var != facet_vals[[1]] & 
                     state == paste0(ifn_or_isg, "+")) %>%
              nrow,
            "of",
            df %>%
              filter(facet_var != facet_vals[[1]]) %>%
              nrow,
            paste0(ifn_or_isg, "+")),
      x=0.3, y=0.47, size=12, hjust=0, fontface='italic') +
    draw_label(
      paste(df %>%
              filter(facet_var == facet_vals[[1]] & 
                     state == paste0(ifn_or_isg, "+")) %>%
              nrow,
            "of",
            df %>%
              filter(facet_var == facet_vals[[1]]) %>%
              nrow,
            paste0(ifn_or_isg, "+")),
      x=0.3, y=0.89, size=12, hjust=0, fontface='italic')
           
  contingency_table <- df %>% 
    group_by(facet_var, state) %>%
    summarize(n=n()) %>%
    spread(key=facet_var, value=n)

  test <- fisher.test(contingency_table %>% select(-state))
         
  return (list(p, contingency_table, test))
  }
```

Now we look at the association of a cell being infected with full wildtype virus and being IFN+:

In [ ]:

```
# perform analysis
p_frac_ifn_wt_list <- plot_hist(ifn_or_isg="IFN", facet_var="wildtype_virus")

# get and show histogram
p_frac_ifn_wt <- p_frac_ifn_wt_list[[1]]
saveShowPlot(p_frac_ifn_wt, 3, 3.8)

# show contingency table and Fisher's exact test result
p_frac_ifn_wt_list[[2]]
p_frac_ifn_wt_list[[3]]
```

We see that cells infected with wildtype virus are less likely to be IFN+, and that when they are they tend to express less IFN.

However, the difference in the number of IFN+ cells among those infected with wildtype virus is **not** statistically significant, probably because the numbers are so small.

##### Known co-infection and IFN induction¶

We next repeat the above analysis to see if known dual-barcode coinfections are associated with IFN induction.
The plots below show that known coinfected cells are just about as likely to be IFN+ as cells that are infected with just one viral barcode.
Again, the caveat to this is that we only identify about half of the coinfections.

In [ ]:

```
# perform analysis
p_frac_ifn_coinfect_list <- plot_hist(ifn_or_isg="IFN", facet_var="coinfection_state")

# get and show histogram
p_frac_ifn_coinfect <- p_frac_ifn_coinfect_list[[1]]
saveShowPlot(p_frac_ifn_coinfect, 3, 3.8)

# show contingency table and Fisher's exact test result
p_frac_ifn_coinfect_list[[2]]
p_frac_ifn_coinfect_list[[3]]
```

##### Gene absence / mutations and IFN induction¶

Now we check to see if the absence of any genes or mutations in any genes are associated with IFN induction.
We use a Fisher's exact test to see if any genes are absent more in the IFN+ cells than the IFN- cells, and then use an FDR correction for multiple hypotheses.

For this analysis, we look at:

- complete absence of genes
- amino-acid substitutions (ignoring synonymous point mutations)
- deletions
  We don't analyze insertions, because the genotypes plot above already makes clear that insertions are extremely uncommon in our data set.

For each type of mutation, we calculate the percentage of cells that have that mutation in their gene in both the IFN+ and IFN- condition.
For substitutions and deletions, we condition this analysis on the cell having the gene (i.e., we exclude from the calculations cells that are completely lacking the gene).

We also use Fisher's exact test to calculate the P-value that there is a difference in the frequency of the mutation among the IFN+ and IFN- cells.

First, we define a function that makes these plots.
We generalize this function so that it can show varying levels of annotations and work for IFN or ISGs:

In [ ]:

```
# function to plot association of mutations with IFN or ISG
# `complete_only`: only show cells with complete genomes sequenced
# `show_counts`: show numerical counts on plot
# `showQvalue`: show Q-values in addition to P-values
# `ifn_or_isg`: perform plotting for IFN or ISG?
plot_muts <- function (complete_only, show_counts, showQvalue, ifn_or_isg) {
    
  mutation_data <- infection_data %>%
    filter(mutation_type != "insertions") %>%
    transform(
      mutation_type=factor(mutation_type, c("gene absent", "deletions", "substitutions")),
      flu_gene=factor(gsub("flu", "", flu_gene), flugenes_noprefix)
      ) %>%
    filter(gene_present | mutation_type == "gene absent") %>%
    filter(complete_state == "complete" | ! complete_only) %>%
    filter(! is.na(has_mutation)) %>%
    mutate(state=if (ifn_or_isg == "IFN") IFN_state else ISG_state) %>%
    group_by(mutation_type, flu_gene) %>%
    summarize(
      pos_mutated=sum(has_mutation & state == paste0(ifn_or_isg, "+")),
      neg_mutated=sum(has_mutation & state == paste0(ifn_or_isg, "-")),
      pos=sum(state == paste0(ifn_or_isg, "+")),
      neg=sum(state == paste0(ifn_or_isg, "-")),
      pos_percent=100 * pos_mutated / pos,
      neg_percent=100 * neg_mutated / neg,
      P=fisher.test(
        matrix(c(pos_mutated, pos - pos_mutated,
                 neg_mutated, neg - neg_mutated),
               nrow=2),
        )[["p.value"]],
      ) %>%
    ungroup %>%
    gather("state", "percent_mutated", pos_percent, neg_percent) %>%
    mutate(state=str_replace(state, "pos_percent", "+"),
           state=str_replace(state, "neg_percent", "-"),
           denom=ifelse(state == "+", pos, neg),
           num=ifelse(state == "+", pos_mutated, neg_mutated),
           label=sprintf("%d /\n%d", num, denom),
           Q=qvalue(P)$qvalue,
           Plabel=if (showQvalue)
             paste0(signif(P, 1), " / ", signif(Q, 1))
             else signif(P, 1)
           )

  ymax <- mutation_data$percent_mutated %>% max
           
  if (showQvalue)
    label_size <- 3
  else
    label_size <- 3.5

  p <- ggplot(
      mutation_data,
      aes(state, percent_mutated, fill=state)) +
    geom_bar(stat='identity') +
    geom_text(aes(label=Plabel), 
      y=1.09 * ymax, x=1.5, fontface="italic", vjust=0, size=label_size) +
    geom_segment(x=1, xend=2, y=1.05 * ymax, yend=1.05 * ymax, size=0.25) +
    facet_grid(mutation_type ~ flu_gene) +
    scale_fill_manual(values=
      if (ifn_or_isg == "IFN") rev(cbPalette[2:3]) else 
          c(cbPalette[3], cbPalette[7])) +
    theme(
      legend.position='none',
      strip.text=element_text(size=14, margin=margin(l=4, r=1)),
      axis.text.x=element_text(size=14)
      ) +
    expand_limits(y=c(0, 1.2 * ymax)) +
    xlab(paste(ifn_or_isg, "state")) +
    ylab("percent of cells with mutation")
    
  if (show_counts)
    p <- p + geom_text(aes(label=label),
      y=0, vjust=-0.1, hjust=0.5, fontface="italic", size=3)
    
  return (p)
  }
```

Now make a plot showing just the cells for which we could completely sequence the infection virions.
This plot shows that absence of NS is strongly associated with IFN induction, that amino-acid substitutions in PB1 are modestly corelated with IFN+, and that deletions in HA and substitutions in NS are perhaps weakly associated.
The numbers of above the plots show the P value and the Q value.
Only the NS absence association remains significant at an FDR of 0.1.

In [ ]:

```
p_muts_ifn_complete <- plot_muts(
  complete_only=TRUE, show_counts=FALSE, 
  showQvalue=FALSE, ifn_or_isg="IFN")

saveShowPlot(p_muts_ifn_complete, width=7, height=4.75)
```

Now make a version of the same plot where we no longer require the genomes of the infecting virions to be completely sequenced, but also include genes from partially sequenced virions.
This will include more cells with low viral burden, as those are the ones for which we often don't sequence the full infecting virus.

The trends here remain largely similar to above, although there is now a borderline trend for IFN+ to be depleted in absence (enriched in presence) of PB2 and PA:

In [ ]:

```
p_muts_ifn_all <- plot_muts(
  complete_only=FALSE, show_counts=FALSE,
  showQvalue=FALSE, ifn_or_isg="IFN")

saveShowPlot(p_muts_ifn_all, width=7, height=4.75, isfig=TRUE)
```

##### Amount of viral mRNA and IFN induction¶

Next we check if the amount of viral mRNA is associated with IFN induction.
Since lack of NS is a major determinant of IFN positivity, we do this faceting by NS presence.
We see that among NS-expressing infected cells there is no association with the amount of flu and IFN induction.
But among NS-deficient cells, there is a significant assocation between high amounts of viral mRNA and IFN induction.

In [ ]:

```
ymax <- cell_infection_data$frac_mRNA_from_flu %>% max

p_viral_load_ifn <- ggplot(
    cell_infection_data %>%
      mutate(NS_state=ifelse(has_fluNS, "has NS", "lacks NS"),
             IFN_state=gsub("IFN", "", IFN_state)),
    aes(IFN_state, frac_mRNA_from_flu, fill=IFN_state)) +
  geom_boxplot(outlier.size=1, outlier.alpha=0.4) +
  stat_compare_means(
    aes(label=..p.format..),
    method="wilcox.test", hjust=0.5, vjust=0, label.x=1.5,
    label.y=1.08 * ymax, fontface="italic"
    ) +
  geom_segment(x=1, xend=2, y=1.04 * ymax, yend=1.04 * ymax, size=0.25) +
  scale_fill_manual(values=rev(cbPalette[2:3])) +
  theme(legend.position="none",
        strip.text=element_text(size=14),
        axis.text.x=element_text(size=14)) +
  facet_wrap(~ NS_state, ncol=1) +
  expand_limits(y=c(0, 1.17 * ymax)) +
  scale_y_continuous(name="frac mRNA from flu") +
  xlab("IFN state")

saveShowPlot(p_viral_load_ifn, width=2, height=3.8)
```

#### Viral features the correlate with ISG expression¶

Now we parallel the analysis in the preceding section, but look for features that associate with ISG expression rather than IFN induction.

##### Viral genetics and ISG expression¶

We see that similarly for IFN induction, ISG expression is more common and higher in cells that fail to express wildtype copies of all viral genes.
The difference here is statisticall significant:

In [ ]:

```
# perform analysis
p_frac_isg_wt_list <- plot_hist(ifn_or_isg="ISG", facet_var="wildtype_virus")

# get and show histogram
p_frac_isg_wt <- p_frac_isg_wt_list[[1]]
saveShowPlot(p_frac_isg_wt, 3, 3.8)

# show contingency table and Fisher's exact test result
p_frac_isg_wt_list[[2]]
p_frac_isg_wt_list[[3]]
```

##### Known co-infection and ISG expression¶

As for IFN induction, there is no substantial association between known co-infections and ISG expression:

In [ ]:

```
# perform analysis
p_frac_isg_coinfect_list <- plot_hist(ifn_or_isg="ISG", facet_var="coinfection_state")

# get and show histogram
p_frac_isg_coinfect <- p_frac_isg_coinfect_list[[1]]
saveShowPlot(p_frac_isg_coinfect, 3, 3.8)

# show contingency table and Fisher's exact test result
p_frac_isg_coinfect_list[[2]]
p_frac_isg_coinfect_list[[3]]
```

##### Viral genetics and ISG expression¶

Most of the viral features that were associated with IFN induction are also associated with ISG expression, although on the absence of NS is significantly associated:

In [ ]:

```
p_muts_isg_complete <- plot_muts(
  complete_only=TRUE, show_counts=FALSE, 
  showQvalue=FALSE, ifn_or_isg="ISG")

saveShowPlot(p_muts_isg_complete, width=7, height=4.75)
```

##### Amount of viral mRNA and ISG expression¶

Similar to above for IFN, cells that have NS exhibit no association between the amount of viral mRNA and ISG expression.
But for cells that lack NS, there is a strong tendency for high-flu cells to have more ISG expression:

In [ ]:

```
ymax <- cell_infection_data$frac_mRNA_from_flu %>% max

p_viral_load_isg <- ggplot(
    cell_infection_data %>%
      mutate(NS_state=ifelse(has_fluNS, "has NS", "lacks NS"),
             ISG_state=gsub("ISG", "", ISG_state)),
    aes(ISG_state, frac_mRNA_from_flu, fill=ISG_state)) +
  geom_boxplot(outlier.size=1, outlier.alpha=0.4) +
  stat_compare_means(
    aes(label=..p.format..),
    method="wilcox.test", hjust=0.5, vjust=0, label.x=1.5,
    label.y=1.08 * ymax, fontface="italic"
    ) +
  geom_segment(x=1, xend=2, y=1.04 * ymax, yend=1.04 * ymax, size=0.25) +
  scale_fill_manual(values=c(cbPalette[3], cbPalette[7])) +
  theme(legend.position="none",
        strip.text=element_text(size=14),
        axis.text.x=element_text(size=14)) +
  facet_wrap(~ NS_state, ncol=1) +
  expand_limits(y=c(0, 1.17 * ymax)) +
  scale_y_continuous(name="frac mRNA from flu") +
  xlab("ISG state")

saveShowPlot(p_viral_load_isg, width=2, height=3.8)
```

#### Figures for paper¶

We have made all of the plots above, and saved some of them to the figures directory already by using the *isfig=TRUE* argument to `saveShowPlot`.
However, there are others that we want to assemble into multi-panel figures.
We do that here.

First, we assemble a figure that shows the calling of cells, infected cells, and IFN+ cells:

In [ ]:

```
p_cell_summary <- plot_grid(
  plot_grid(p_cellcounts, p_flu_vs_cell,
            ncol=1, rel_heights=c(1, 1.5), scale=0.9,
            labels=c("A", "B"), label_size=18, vjust=1),
  plot_grid(p_frac_flu, labels="C", scale=0.95, label_size=18, vjust=1),
  plot_grid(p_nflu_genes, labels="D", scale=0.95, label_size=18, vjust=1),
  plot_grid(p_flu_rel_expr, p_coinfect,
            scale=0.95, ncol=1,
            labels=c("E", "F"), label_size=18, vjust=1),
  plot_grid(p_ifn_dist, labels="G", scale=0.95, label_size=18, vjust=1),
  nrow=1, scale=0.95, rel_widths=c(1, 0.7, 0.6, 0.75, 0.7), align='h'
  ) + 
  theme(plot.margin=unit(c(t=0, r=0, b=-0.3, l=0), "in"))

saveShowPlot(p_cell_summary, width=15.5, height=4.9, isfig=TRUE)
```

Now a supplementary figure to the one above with the single-cell transcriptomic data:

In [ ]:

```
p_cell_summary_supp <- plot_grid(
  p_frac_has_gene,
  p_ifn_genes_corr,
  p_isg_dist,
  p_isg_corr,
  ncol=2,
  scale=0.9,
  rel_heights=c(0.68, 1),
  labels=c("A", "B", "C", "D"), label_size=18, vjust=2, hjust=-1
  )

saveShowPlot(p_cell_summary_supp, width=9.5, height=7.5, isfig=TRUE)
```

Next we assemble a figure that shows how viral mutations are associated with heterogeneity and IFN induction:

In [ ]:

```
p_mutations <- plot_grid(
  p_frac_flu_wt,
  p_frac_ifn_wt,
  p_muts_ifn_complete,
  p_viral_load_ifn,
  labels=c("A", "B", "C", "D"), label_size=18, vjust=1.2, hjust=-1,
  nrow=1, scale=0.93, rel_widths=c(1, 1, 2.4, 0.6), align='h'
  ) + 
  theme(plot.margin=unit(c(t=0, r=0, b=-0.3, l=0), "in"))

saveShowPlot(p_mutations, width=15.5, height=4.9, isfig=TRUE)
```

Next assemble a merged figure supplement that shows how viral mutations are associated with ISG expression:

In [ ]:

```
p_mutations_isg <- plot_grid(
  p_frac_isg_wt,
  p_muts_isg_complete,
  p_viral_load_isg,
  labels=c("A", "B", "C"), label_size=18, vjust=1.2, hjust=-1,
  nrow=1, scale=0.93, rel_widths=c(1, 2.4, 0.6), align='h'
  ) + 
  theme(plot.margin=unit(c(t=0, r=0, b=-0.3, l=0), "in"))

saveShowPlot(p_mutations_isg, width=12.5, height=4.9, isfig=TRUE)
```
